# Analysis of the heterogeneity and complexity of murine extraorbital lacrimal gland via single-cell RNA sequencing

**DOI:** 10.1101/2023.05.10.540166

**Authors:** Sen Zou, Xinwei Jiao, Shenzhen Huang, Jiangman Liu, Hongli Si, Di Qi, Xiaoting Pei, Dingli Lu, Yimian Wang, Zhijie Li

## Abstract

**Purpose:** The lacrimal gland is essential for maintaining ocular surface health and avoiding external damage by secreting the aqueous layer of the tear film. However, a healthy lacrimal gland’s inventory of cell types and heterogeneity remains understudied.

**Methods:** Here, 10X genome-based single-cell RNA sequencing was used to generate an unbiased classification of cellular diversity in the extraorbital lacrimal gland (ELG) of C57BL/6J mice. From 48,077 high-quality cells, an atlas of cell heterogeneity was produced, and cell types were defined by classic marker genes. The possible functions of these cells and the pseudotime trajectories for certain cell populations were analyzed through bioinformatics. In addition, a preliminary analysis of the cell-cell communication network in ELG was performed.

**Results:** Over 41 subclasses of cells were identified, including 9 kinds of glandular epithelial cells, 7 kinds of fibroblasts, 10 kinds of myeloid-derived immune cells, at least 10 kinds of lymphoid-derived immune cells, 3 kinds of vascular-associated cell subsets, and 2 kinds of Schwann cells. Analysis of cell–cell communication networks revealed that innate lymphoid cells were closely associated with other cells.

**Conclusions:** This study provides the first comprehensive transcriptome atlas and related database of mouse ELG. This resource can aid in a deeper understanding of lacrimal gland biology and pathophysiology of its related diseases.

## 1. Introduction

The lacrimal gland is an exocrine organ that secretes the aqueous layer of the tear film [1, 2]. Tear film components, water, mucin, lipids, electrolytes, and antibacterial peptides, are important for lubricating and protecting the ocular surface from external damage [3, 4]. Obstruction of tear production causes different degrees of ocular surface dryness and may lead to dry eye disease [5]. Dry eye disease is one of the most common ophthalmic diseases, and affects approximately 5–50% of the global population, especially the elderly [6, 7]. Therefore, an in-depth understanding of the structural composition and physiological function of the lacrimal gland is essential.

The lacrimal gland is composed of different cell types with interrelated functions [1, 8, 9]. Although traditional histology provides significant information to understand lacrimal gland structure, its constituent cells and molecular characteristics are yet to be elucidated. The emergence of single-cell RNA sequencing (scRNA-seq) technology provides an important tool to determine the transcriptome of a single cell that can identify cell types in an unbiased manner [10]. Recently, scRNA-seq was used to study the cellular diversity during early development of mouse extraorbital lacrimal glands (ELGs), mainly focusing on different epithelial cell lineages [11]. In addition, the cellular heterogeneity of the lacrimal gland has been elucidated using the lacrimal organoid to highlight the single-cell transcriptome characteristics of mouse and human lacrimal glands [12]. These studies helped in understanding the complexity of cell types in the lacrimal gland, especially that of epithelial cells.

The lacrimal gland is a special type of exocrine gland, mainly comprising the epithelia that secrete tears and are surrounded by mesenchymal company [13, 14]. The secreting epithelia are mainly composed of the glandular epithelial cells, acinar cells, ductal cells, and myoepithelial cells (MECs) [15]. Acinar cells constitute the main cells of terminal acini and secrete the main components of the aqueous layer in tear film. Similarly, one to two layers of cuboidal ductal epithelial cells, lined with ducts, secrete water, electrolytes, and peptides or proteins to modify the primary lacrimal gland fluid secreted by acinar cells [16–18]. Stellate-shaped MECs that surround and interact with glandular secretory acini assist lacrimal gland secretion [19]. These three kinds of epithelial cells (acinar, cuboidal ductal epithelial cells, and stellate shaped MECs) work together to secrete tear fluid to the ocular surface through the primary, secondary, and main ducts, respectively [20].

Meanwhile, the mesenchymal company, fibroblasts, immune cells, vessels, nerve fibers, pericytes around blood vessels, and Schwann cells surrounding nerve fibers are compartments that surround the epithelial cells of the lacrimal gland [15]. Immune cells are a heterogeneous cell population predominantly dispersed in the extracellular matrix, mainly consisting of T and B lymphocytes, macrophages, dendritic cells (DCs), monocytes, and mast cells [21–25]. These immune cells maintain the structural integrity and immunological condition of the lacrimal gland via different molecular mechanisms. Furthermore, plasma cells, differentiated from B lymphocytes, secrete immunoglobulins (Ig), especially IgA with secretory component forms, which maintain the immune stability of the ocular surface [26, 27]. Although extensive studies have identified lacrimal gland cellular diversity and corresponding functions, there are several unanswered questions.

The mouse model is the most studied representation of the physiological and pathophysiological functions of the human lacrimal gland [14, 28, 29]. Many of the findings and data are commonly applied to humans. Here, we generated an atlas of the transcriptome of 48,077 single cells based on 6 ELG samples from 6 donors based on the scRNA-seq to define 13 major cell types. This project revealed a previously unrecognized cellular heterogeneity and complexity in the lacrimal gland. In addition, this study provides many molecular markers for known and newly discovered cell types, providing a resource of lacrimal gland cellular composition, histology, physiology, and pathophysiology.

## 2. Materials and Methods

### 2.1. Animals

Twelve- to fourteen-week-old wildtype C57BL/6J male mice were purchased from the Nanjing Model Animal Institute of Nanjing University, China, and housed in light-tight circadian chambers, with controlled temperature (∼ 18–23 °C), humidity (50–60%), and ventilation. The mice had *ad libitum* access to food and water under a 12 h light/12 h dark cycle. All experimental procedures were approved by the Henan Province People’s Hospital Institutional Animal Care and Use Committee (HNEECA-2021-15) and performed in compliance with the Association for Research in Vision and Ophthalmology statement for the Use of Animals in Ophthalmic and Vision Research.

### 2.2. Tissue dissociation and single cell preparation

All ELGs were harvested at the same time (ZT3:00-ZT4:00) for each animal to avoid the influence of the circadian rhythm [30, 31]. The animals were euthanized by CO2 overdose and careful cervical dislocation to prevent perfusion. ELGs were collected from euthanized mice, centrifuged (4℃, 500 ×*g*) to remove the tissue preservation solution, washed twice with Eagle’s minimal essential medium, then quickly cut into 2–3 mm tissue sections using scissors. The tissue sections were incubated in an enzymatic digestion solution containing 2 mg/ml collagenase I, 2 mg/ml collagenase IV, and 1 mg/ml DNAas I enzyme in a C-tube, which was placed in a tissue processor for 1–2 h at 37 °C. The cell suspension was collected using a 40μm filter; erythrocyte lysis solution (QIAGEN. 8570396) was added to the filtered supernatant of the cell suspension, to remove erythrocytes, and centrifuged at 220 ×*g* for 8 min at 4 °C to remove the supernatant. Dead cells and debris were removed using the Dead Cell removal kit (Miltenyi Biotec no. 130-090-101) as per the manufacturer’s instructions. The cells were washed with phosphate-buffered saline (PBS), resuspended in an appropriate volume, and counted using a hemocytometer.

### 2.3. Single-cell RNA sequencing

For scRNA-seq experiments, two ELGs from each male mouse were combined into one sample; six ELG samples from six young mice were analyzed (n = 6). We focused on characterizing the heterogeneity and complexity of mouse lacrimal cells in the lacrimal gland based on the assumption of consistent cellular heterogeneity and complexity in the same species of age-matched mice with the same genetic background. After dissociation of the lacrimal gland tissue, single cells were diluted and suspended in calcium and magnesium free PBS containing 0.04% w/v bovine serum albumin. Approximately 10,000 cells, per sample, were added to each channel, and the target cell recovery was estimated at 8,000 cells. To generate single cell Gel bead-in-EMulsions (GEMs), the single cell suspension was loaded onto the Chromium Single Cell Controller (10X Genomics, Pleasanton, CA, USA). After generation of GEMs, reverse transcription reactions were performed using full-length cDNA with barcodes. The emulsion was disrupted using recovery agent and the cDNA purified with DynaBeads Myone Silane Beads (Thermo Fisher Scientific, Waltham, MA, USA). The cDNA was amplified for 14 PCR cycles corresponding to the concentration of the cDNA. Subsequently, the amplified cDNA was fragmented, end-repaired, A-tailed, and ligated to indexed articulators for library amplification. These libraries were sequenced through the Illumina HiSeq X Ten platform (Illumina, San Diego, CA, USA).

### 2.4. Data processing and quality control

To generate normalized summary data across samples, Cell Ranger (version 5.0.0, https://support.10xgenomics.com/single-cell-gene-expression/software/pipelines/latest/what-is-cell-ranger) was used to process raw data, perform multiplexed decomposition of cell barcodes, and map and downsample the reads of the transcriptome. This generated a raw unique molecular identifier (UMI) count matrix, which was converted to Seurat objects by R package Seurat (version 3.1.1).

To remove low-quality cells and likely multiplet captures, we assumed a Gaussian distribution of each cell’s UMI/gene number and filtered the cells out with UMI/gene numbers outside the limit of a mean value of +/– 2-fold of standard deviation. Based on the fraction of mitochondrial genes expressed, we visually inspected the cellular distribution and discarded low-quality cells where >10% of the counts belonged to mitochondrial genes. Additionally, we used the DoubletFinder package (version 2.0.2) [32] with default parameters to identify potential doublets. For populations of cells that were present only in a single sample, we defined them as interference signals and removed them. The final, remaining 48,077 single cells were used for downstream analysis. The counts matrix of each sample was merged through the cbind function in R, and the lognormalization operation was performed using the NormalizeData function in the Seurat package. Finally, to generate the integrated data matrix, the ScaleData function in the Seurat package was used for scaling. To remove the batch effects in single-cell RNA-sequencing data, mutual nearest neighbors (MNN) analysis was conducted to perform a dimensionality reduction analysis, based on the scaled integrated data matrix using the batchelor package. The FindAllMarkers function in Seurat was performed to identify marker genes of each cluster.

### 2.5. Cell type identification

Major cell clusters were identified using the FindClusters function provided by Seurat (default resolution was set to 0.8) and visualized using UMAP (uniform manifold approximation and projection). Multiple cell-specific enriched genes, previously described in literature, were used to determine the consistency of each cell type. To obtain the heterogeneity of each cell type, dimensionality reduction and clustering analyses were performed for each individual, major cell subtype.

### 2.6. Gene set variation analysis (GSVA) and Kyoto Encyclopedia of Genes and Genomes (KEGG) pathway enrichment analysis

R package GSVA27 (version:1.36.2, http://bioconductor.org/packages/release/bioc/html/GSVA.html) was used to perform GSVA with reference gene sets (c2.cp.kegg.v7.1.symbols.gmt and c5.go.bp.v7.2.symbols.gmt) from MSigDB v7.1.28 to compare GO_BP terms and KEGG pathways in the epithelial and fibroblast subset cells in the ELG. The enrichment scores of each GO_BP term and KEGG pathway were calculated to obtain a score matrix, followed by differential analysis using R package limma (version 3.44.3). A p value cutoff of 0.05 was set as the term for significant enrichment.

### 2.7. Trajectory analysis

To determine the potential relationship between different cell subsets of interest in murine ELG, Monocle2 (version 2.8.0) in R was used to determine pseudotime trajectories of some cell subsets [33]. Differentially expressed genes (DEGs) were determined for some of the cell subsets. The UMI matrix was used as input and variable genes detected by Seurat were used for building traces. Branches in the cell trajectory represent cells with alternative gene expression patterns. Cells were ordered through the orderCells function. The plot_pseudotime_heatmap function was used to visualize the scaled expression of branching curated genes associated with certain cell fates ordered according to pseudotime. Cells were manually selected as root based on the genes associated with highly proliferating amplified cells in each pseudo-time trajectory.

### 2.8. Cell–cell communication analysis

Cell–cell communication analysis was performed using CellPhoneDB (https://www.cellphonedb.org/; https://github.com/Teichlab/cellphonedb) [34] to explore potential interactions between different cell types in the mouse ELG. Single-cell transcriptome data, annotated as different cell types in the lacrimal gland, were entered into CellPhoneDB Python (package 2.0.0). Enriched receptor-ligand interactions between the two cell types were derived, based on the expression of one cell type for the receptor and the other for the corresponding ligand. The most relevant cell type-specific interactions between ligand and receptor were then identified. p values for the cell type-specific likelihood of receptor-ligand complex were obtained. Receptors or ligands with a p value < 0.05 were considered biologically relevant.

### 2.9. Immunohistochemistry and immunofluorescence of murine ELG

The histologic evaluation of murine ELG was executed using immunohistochemistry and immunofluorescence staining techniques as previously described [15, 35]. Briefly, the paraffin-embedded lacrimal gland was sectioned, deparaffinized, rehydrated, and incubated with different antibodies overnight at 4 °C to identify cell types (**Table. 1**). This was followed by incubation with the secondary antibody, and nuclei staining with Mayer hematoxylin solution (Cat # G1004, Servicebio Company, Wuhan, China). Previously described immunofluorescent technology was used for some cell types in the ELG [31].

The nucleus was labeled with 4’,6-diamidino-2-phenylindole (1:500; Sigma-Aldrich, St. Louis, MO). A DeltaVision high resolution image system (GE Healthcare, USA) was used to examine and image lacrimal gland slices. Isotype antibody (Rabbit IgG, GB111738, Servicebio Company) was used instead of the primary antibody as a negative control to exclude non-specific staining.

### 2.10. Statistical analysis

Statistical analyses were performed using Graphpad Prism software 7 (GraphPad Software Inc.) and R software (http://www.r-project.org). Data were presented as the mean and standard error of the mean (SEM) unless indicated otherwise. Beanplot R package was used to generate violin plots, with the data distribution bandwidth evaluated by kernel density estimation. A p value < 0.05 was considered statistically significant.

## 3. Results

### 3.1. Preparation of mouse ELG samples and generation of a single-cell transcriptome atlas

Six fresh lacrimal gland samples were collected from adult mice to obtain sufficient cell numbers. Samples were sent to the laboratory within 8 h, single-cells were dissociated, and scRNA-seq analysis was performed using the 10X Genomics Chromium platform (**Fig. 1A**). Cell Ranger quantitative quality control was performed to ensure a high-quality cell count distribution between 9,245 and 15,122. Standard QC measures were implemented to exclude bicellular, multicellular, and apoptotic cells. The final cell numbers, the average number of genes per cell, the average UMI per cell, and the average mitochondrial gene ratio per cell of each sample was 6,517–9,014; 794–1,319; 2,334–10,240; and 0.023–0.033, respectively (**Fig. S1)**. Based on the similarity of transcriptome features, the scRNA-seq data were analyzed by an unsupervised graph clustering method using Seurat to classify individual cells into cell populations. UMAP for data visualization showed 21 different cell clusters based on gene expression profiles, indicating that there were different cell types at the single-cell transcriptomic level (**Fig. 1B, C and Fig. S2)**. Marker genes and cell numbers for each cluster of integrated 6 samples and each sample were shown in **Table. S1, Table. S2, and Table. S3**.

**Fig. 1.**
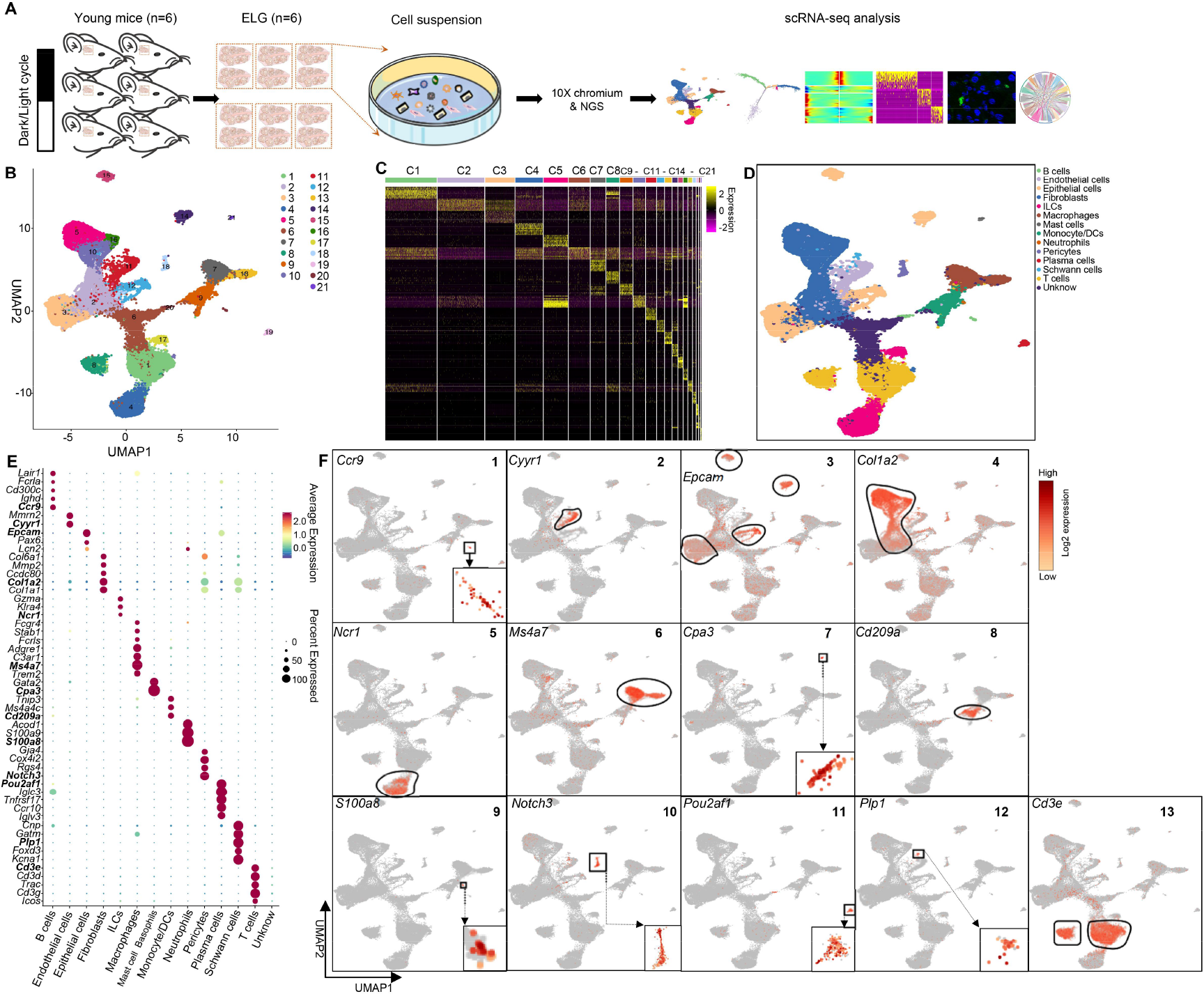
Global single-cell transcriptome landscape and the identification of cell populations from six mouse ELGs by scRNA-seq. **(A)** Schematic workflow for scRNA-seq. Murine ELGs were used to generate data using the 10X Genomics system that allows the quantification of transcript abundance across cells and genes. **(B)** UMAP visualization of adult ELG cells from six donors for cluster identification. Each color represents a distinct cell population. **(C)** Heatmap of Z scores for scaled expression values of DEGs for 21 clusters. The specific genes and their relative expression levels in all sequenced lacrimal gland cells are shown for each cluster. Color scale: purple for low expression, yellow for high expression. Gene list of the heatmap for each cluster was shown in **Table. S5**. **(D)** UMAP visualization of ELG cells for cluster identification. Each color represents a distinct cell population. **(E)** Normalized expression levels of selected DEGs from ELG cell populations shown in (D). The gene expression frequency of each cell type is indicated by the spot size, and the expression level is indicated by color intensity. Color scale: red for high expression and blue for low expression. **(F)** UMAP visualization illustrates the expression of representative marker genes in 13 major cell types in (E). DEG, differentially expressed gene; scRNA-seq, single-cell RNA-sequence; C, cluster; ELG, extraorbital lacrimal gland.

### 3.2. scRNA-seq–based identification of cell types in murine ELG

Cell type identity was assigned to clusters, based on previously reported expression of unique or specific marker genes. Thirteen cell types were distinctly clustered based on the marker genes in **Fig. 1D, E, F**: (1) epithelium characterized by *Epcam* [36] and *Lcn2* [37]; (2) fibroblast by *Pdgfra* [38]*, Col1a1* [39], and *Col1a2* [39]; (3) T cells by *Cd3d*/*e*/*g* [40] and *Trac* (T cell receptor alpha constant) [41], *Trbc1* (T cell receptor beta constant 1) [42], or *Trbc2* (T cell receptor beta constant 2) [43]; (4) B cell by *Cd79a* [39], *Cd79b* [44], and *Ccr9* [45], and plasma cells by *Pou2af1* [46] and *Tnfrsf17* [47]; (5) monocyte/DCs by *Flt3* [48]*, Cd209a* [48], and *Tnip3* [45]; (6) macrophages by *Cd68* [49], *Ms4a7* [50], *Adgre1* [51], and *Trem2* [52]; (7) natural killer (NK) cells and innate lymphoid cell 1 (ILC1) by *Ncr1* [36]*, Gzma* [53], and *Klra4* [41]; (8) ILC2s by *Il1rl1* [54]; (9) mast cells by *Cpa3* [55]*, Gata2* [56], and *Ccl2* [57]; (10) neutrophils by *S100a8* [43]*, S100a9* [58], and *Acod1* [59]; (11) endothelial cells by *Pecam1* [60], *Cdh5* [61] and *Cyyr1* [62]; (12) pericytes by *Rgs4* [63]*, Des* [64], and *Notch3* [65]; and (13) Schwann cells by *Plp1* [66]*, Kcna1* [67], and *Cnp* [68]. Altogether, our datasets provide a comprehensive resource to further identify and characterize cell types of murine ELG at high resolution. The differentially expressed marker genes in **Fig. 1E** for each celltype were shown in **Table. S4**.

### 3.3. Glandular epithelial cells in murine ELGs

Adult lacrimal gland epithelial cells mainly consisted of terminally differentiated acinar cells, ductal cells, and MECs. Epithelial cells identified by the epithelial cell-specific marker genes *Epcam and Lcn2,* were initially analyzed to elucidate the heterogeneity and transcriptomic profiling of the lacrimal gland epithelial cell population (**Fig. 1E, F**). All glandular epithelial cell populations were composed of 9 distinct clusters (**C1–C9**) under UMAP visualization (**Fig. 2A, B)**. The top 10 pathways in each cluster were determined with t-values ranked from largest to smallest and p value< 0.05 were plotted on a heat map by the gene set variation analysis (GSVA) for Gene Ontology (GO)-biological process (BP) terms and KEGG pathways, respectively [69, 70]. Differential genes expressed by each cluster of cells were enriched in different GSVA-GO-BP terms and GSVA-KEGG pathways (**Fig. S3**). Cluster 9 mainly expressed GSVA-GO terms and KEGG pathways related to cell division and the cell cycle (**Fig. S3**).

**Fig. 2.**
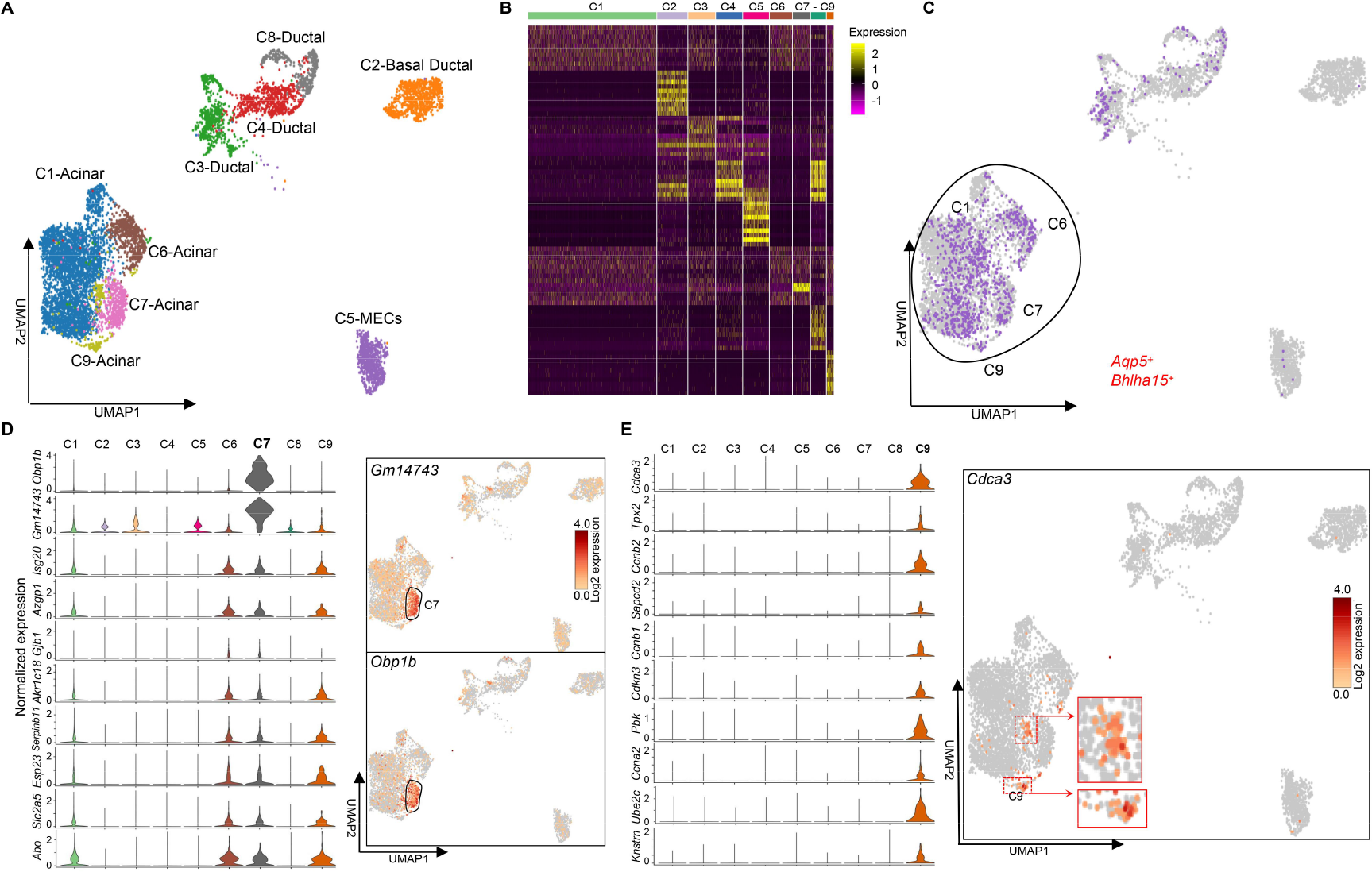
Identification of lacrimal gland epithelial cell clusters by scRNA-seq. **(A)** Lacrimal gland epithelial cell transcriptome visualized with UMAP and colored according to unsupervised clustering. **(B)** Heatmap showing specific genes for each of the nine clusters (top 10 marker genes in each cluster). Color scale: purple for low expression and yellow for high expression. The gene list in the heatmap for each cluster is shown in Table. S6. **(C)** *Aqp5* and *Bhlha15* co-expressed in acinar epithelial subsets of the murine ELG. **(D)** Violin plot of the top 10 marker genes expressed by **Cluster 7** (*left*) and UMAP visualization of *Gm14743* and *Obp1b* (*right*). Color scale: dark brown, high expression; yellow, low expression. **(E)** Violin plot of the top 10 marker genes expressed by **Cluster 9** (*left*) and UMAP visualization of proliferative gene *Cdca3* (*right*).

#### Acinar cells of the mouse ELG

**Cluster 1, 6, 7, and 9** accounted for 73.35%, 12.91%, 9.80%, and 3.94% of all lacrimal acinar cells, respectively, and they highly co-expressed *Aqp5* and *Bhlha15* (**Fig. 2C**), suggesting that these cell clusters represent acinar epithelial cells [18, 35, 71–74]. Further analysis revealed that the cells in Clusters 1, 6, and 7 express similar top 10 marker gene patterns (**Fig. S4**). However, the cells in **Cluster 7** preferentially expressed two odorant-binding protein associated genes: *Gm14743,* which encodes a protein superfamily sharing structural features of the lipocalins, fatty acid-binding proteins, avidins, triabin, and a group of bacterial metalloprotease inhibitors [75], and *Obp1b* (encoding odorant-binding protein 1b), which is mainly localized to secretory glands including the lacrimal glands, binds chemical odorants [76, 77] (**Fig. 2D**). The cells in **Cluster 9** distinctively expressed mitotic cell cycle machinery-related genes (*Cdca3, Ccnb2, Pbk,* and *Ube2c*) (**Fig. 2E**, left) with ***Cdca3*** being the most highly expressed (**Fig. 2E,** right). The top functional enrichment of KEGG and GO for **C1** (marker gene: *Mrap*, *Esp3*, and *AI463170*), **C6** (marker gene: *St6galnac1*, *Mrap*, and *Akr1c18*), and **C7** (marker gene: *Obp1b, Gm14743*, and *Isg20*) are shown in **Fig. S4**.

The pseudotime trajectory was analyzed to understand the possible relationship for variations among the four glandular acinar epithelial cell clusters (**C1, C6, C7,** and **C9**). As cells in **Cluster 9** strongly and exclusively expressed genes related to highly proliferating transit-amplified cells (*Cdca3*, *Ube2c*, *Ccnb2*, *Pbk*, *Cdkn3*, *Ccnb1*, *Ccna2*, *Knstrn*, *Tpx2*, and *Sapcd2*) (**Fig. 2E**), this cluster was assumed to be the starting cell (Pre-branch) that drove the differentiation of normal glandular acinar cells to Branches 1 (State 1) and 2 (States 3, 4, and 5) (**Fig. 3A–C**).

**Fig. 3.**
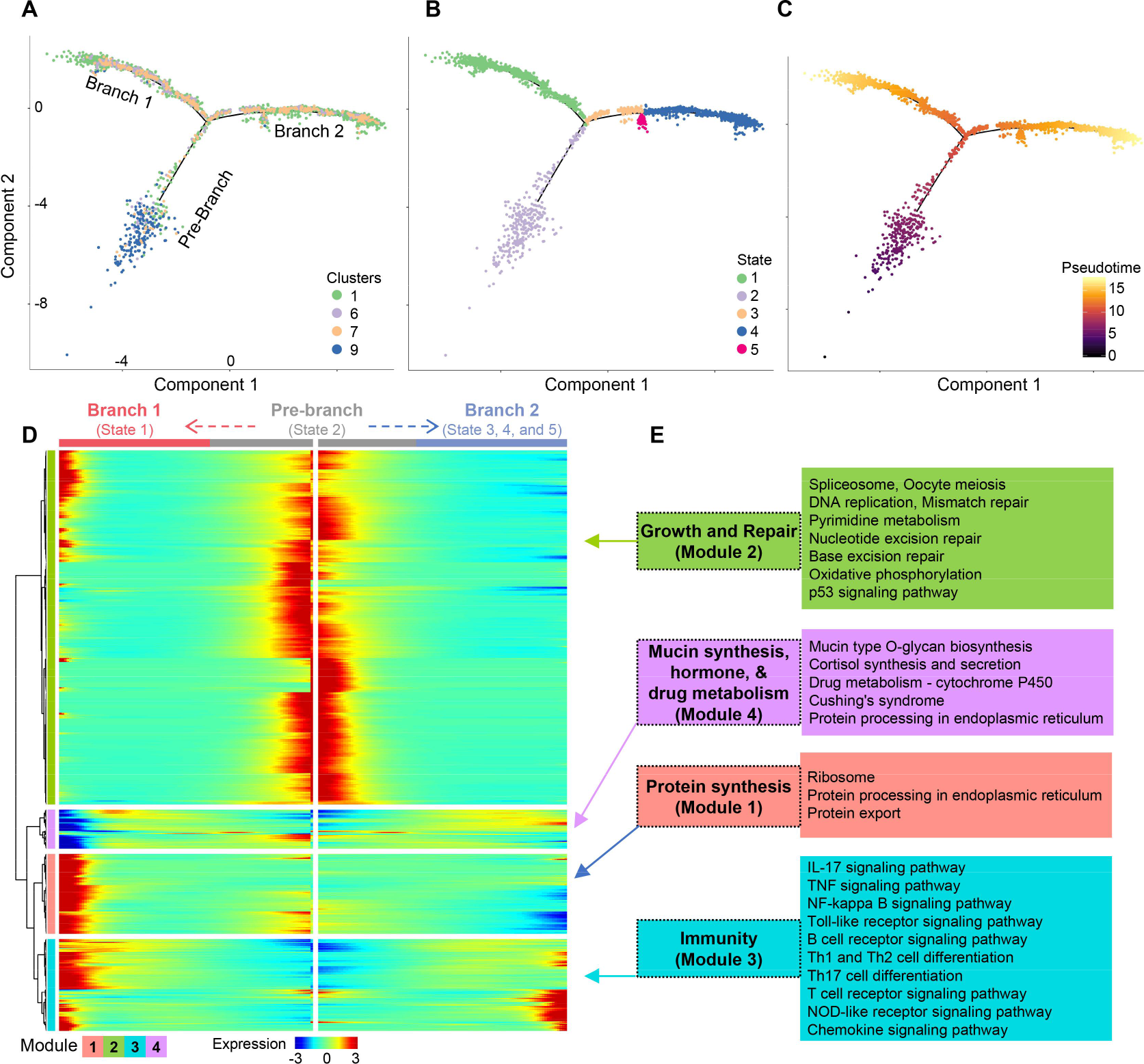
Subpopulation of glandular acinar cells and their pseudotime trajectory in the murine ELG **(A)** Trajectory reconstruction of all single acinar epithelial cells into distinct cell clusters. Different colors represent different cell clusters revealing three branches: Pre-branch (before bifurcation), Branch 1, and Branch 2 (after bifurcation). **(B)** Trajectory reconstruction of all single acinar epithelial cells based upon the states. **(C)** Trajectory reconstruction of all single acinar epithelial cells based upon pseudotime. Color scale: darker colored dots (Pre-branch) represent earlier differential stage cells and lighter colored dots (Branches 1 and 2) represent later mature cells. **(D)** Kinetic heatmap generated using BEAM regression model capturing the most DEGs between Branches 1 and 2. The red and blue colors represent upregulation and downregulation, respectively. **(E)** Top functional KEGG pathways from four different gene expression modules.

Kinetic analysis of DEGs from four different acinar epithelia clusters was performed to determine the differentiation process- and biological functions of the lacrimal acinar epithelial cells. Most of the highly expressed genes were categorized into four modules (**Fig. 3D**). **Module 1** mainly contained genes highly expressed in States 1 of Branch 1, which were enriched in protein synthesis process; **Module 2** mainly contained genes highly expressed in the Pre-branch state, which were involved in growth and repair processes; **Module 3** mainly contained genes highly expressed in Branches 1 and 2, involved in innate- and adaptive immune processes; **Module 4** was expressed in states 2-5 and was mainly involved in mucin synthesis, cortisone secretion and synthesis, and drug metabolism (**Fig. 3E**). In conclusion, there was a dynamic process of differentiation between lacrimal acinar epithelial populations and involvement in different physiological processes and functions.

#### Myoepithelial cells in murine ELG

Myoepithelial cells (MECs) are a unique cell type containing epithelial and smooth muscle cell phenotypes. They surround the glandular secretory acini, express α-smooth muscle actin, have a contractile function, and are primarily responsible for lacrimal gland secretion [15, 19]. **Cluster 5** cells (accounting for 8.63% of all lacrimal gland epithelial cells) expressed high levels of MEC marker gene *Acta2* [78] (**Fig. 4A**). This cell cluster consistently displayed high expression of classical genes associated with MEC function, including *Myl9, Mylk, Tpm2, Myh11,* and *Cnn1* (encoding calponin-1, a basic smooth muscle protein) [19] (**Fig. 4A**). Thus, **Cluster 5** was clearly identified to MECs in the ELG. Moreover, MECs in mouse ELG specifically expressed *Pcp4* (encoding Purkinje cell protein 4), *Osr1* (encoding odd-skipped related transcription factor 1, participating in the normal development of body parts), *Lmod1* (encoding leiomodin 1 protein), *Prkg1* (encoding cGMP-dependent protein kinase 1), *Dmpk*, *Cck* (encoding cholecystokinin), and *Gm9885* (**Fig. 4B**). Notably, these MECs uniquely expressed aquaporin 4 (*Aqp4*), a water-selective channel that is mainly found in the brain and is important for water homeostasis [79] (**Fig. 4B**). Additionally, cellular immunohistochemistry techniques confirmed the presence of MECs and surrounding acinar cells (**Fig. 4C**). Finally, functional enrichment analysis of DEGs in this cell subpopulation by KEGG showed muscle contraction and other related functions (**Fig. 4D**).

**Fig. 4.**
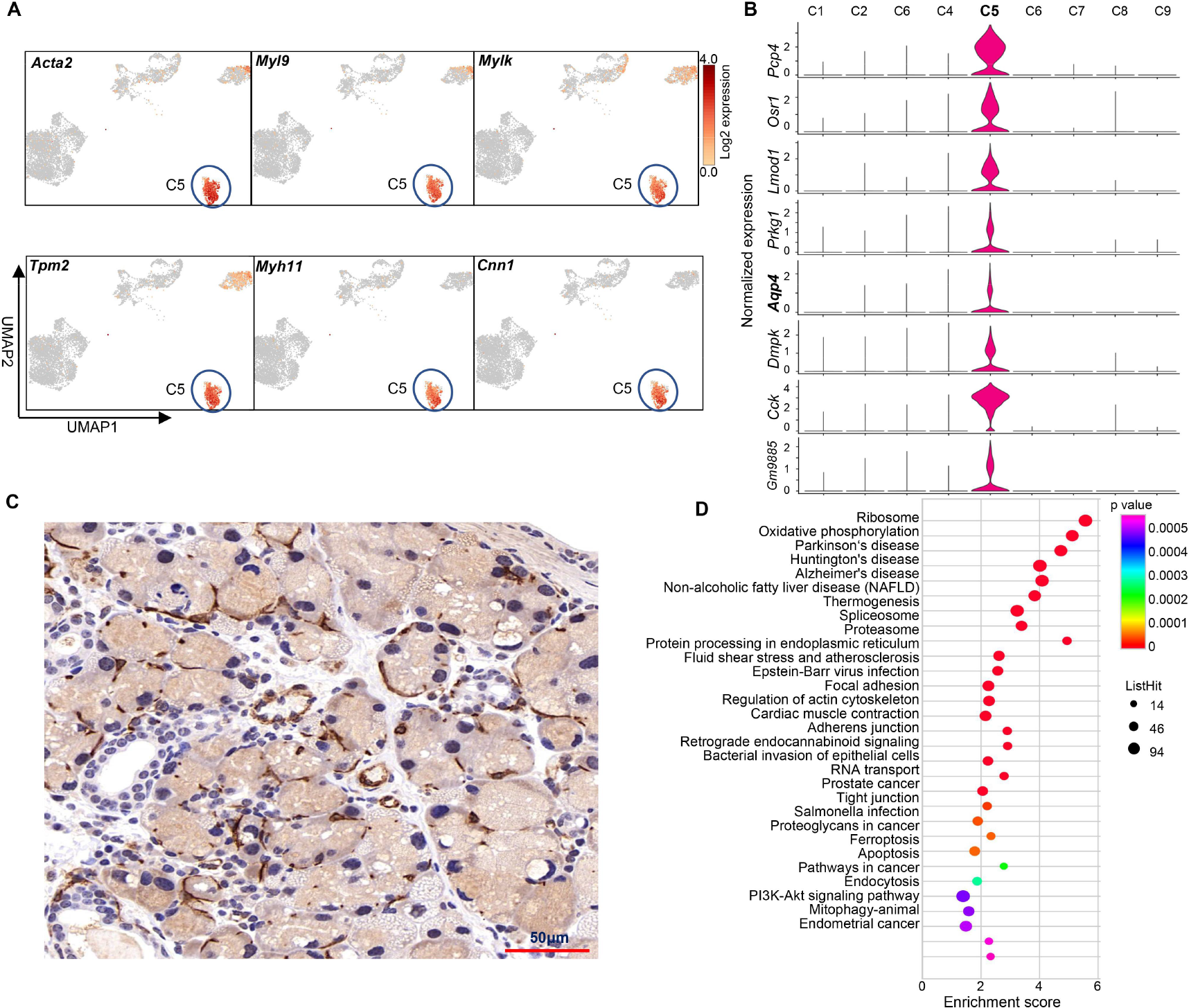
Identification of myoepithelial cells (MECs) in the murine ELG. **(A)** Identification of lacrimal gland MECs with classic marker genes visualized with UMAP Color scale: dark brown, high expression; yellow, low expression. **(B)** Violin plot of cluster-specific top 10 marker genes of **Cluster 5** cells (MECs) in ELG. **(C)** Immunostaining with a myoepithelial cell marker, anti-alpha smooth muscle actin (brown) under 40X magnification. **(D)** Functional KEGG analysis of MECs-expressed DEGs in mouse ELG.

#### Ductal epithelial cells of mouse ELG

Na^+^/K^+^ ATPase is enriched in lacrimal ductal cells by 3–5 times compared with acinar cells [18, 80, 81]. Single cell resolution showed that **Clusters 2** (10.20%), **3** (8.98%), **4** (8.75%), **5** (8.63%), and **8** (5.09%) in all lacrimal gland epithelial cells were relatively high expression of Na^+^/K^+^ ATPase family genes (*Atp1a1, Atp1b1,* and *Atp1b3*) (**Fig. 5A, B**).

**Fig. 5.**
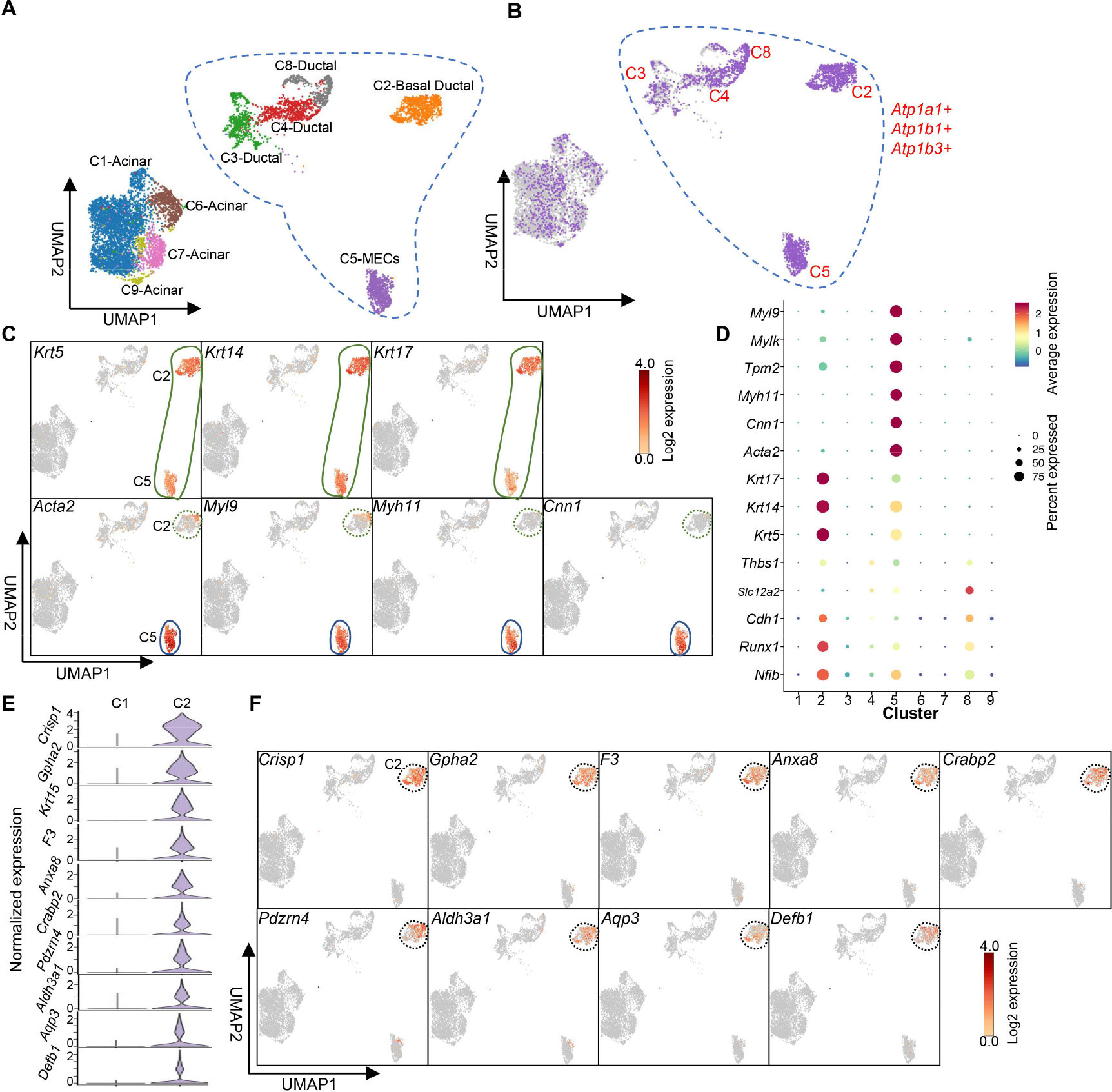
Identification of ductal cell subsets in the mouse ELG by scRNA-seq. (A, B) Identification of lacrimal gland ductal epithelial cells visualized with UMAP (A). *Atp1a1*, *Atp1b1*, and *Atp1b3* co-express in the ductal epithelial subsets in ELGs (B). **(C)** UMAP visualization of marker genes of **Clusters 2** and **5** subset cells. **(D)** Dotplot of top 14 marker genes for basal ductal cells and MECs in 9 different clusters. **(E)** Violin plot of top 10 marker genes of **Cluster 2** cells. **(F)** UMAP visualization of marker genes of **Cluster 2** subset cells.

Thus, we infer that these cell clusters represent ductal cells in the lacrimal gland. However, **Cluster 2** specifically co-expressed classical basal cell keratins *Krt5*, *Krt14,* and *Krt17* (**Fig. 5C** upper, **D**) with MECs, but not MEC-specific marker genes *Acta2*, *Myl9, Myh11,* and *Cnn1* (**Fig. 5C** lower, **D**). Thus, **Cluster 2** can be defined as a basal ductal cell subset [19]. In addition, *Crisp1*, *Gpha2*, *F3*, *Anxa8*, *Crabp2*, *Pdzrn4*, *Aldh3a1*, *Aqp3*, and *Defb1* marker genes were specifically expressed by mouse ELG ductal basal cells **(Fig. 5E, F)**.

Further analysis showed that **Cluster 8** specifically expressed luminal progenitor marker genes, *Kit* and *Cd14* [**Fig. 6A** (1,2)]. This indicated that **Cluster 8** represents c- *Kit*^+^ intercalated duct cells, which have progenitor properties as defined in the salivary and mammary glands [82, 83]. **Cluster 8** also highly expressed *Cxcl17*, a small mucosal chemokine that is widely expressed in different mucosal tissues [84], *Sftpd* (encoding surfactant protein D), a multimeric collectin that is involved in innate immune defense, and *Ltf* (encoding lactoferrin, LTF), a secreted protein important for ocular surface defense against microbial attack [85] [**Fig. 6A** (3, 4, 5)]. Moreover, it uniquely expresses cell growth, death, and repair-related genes such as *Lratd1* (encoding LRAT domain containing 1, involved in cell morphogenesis and cell motility), *Tspan8* (encoding tetraspanin 8, which mediates the regulation of cell development, activation, growth, and motility) [86], and *Cidea* (encoding cell death inducing DFFA like effector A, which activates apoptosis); ionic channel-related genes such as *Slco4c1* (encoding solute carrier organic anion transporter family member 4C1, belonging to the organic anion transporter family) and *Clic6* (encoding chloride intracellular channel protein 6); and nervous activity-related genes such as *Ncald* (encoding neurocalcin delta) and *Ramp1* (encoding receptor activity modifying protein 1) (**Fig. 6B**).

**Fig. 6.**
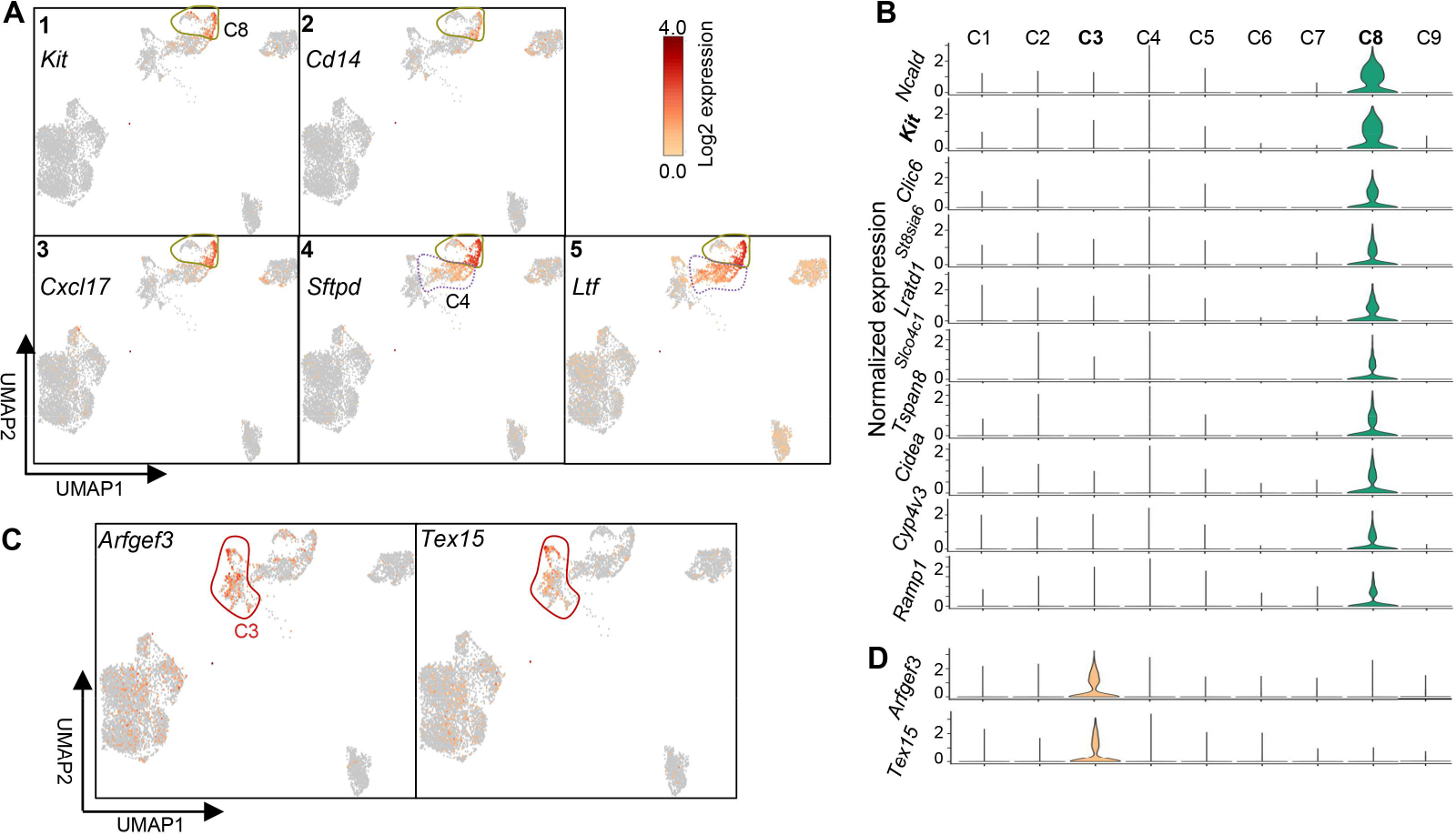
Identity of duct epithelial cell subsets in Clusters 3 and 8 of the ELG. **(A)** Visualization of *Kit, Cd14, Cxcl17, Sftpd,* and *Ltf* expression in UMAP. Color scale: dark brown, high expression; yellow, low expression. **(B)** Top 10 genes specifically expressed by the cell subset of **Cluster 8** in ELG. **(C)** Visualization of *Arfgef3* and *Tex15* expression of **Cluster 3** in UMAP. **(D)** Violin plots of *Arfgef3* and *Tex15* expression levels and gene distribution in **Cluster 3**.

Notably, **Cluster 4** co-expressed *Sftpd* and *Ltf* together with **Cluster 8 [Fig. 6A** (4, 5)]. However, **Cluster 4** did not express the **Cluster 8**-specific genes, *Kit, Cd14,* and *Cxcl17* **[Fig. 6A (**1, 2, 3)]. As surfactant protein D (SFTPD) is enriched in ductal cells of adult human lacrimal glands [87] and also expressed by ductal cells of mouse ELG [11], we defined **Cluster 4** as a ductal cell population of mouse ELG. **Cluster 3** only expressed two characteristic markers: *Arfgef3* (encoding ADP ribosylation factor guanine nucleotide exchange factor 2) and *Tex15* (encoding testis expressed 15) compared to other ductal cell subpopulations (**Fig. 6C, D**).

### 3.4. Fibroblasts in murine ELG

The fibroblasts in the mouse ELG were distinguished by their characteristic marker genes *Pdgfra*, *Col1a1*, and *Col1a2* **(Fig. 1E, F)**. Based on the similarity of differential gene expression and the reference database, 7 different fibroblast clusters were identified in murine ELG under a steady state: *Lipo1^+^*(**C1**) (33.80%), *Ccl5^+^*(**C2**) (30.98%), *Lsamp^+^*(**C3**) (20.05%), *Serping1*^+^(**C4**) (9.74%), *Smpd3^+^*(**C5**) (3.76%), *Mfap4*^+^(**C6**) (0.93%), and *Chia1*^+^(**C7**) (0.74%) **(Fig. 7A, B and Fig. S5A)**. Their DEGs are enriched in different biological processes and signaling pathways (**Fig. S5B, C**). Two universal fibroblast populations were also verified in mouse ELG: *Col15a1*^+^ adventitial fibroblasts (**Fig. 7C**) and *Pi16^+^* parenchymal fibroblasts (**Fig. 7D**), which is consistent with the finding of a recent study that integrated single-cell transcriptomic data from multiple organ fibroblasts [88]. The cell subset in *Lsamp^+^* **Cluster 3** was defined as *Col15a1*^+^ adventitial fibroblasts because this cluster of cells specifically expressed *Col15a1* and *Penk* [88] (**Fig. 7C**), whereas the cell subset in **Cluster 5** were defined as *Pi16^+^* parenchymal fibroblasts because this cluster of cells specifically expressed *Pi16, Dpp4,* and *Ly6c1* [88] (**Fig. 7D**). Other top marker genes uniquely expressed by the two clusters of cells (*Col15a1*^+^ **C3** and *Pi16^+^* **C5**) were also shown in a lacrimal niche-specific feature, respectively (**Fig. 7E**). Of note, these two clusters of cells uniquely expressed high levels of resident stem cell-associated genes: *Cd34* [89] and *Ly6a* [encoding lymphocyte activation protein-6A, also known as stem cell antigen-1 (Sca-1)] [90, 91] (**Fig. 7F)**. This evidence suggested a trajectory relationship among fibroblast clusters. Therefore, the Monocle2 algorithm was used to construct dynamic processes between different cell clusters along pseudotime trajectories based on the transcriptome level of each cluster cells. The results suggest that different cell clusters formed a tree-like structure (**Fig. 8A– C**). Cell populations, that exhibited stemness characteristics, of **Clusters 3** and **5** were located at the initiating state and differentiated into two different fates, Branches 1 and 2. Before bifurcation, **Clusters 3** and **5** gradually mainly evolved into **Cluster 4** and a small number of **Cluster 6** populations; Branch 1 consisting of **Clusters** 2, 4 and 7; Branch 2 consisting of mostly **Cluster 1** cells and a small number of Cluster 4 cells. Collectively, our analysis suggests that the lacrimal gland has two kinds of universal fibroblasts (*Pi16*^+^ and *Col15a1*^+^) and 5 possible universal cell-derived fibroblasts in the steady-state.

**Fig. 7.**
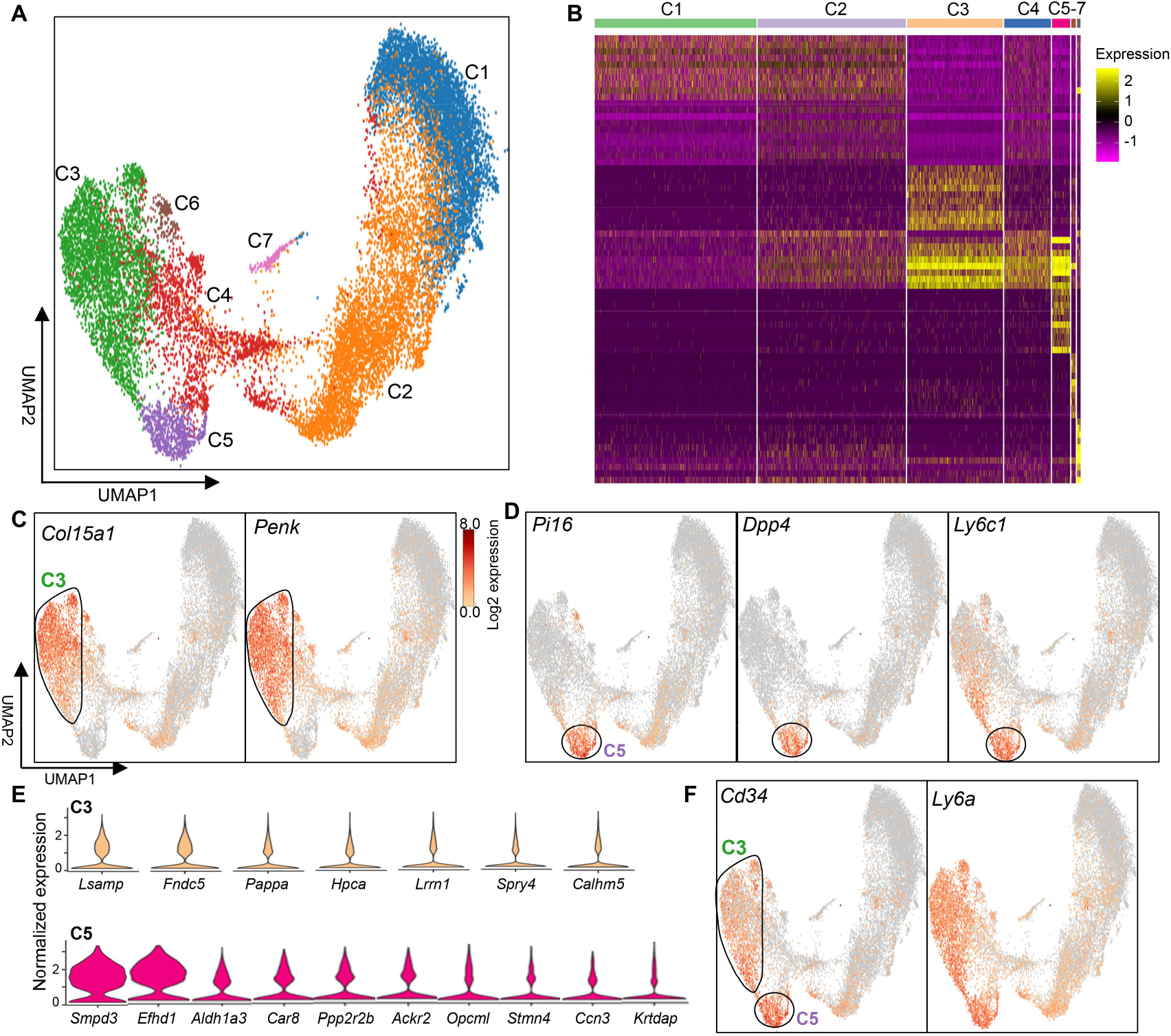
Subpopulation of fibroblasts in the murine ELG at the scRNA-seq level. **(A)** UMAP visualization of all fibroblast subsets by cluster ID. **(B)** Heatmap showing top 10 fibroblast marker genes for each of the 7 clusters. Color scale: yellow for high expression; purple for low expression. Gene list of the heatmap for each cluster was shown in **Table. S7**. **(C)** Expression of adventitial fibroblast specific-marker genes *Col15a1* and *Penk* in **Cluster 3**. Color scale: dark brown, high expression; yellow, low expression. **(D)** Expression of parenchymal fibroblast marker genes *Pi16*, *Dpp4*, and *Ly6c1* in Cluster 5. **(E)** Top cluster-specific marker genes of **Clusters 3** and **5** for fibroblasts in the ELG. **(F)** Co-expression of stem cell marker genes *Cd34* and *Ly6a* in **Clusters 3** and **5** of fibroblasts.

**Fig. 8.**
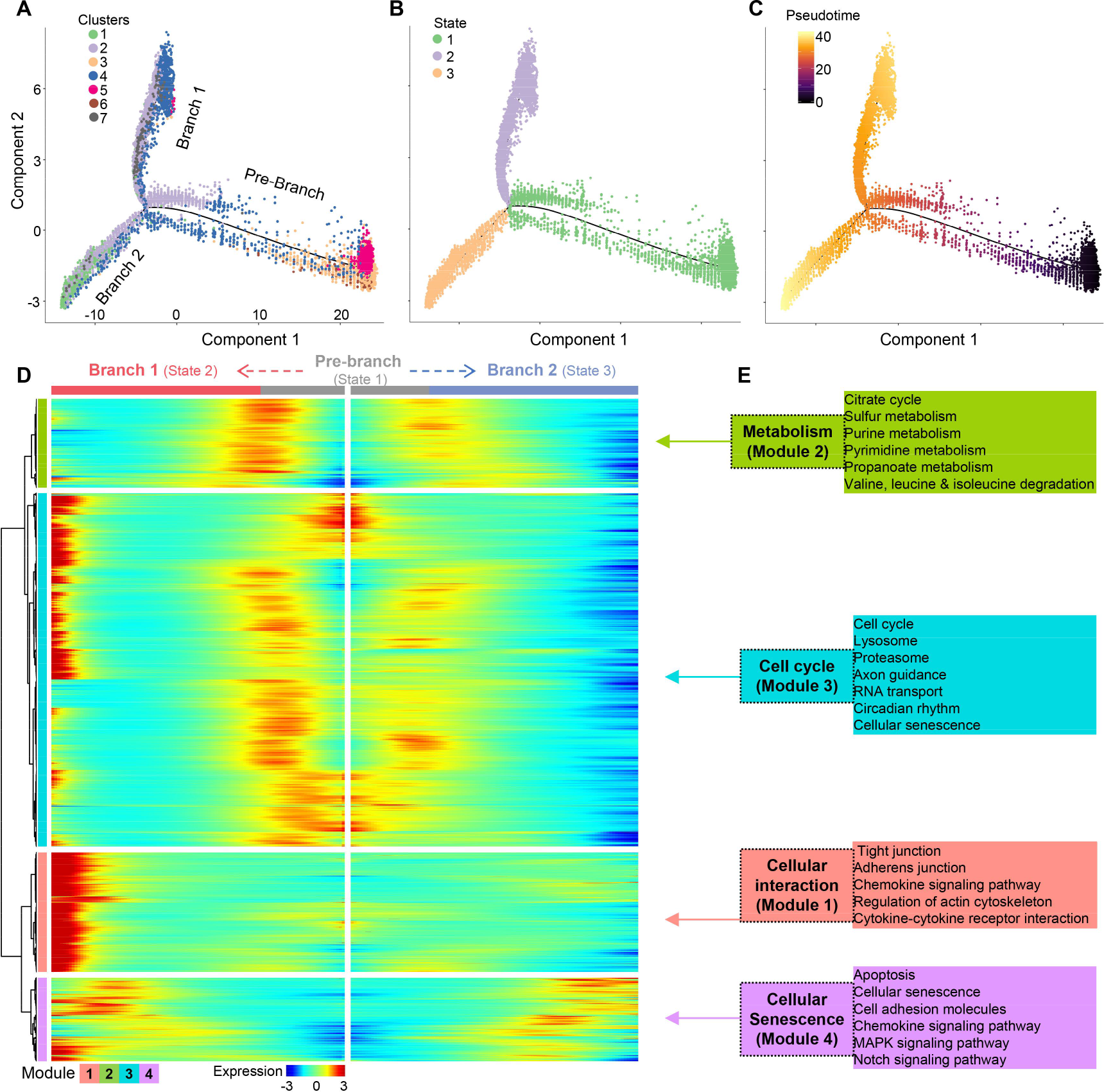
Pseudotime trajectory analysis of the fibroblast single-cell transcriptome in the murine ELG. **(A)** Trajectory reconstruction of all single fibroblast subsets based on cell clusters. Different colors represent different cell clusters revealing three branches: pre-branch (before bifurcation), Branch 1, and Branch 2 (after bifurcation). **(B)** Trajectory reconstruction of all single fibroblast subsets based on the states. **(C)** Trajectory reconstruction of all single fibroblast subsets based on the pseudotime. Color scale: darker colored dots (Pre-branch) represent earlier differential stage cells and lighter colored dots (Branches 1 and 2) represent later mature cells. **(D)** Kinetic heatmap generated using the BEAM regression model capturing top DEGs between States 1 and State 2. Red and blue colors represent upregulation and downregulation, respectively. **(E)** Top Functional KEGG pathways from four different gene expression modules. KEGG-Kyoto Encyclopedia of Genes and Genomes.

Kinetic analysis of DEGs in different fibroblast subpopulations was performed using Monocle 2 to determine the physiological processes and biological functions of the fibroblast population in the lacrimal gland. The pseudotime expression of the fibroblast population formed four modules, which were almost evenly distributed in two branches (two states) (**Fig. 8D**). Module 1 was mainly enriched in cellular interaction related processes; Module 2 was mainly involved in basic metabolism processes; Module 3 was mainly involved in cell cycle functions; and Module 4 was involved in cellular senescence processes (**Fig. 8E**). In conclusion, fibroblasts in the murine lacrimal gland are involved in different physiological processes and functions through a dynamic differentiation process.

### 3.5. Vascular-associated cells in the ELG

Blood vessels are mainly composed of vascular endothelium and vascular mural cells.

The blood vessel endothelial cell population was initially distinguished by *Pecam1 and Cyy1* to understand vascular-related cell heterogeneity in the lacrimal gland (**Fig. 1E, F and Fig. 9A**). There were three heterogeneous endothelial cell subsets in murine lacrimal glands (**Fig. 9A–F**). **Cluster 1** cells (39.81%) co-expressed endothelial and fibroblast/mesenchymal proteins (*Col1a2, Col3a1,* and *Col1a1*) (**Fig. 9D**), indicative of cells “transitioning” from the endothelium to the mesenchymal, a fundamental process during development [92, 93]. **Cluster 2** cells (37.55%) express *Esm1* (endothelial cell-specific molecule 1) and *Rgcc* (encoding protein regulator of cell cycle) as a capillary marker [94] (**Fig. 9E**). **Cluster 3** cells (22.64%) express *Vwf* and *Vcam1*, which are large vessel endothelial markers (**Fig. 9F**), and the expression level correlates with vessel diameter [95]. As predicted, DEGs from each of the three endothelial cell subpopulations were differentially enriched in signaling pathways related cell junctions, leukocyte trans- endothelial migration, and fluid shear stress (**Fig. 9G–I**). These data confirm the heterogeneity of vascular endothelial cells in the mouse lacrimal gland, as in the other tissues recently studied [94].

**Fig. 9.**
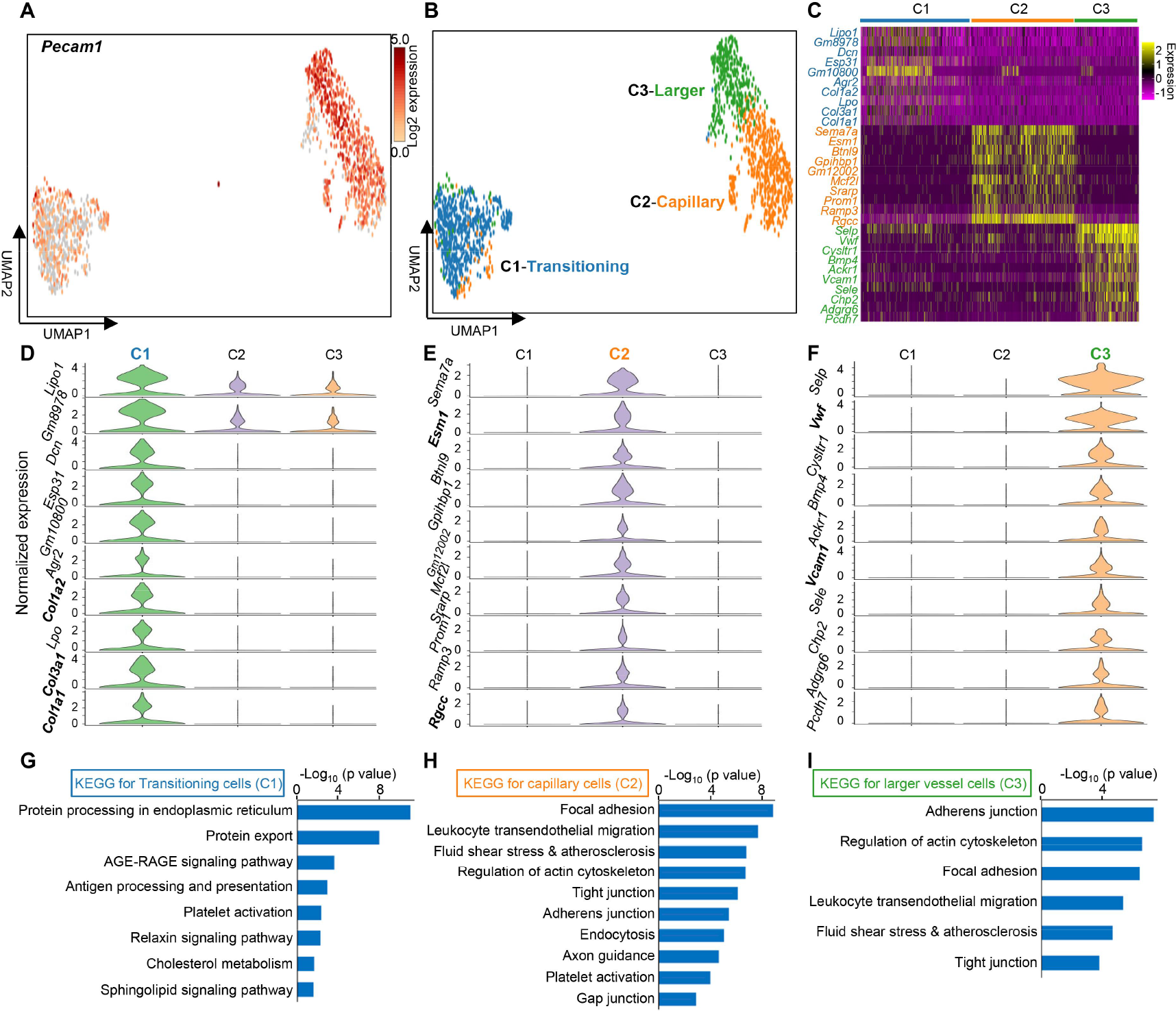
Subpopulation of vascular endothelial cells in the murine ELG. **(A)** UMAP visualization of *Pecam1*^+^ endothelial cells clusters in the lacrimal gland. Color scale: dark brown, high expression; yellow, low expression. **(B)** UMAP visualization of three distinct endothelial cell clusters. **(C)** Heatmap of the top 10 marker genes for each cluster of vascular endothelial cells. Color scale: dark for low expression, yellow for high expression. **(D–F)** Violin plots of the top 10 marker genes for each cluster [Cluster1-(D), Cluster2-(E), Cluster3-(F)] of vascular endothelial cells. **(G–I)** Functional KEGG analysis for transitioning (C1)-, capillary (C2)-, and larger (C3) endothelial cells.

Cells with vascular mural cell characteristics were down-dimensioned into two cell populations at a single-cell resolution (**Fig. 10A, B**). **Cluster 1** cells (70.15%) specifically expressed *Kcnj8, Vtn,* and *Abcc9*, indicative of pericytes [96] (**Fig. 10C, D**). However, **Cluster 2** (29.85%) specifically expressed *Tagln* (encoding transgelin-2), *Myh11* (encoding myosin heavy chain 11), and *Acta2* (encoding actin alpha 2 for smooth muscle), indicative of smooth muscle cells (SMCs) [97] (**Fig. 10E, F**). Functional KEGG analysis from the DEGs expressed by pericytes showed that pathways were significantly associated with focal adhesion, fluid shear stress and atherosclerosis, relaxin-signaling, vasopressin-related water reabsorption, and chemokine signaling (**Fig. 10G**). Pathways from the DEGs expressed by SMCs were associated with vascular smooth muscle contraction, regulation of actin cytoskeleton, tight junction and adherens junction (**Fig. 10H**). Additionally, SMCs of the vessel wall were verified through immunohistochemical staining using anti-acta2 antibody (**Fig. 10I**). Collectively, these findings provide basic information to understand transcriptomic profiling of vascular mural cells in the lacrimal gland.

**Fig. 10.**
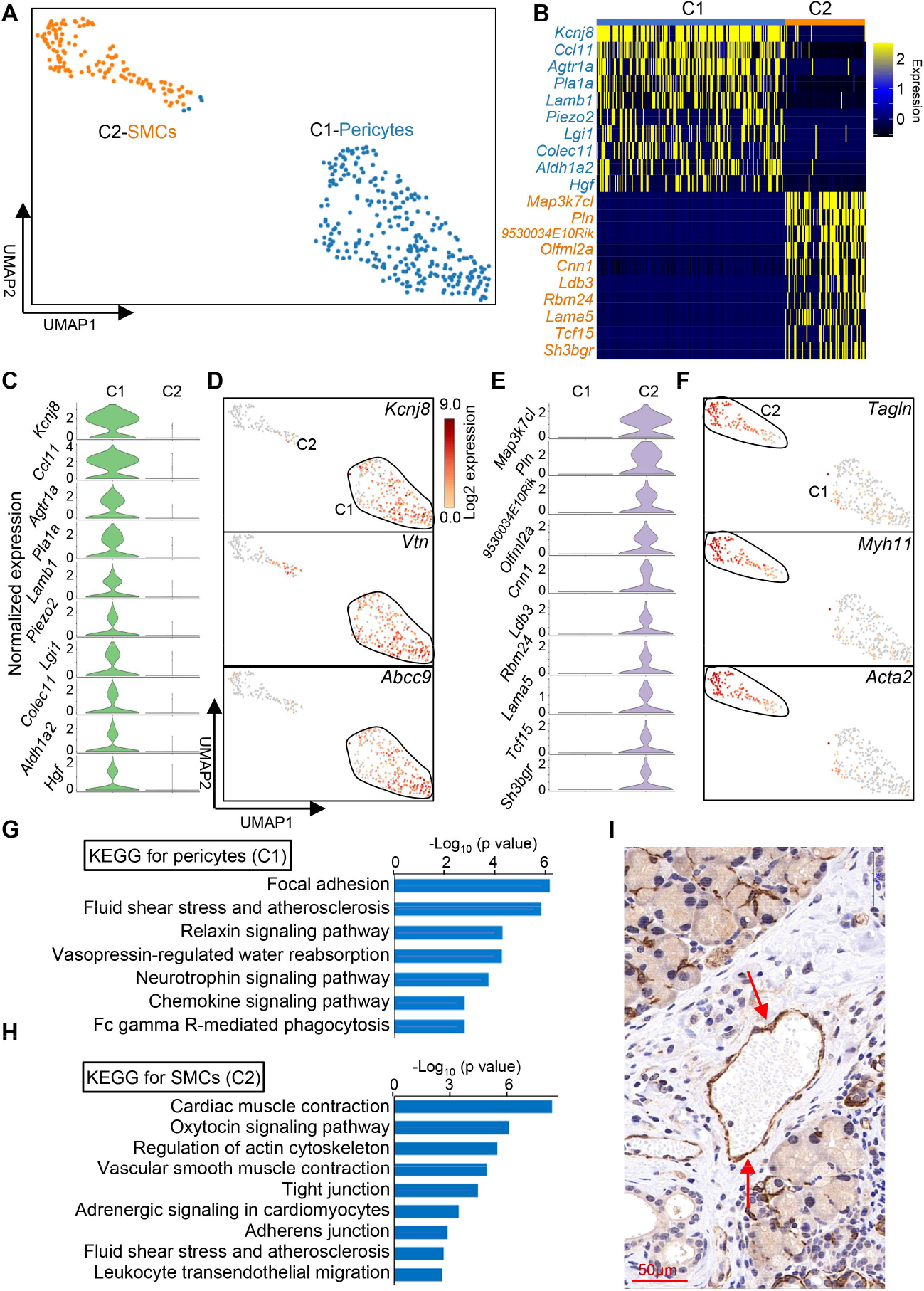
Subpopulation of vascular mural cells in the lacrimal gland. **(A)** UMAP visualization of vascular mural cell clusters in the lacrimal gland. **(B)** Heatmap of the top 10 marker genes for each cluster of vascular mural cells. Color scale: yellow, high expression; dark, low expression. **(C)** Violin plots of the top 10 marker genes of pericytes (C1) around the vessel wall in the ELG. **(D)** UMAP visualization of pericytes by *Kcnj8*, *Vtn*, and *Abcc9* in the lacrimal gland. Color scale: dark brown, high expression; yellow, low expression. **(E)** Violin plots of the top 10 marker genes of the SMCs (C2) of the vessel wall in the ELG. **(F)** UMAP visualization of SMCs by *Tagln*, *Myh11*, and *Acta2* in the lacrimal gland. **(G, H**) Functional KEGG analysis for pericytes (G) and SMCs (H). **(I)** Immunohistochemistry of the lacrimal gland showing vessel wall staining using the anti-acta2 antibody. Arrow shows stained vessel walls (20× magnification, scale bar= 50 μm). SMCs, smooth muscle cells.

### 3.6. Schwann cells in murine ELG

The nerve fibers innervating the lacrimal gland are enwrapped by Schwann cells [98]. Similarly, this study found a small number of Schwann cells in murine ELG at the scRNA-seq level. This cell subpopulation characteristically expressed pan-Schwann cell markers, including *Mbp, Plp1*, *Kcna1*, *Cnp,* and *Foxd3* (**Fig. 11A**). The Schwann cells were further divided into two clusters: subcluster 1 predominantly expressed *Ugt8a, Ncmap, Cldn19, Gjc3,* and *Gldn* (**Fig. 11B**), whereas subcluster 2 predominantly expressed *Gfra3, Fabp7,* and *Zfp536* (**Fig. 11C**). However, both clusters expressed myelin protein genes *Mbp* and *Plp1* (**Fig. 11A**). Therefore, we confirm that Schwann cells in murine ELG belong to myelinated Schwann cells [63]. Functional enrichment analysis of the DEGs expressed by these Schwann cells showed that their main functions were closely related to pathways associated with neural activity and axon guidance (**Fig. 11D**).

**Fig. 11.**
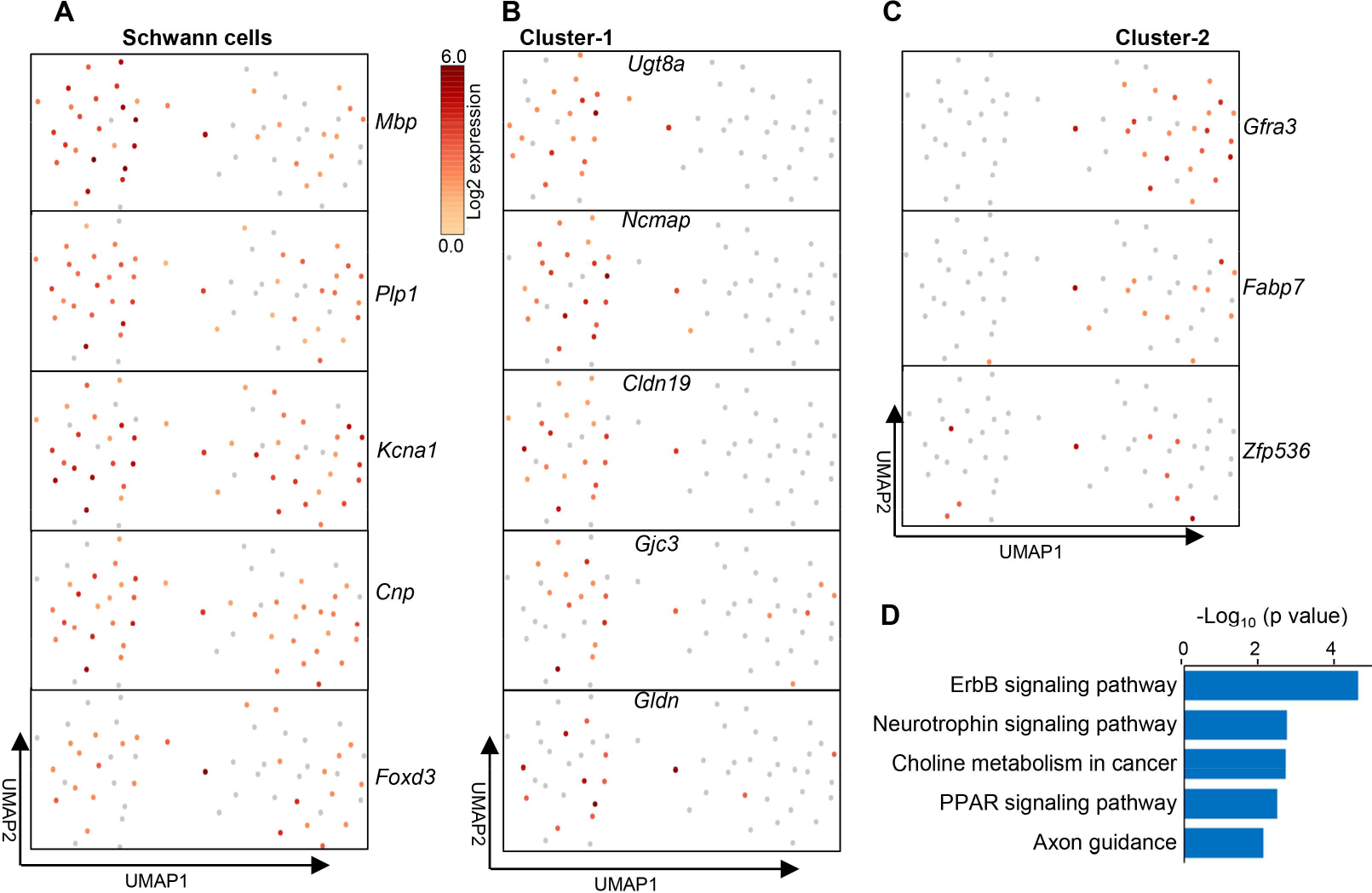
Schwann cell subpopulation in the lacrimal gland. **(A–C)** UMAP visualization of Schwann cells in the mouse ELG. Color scale: dark brown, high expression; yellow, low expression. **(D)** Functional KEGG analysis for DEGs expressed by Schwann cells in the mouse ELG.

### 3.7. Innate lymphoid cells in murine ELG

Innate lymphoid cells (ILCs) are non-T- and non-B heterogeneous innate immune cells [99]. They are mainly composed of NKs, ILC1s, ILC2s, ILC3a, and lymphoid tissue inducer cells based on the characteristics of surface molecules producing cytokines, and the dependence of development on transcription factors [99]. These cells reside in different peripheral tissues [100], including the cornea [101]. Four distinct ILC subsets (total of 4,996 cells) were defined in murine ELG according to the molecular characteristics of lineage marker negative (Lin^-^) (*Ptprc^+^, Cd3^-^, Cd19^-^, Ly75*^-^*, CD68^-^*) of this population [99, 102] (**Fig. 12A–F**).

**Fig. 12.**
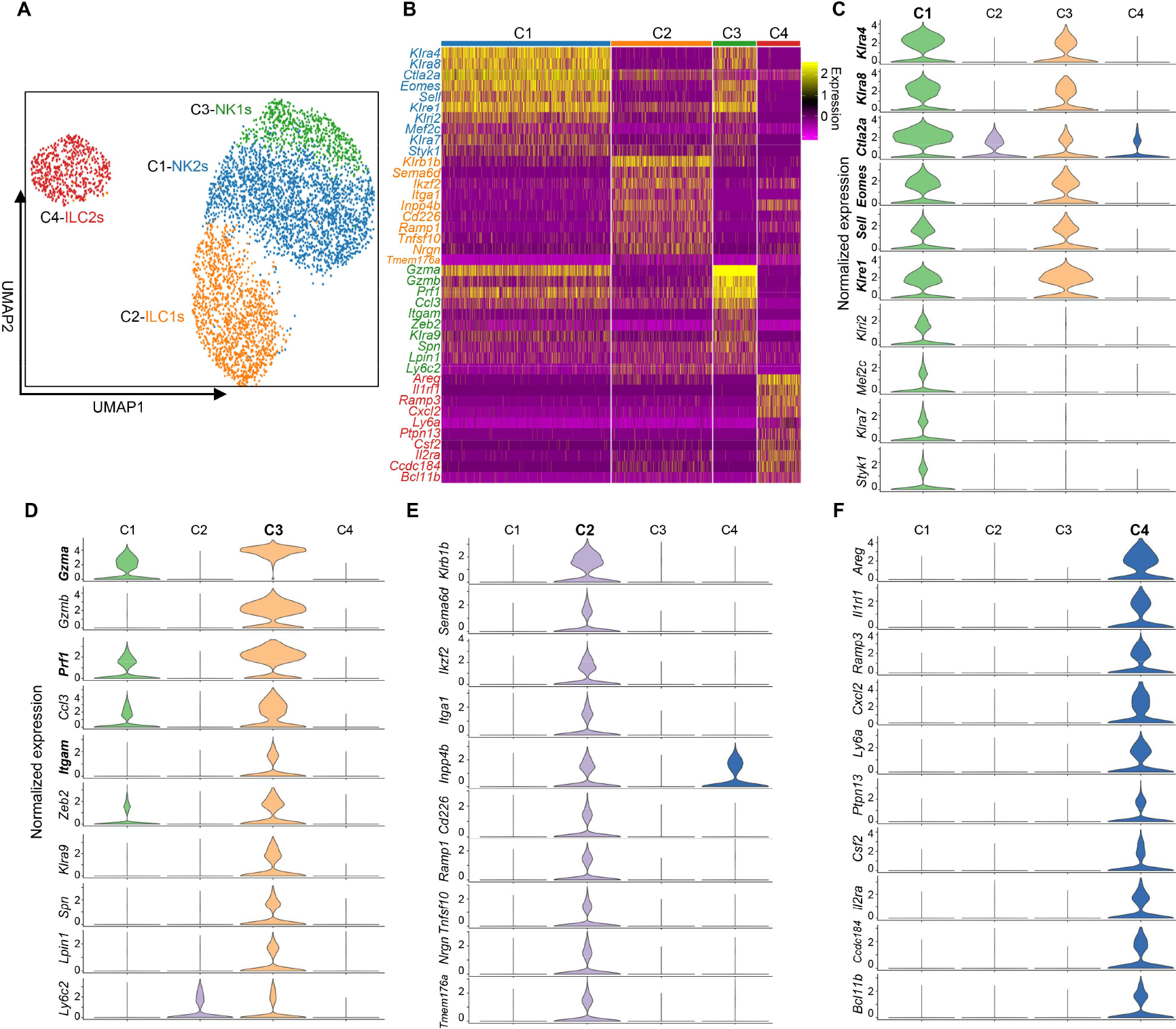
Subpopulation of innate lymphoid cells in the murine ELG. **(A)** UMAP visualization of innate lymphoid cell clusters in the murine ELG. **(B)** Heatmap of the top 10 marker genes for each cluster of innate lymphoid cells. Color scale: yellow for high expression; purple for low expression. **(C–F)** Violin diagram of the top 10 marker genes expressed by ILCs. Cluster 1 (C), Cluster 3 (D), Cluster 2 (E), Cluster 4 (F).

**Clusters 1** and **3** were considered classical NK cells because they characteristically expressed NK cell-specific marker genes, including *Klra4*, *Klra8, Gzma*, *Prf1* (encoding Perforin 1), *Eomes*, *Sell* (CD62L), *Klri2*, and *Klra7* (**Fig. 12C, D and Fig. 13A**) [103]. The NKs were further divided into two subgroups based on the predominance of *Itgam* (*CD11b*) and *Cd27* expression: *CD27*^-^*CD11b*^+^NK1 (12.19%) and *CD27*^+^*CD11b*^-^NK2 (47.48%), respectively [**Fig. 13B** (1,2,3)] [103]. Moreover, NK1s expressed higher levels of *Prf1*, *Gzma*, and *Gzmb* compared with NK2s [**Fig. 13B** (4, 5, 6)]. However, **Cluster 2** cells (28.22%) were referred to as **ILC1s** because this cluster specifically expressed genes unique to ILC1s such as *Il7r* (encoding CD127) [104–106], *Itga1* (encoding CD49a), and *Klrb1b* (**Fig. 14A, B**). Furthermore, this cluster co-expressed certain genes with NKs [107, 108], but lacked expression of NK-specific transcription factor (TF) *Eomes* [109, 110] (**Fig. 14A, B**). **Fig. 14C** displayed the functional signaling pathway enriched by the differentially expressed genes of ILC1s in the mouse lacrimal gland. Meanwhile, **Cluster 4** cells (12.11%) were defined as **ILC2s** because they express the gene for ligands of the EGF receptor ligand *Areg* (encoding amphiregulin) [111], *Gata3* (encoding gata-binding protein 3, the ILC2 development-dependent transcription factor) [111–113], *Il1rl1* (encoding IL-1 receptor-like 1, IL1RL1) [114], and *Cxcl2* (**Fig. 14D, E**). Finally, we displayed the functional pathways of DEGs for ILC2s (**Fig. 14F**). Collectively, mouse ELG harbors at least four different innate lymphoid cells at a steady state.

**Fig. 13.**
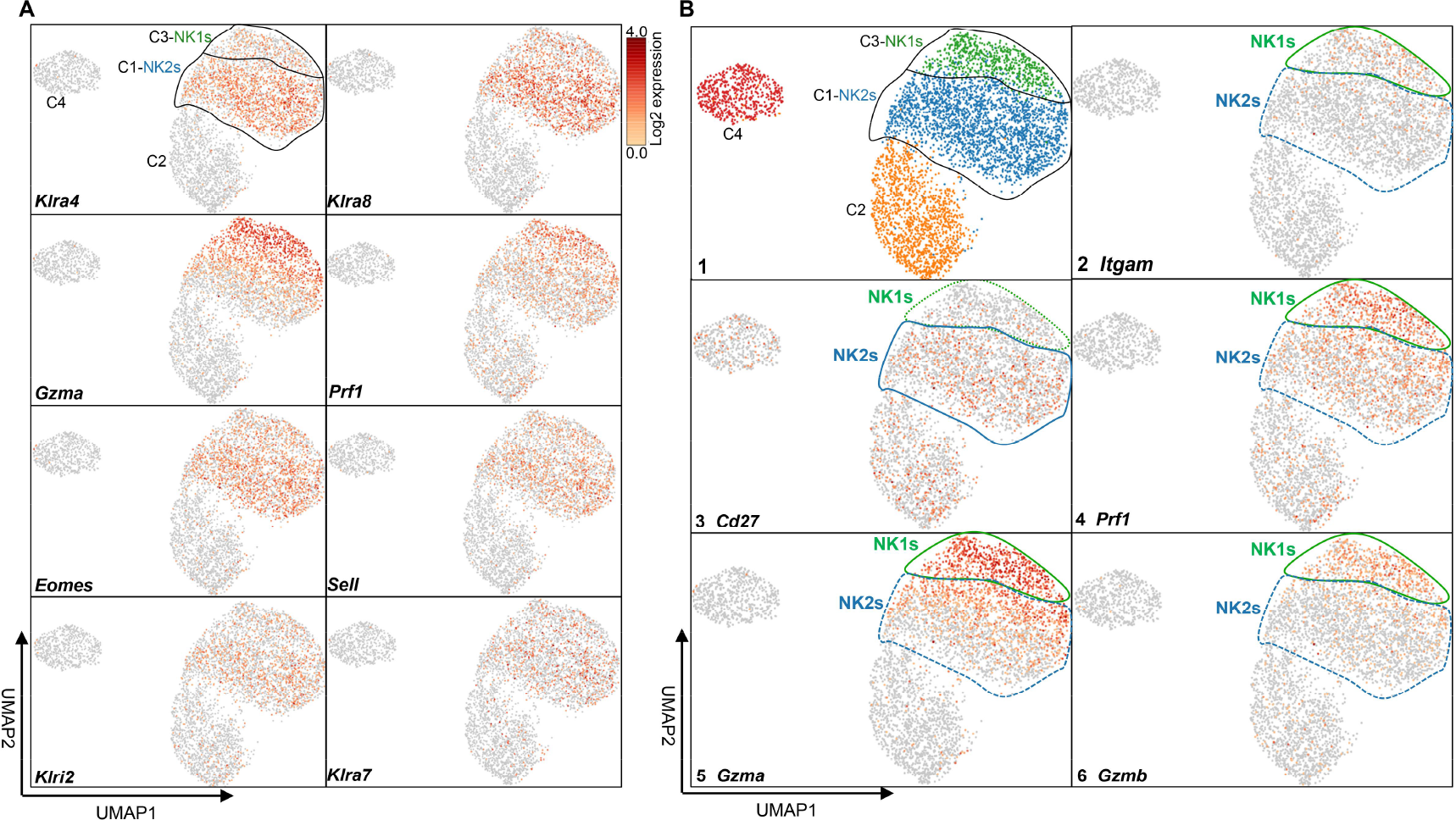
Comparison of NK1 cells and NK2 cells in murine ELG. **(A)** UMAP visualization of NK cells in mouse ELG. Color scale: dark brown, high expression; yellow, low expression. **(B)** Identification of NK1 cells and NK2 cells in mouse ELG. NK, natural killer.

**Fig. 14.**
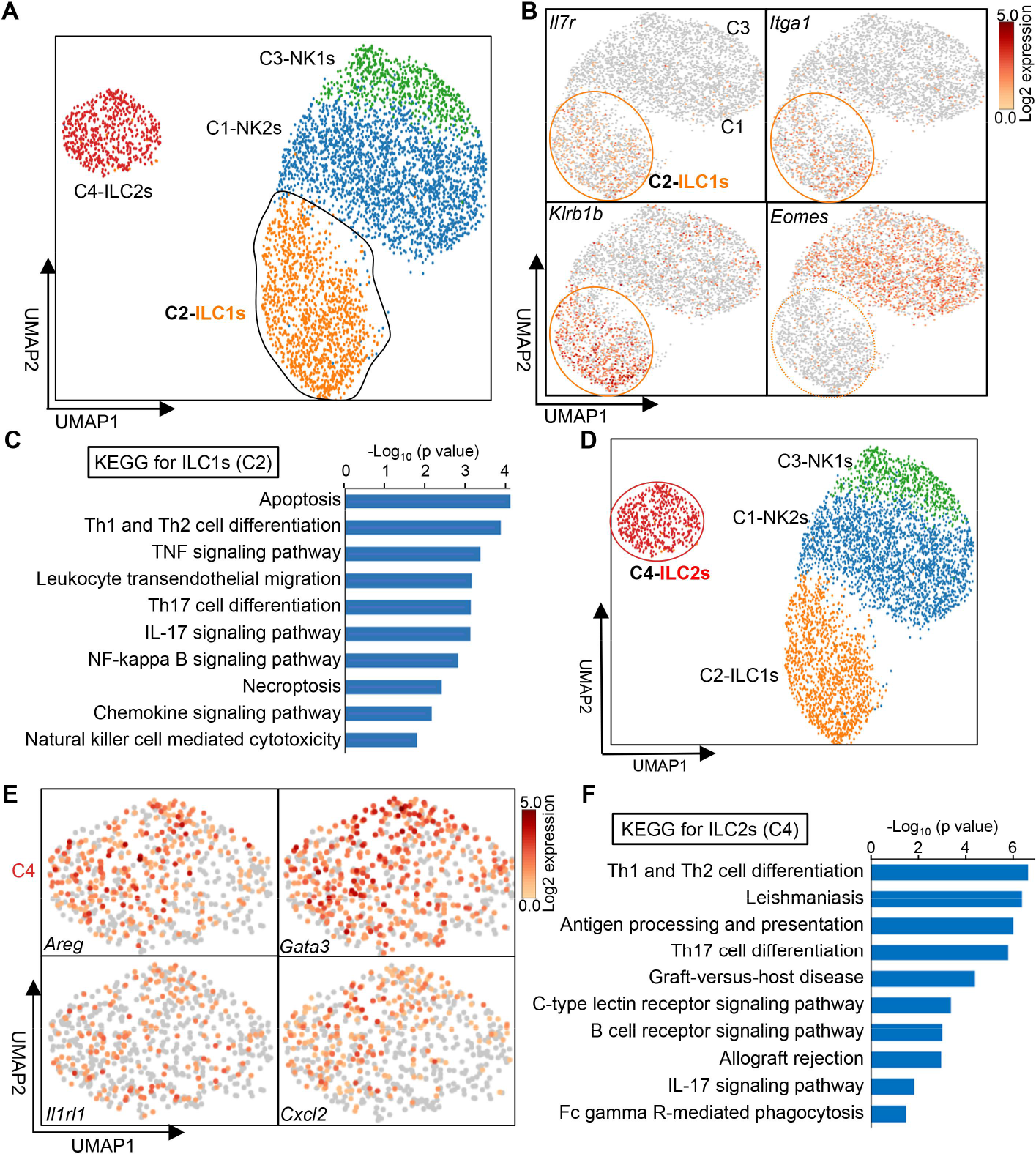
Identification of ILC1s (C2) and ILC2s (C4) in the murine ELG. **(A)** UMAP visualization of ILC1s (**C2**) in the murine ELG. **(B)** Classical marker gene expression of ILC1s in UMAP visualization. Color scale: dark brown, high expression; yellow, low expression. **(C)** Functional KEGG analysis of ILC1-expressed genes in the murine ELG. **(D)** UMAP visualization of ILC2s (**C4**) in the murine ELG. **(E)** Classical marker gene expression of ILC2s (**C4**) in UMAP visualization. **(F)** Functional KEGG analysis of ILC2-expressed genes in the murine ELG. ILC, innate lymphoid cell.

### 3.8. Myeloid cells in murine ELG

#### Macrophages

Macrophages are distributed in various glands [115]. The rat ELG also harbors some resident macrophages [24]. However, the diversity and transcriptional features of these macrophages in murine- and human lacrimal glands are unclear. Analysis of the macrophage population in mouse ELG was carried out at the scRNA-seq level. The cell population with macrophage characteristics clearly aggregated into 2 sub-clusters in UMAP coordinates by dimension reduction analysis (**Fig. 15A**). We named the two populations of cells (**Clusters 1–2**) as *Itgax*^hi^ (**Cluster 1**) and *Folr2*^hi^ macrophages (**Cluster 2**), respectively (**Fig. 15A–C**). The top 10 marker genes, percentages for each cluster of macrophages, and relative percentages of these 2 macrophage subsets are shown in **Fig. 15B, C,** and **D**, respectively. These 2 subset cell populations displayed high co-expression of *Apoe*, *C1qa* (encoding complement chains), *Lyz2*, cathepsin genes (*Ctsd, Ctsb,* and *Ctsz*), secretoglobin (Scgb) family (*Scgb2b7, Scgb1b3, Scgb2b2, Scgb1b7, Scgb2b24, Scgb2b20, Scgb1b20,* and *Scgb2b12*), *Cd74,* and high-level MHC II molecules (*H2-Aa, H2-Ab1, H2-D1, H2-Eb1,* and *H2-K1*) (**Fig. S6A**), while lacking expression of the classical dendritic cell chemokine receptor *CCR7* and transcription factor *Zbtb46* [116] (**Fig. S6B**).

**Fig. 15.**
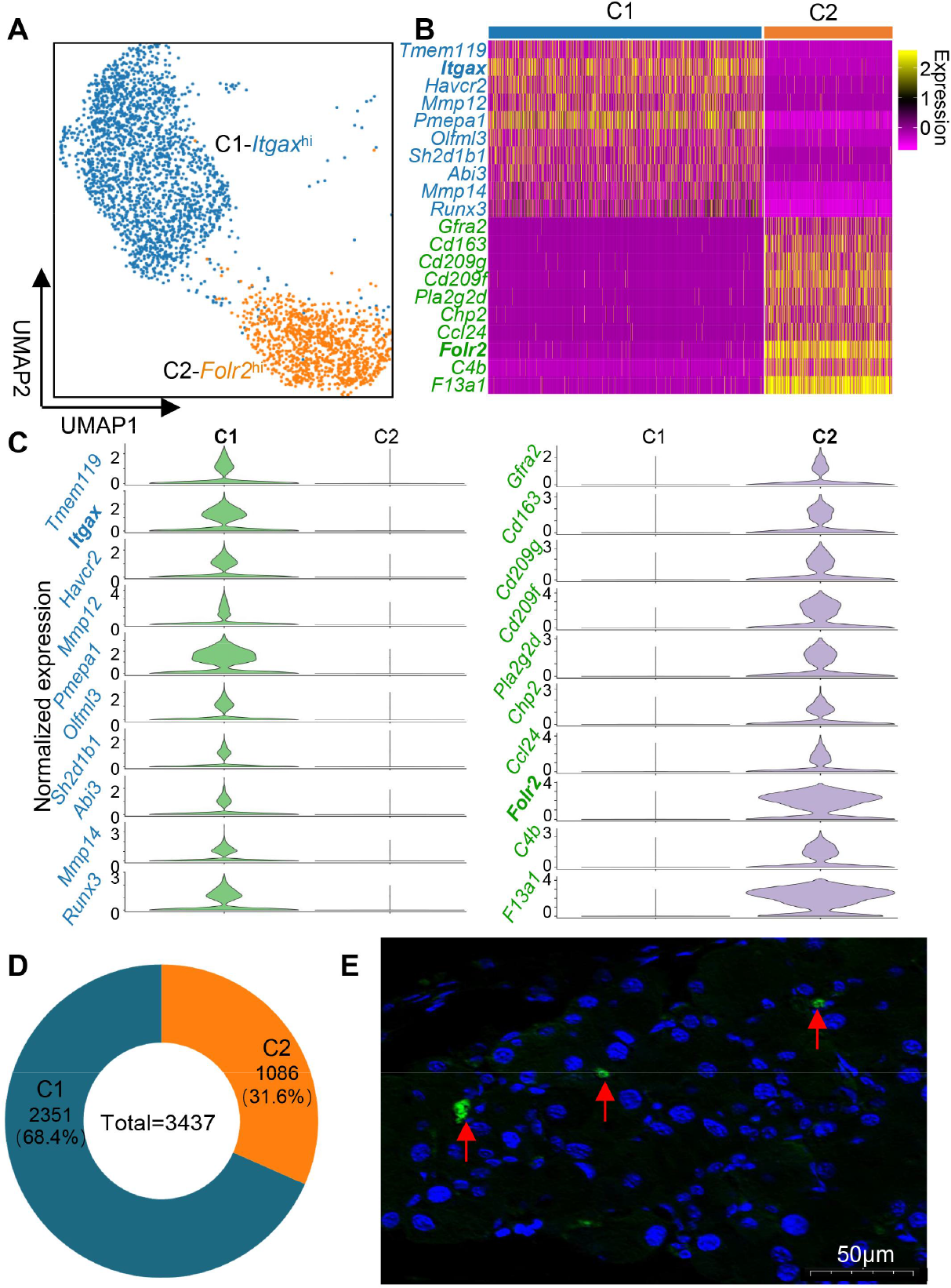
Subpopulation of macrophages in the lacrimal gland. **(A)** UMAP representation and unsupervised clustering of single cells within the macrophage cluster. **(B)** Gene-expression heatmap of the top 10 marker genes for each cluster of macrophages. Color scale: yellow, high expression; purple, low expression. **(C)** Violin diagram of the top 10 marker genes expressed by macrophages for Cluster 1 (*left*) and Cluster 2 (*right*). **(D)** Proportions of each macrophage cluster within the total number of macrophages. **(E)** Immunohistochemistry staining for macrophages in the lacrimal gland using the macrophage-specific marker, CD64. Arrows point to stained vessel walls (20x magnification).

*Itgax*^hi^ macrophages in **Cluster 1** uniquely express *Trem2* (encoding triggering receptor expressed on myeloid cells 2), *Il1b*, *Nlrp3, Cd9, Spp1* (encoding osteopontin), *Hvcn1, Cx3cr1* (encoding fractalkine receptor), and *Napsa* **(Fig. S6C)**, except for the top 10 genes mentioned above (**Fig. 15C**). Of note, *Folr2*^hi^ macrophages in **C2** are composed of *Timd4^+^*, *Lyve1^+^* (encoding hyaluronan receptor), and *Folr2^+^* (encoding folate receptor 2) (**Fig. S6D**). Thus, *Folr2*^hi^ macrophages were also defined as *TLF*^+^ macrophages as this signature is similar to a recently discovered macrophage signature in other major organs [117]. This cluster of cells also uniquely expressed *Mrc1* (encoding mannose receptor C-type 1/CD206)*, F13a1* (encoding coagulation factor XIII A subunit)*, Pf4* (encoding CXCL4), *Cd163*, *Cd209f*, and *Gas6* (**Fig. S6E**).

*Itgax*^hi^ macrophages in **C1** and *Folr2*^hi^ cells in **C2** co-expressed transcription factors *Mafb, Atf3,* and *Zeb2* (**Fig. S6A, *down***). Moreover, these two subsets largely overlapped with the expression of nuclear factor κB (NF-κB) inhibitors *Nfkbia* (encoding IκBα) and *Nfkbiz* (IκBζ), together with *Pltp* (encoding phospholipid transfer protein) [118, 119]*, Cd14, CD83, Cd86, Fcgr3*, *Adgre1* (encoding F4/80), *Csf1r*, and *Cd68* (**Fig. S6A**). Furthermore, functional KEGG enrichment analysis of the DEGs in the 2 cell populations revealed that the macrophage subsets had 2 distinct transcriptional signatures, similar to macrophages in other parts of the body [120, 121] (**Fig. S6F**). Immunofluorescent staining using the CD64 macrophage-specific marker [120] confirmed the macrophage identity in the ELG (**Fig. 15E**).

#### Monocyte/dendritic cells (Mon/DCs) in the murine ELG

The Mon/DCs in resting murine lacrimal glands were segregated into six unique clusters in UMAP coordinates based on gene expression similarity and reference datasets (**Fig. 16A, B and Fig. S7A)**; the proportion of different clusters is shown in **Fig. 16C**. All cells in these clusters broadly expressed MHC-II genes (*H2-Aa, H2-D1,* and *H2- K1*), *Cd74*, *S100a4*, *Ifi30*, *Napsa*, *Spi1 (PU.1)*, and *Klf4* (**Fig. S7B**). Cells of **Clusters 2** (24.90%) and **5** (5.53%) preferentially expressed genes associated with monocytes, including *Csf1r, Cebpb,* and *Mafb* (**Fig. S7C**), which are required for cell differentiation of hematopoietic stem cells into monocytes [122–126]. In addition, **Clusters 2** and **5** cells did not express *Zbtb46* (*Zbtb46* distinguished cDCs from monocytes and macrophages [123, 127, 128]) and *Flt3* (critical molecule for DC development [129, 130]) (**Fig. S7C**).

**Fig. 16.**
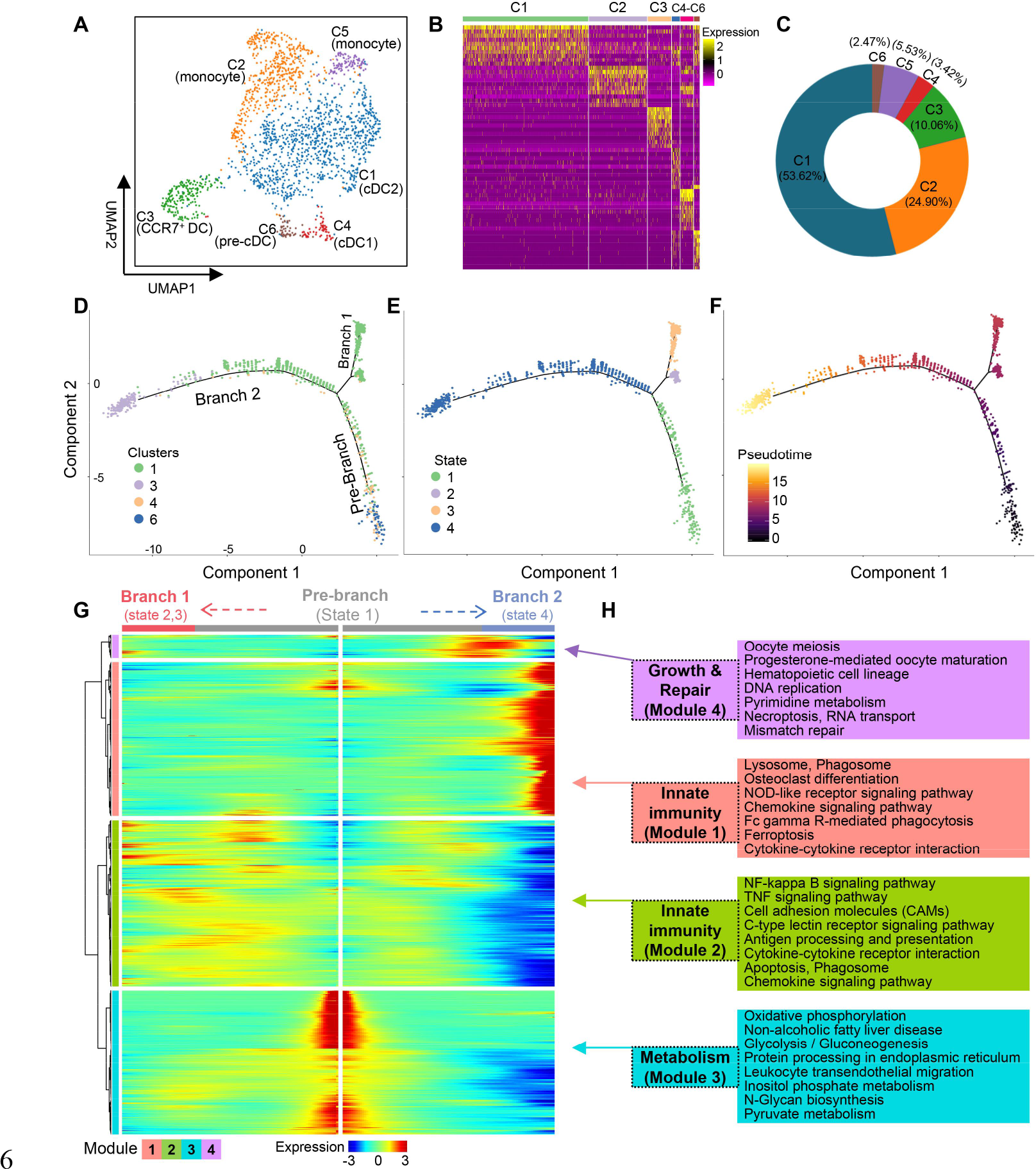
Subpopulation of monocyte/dendritic cells (Mon/DCs) in the lacrimal gland. **(A)** UMAP representation of Mon/DC clusters in the lacrimal gland. **(B)** Heatmap of the top 10 marker genes for each cluster of Mon/DCs. Color scale: yellow, high expression; purple, low expression. Gene list of the heatmap for each cluster was shown in **Table. S8**. **(C)** Proportion of each cluster within the total Mon/DCs. **(D)** Trajectory reconstruction of all single DCs based on cell clusters. Different colors represent different cell clusters revealing three branches: Pre-branch (before bifurcation), Branch 1, and Branch 2 (after bifurcation). **(E)** Trajectory reconstruction of all single DC clusters based on states. **(F)** Trajectory reconstruction of all single DC clusters based on pseudotime. Scale: darker colored dots (Pre-branch) represent earlier differential stage cells and lighter colored dots (Branches 1 and 2) represent later mature cells. **(G)** Kinetic heatmap generated using the BEAM regression model capturing the most DEGs between Branch 1 and Branch 2. Red and blue colors represent upregulation downregulation, respectively. **(H)** Top functional KEGG pathways from four different gene expression modules.

Cells in **Cluster 2** preferentially and highly expressed *Ly6c2* (encoding CD16), while **Cluster 5** cells did not express *Ly6c2* (**Fig. S7C**). Overall, this information indicates that cells in **Clusters 2** and **5** are monocytes that were excluded from further analysis.

Dendritic cells belong to another group of immune cell subset in the normal lacrimal gland [131, 132]. Under a steady state, circulating committed precursors of cDCs (pre-cDCs) arising from hematopoietic stem cells of bone marrow enter tissues by circulation and further differentiate into two main subsets, cDC1s and cDC2s through residual proliferative capacity [133]. Our analysis revealed that the cell subset of **Cluster 6** preferentially expressed *Mki67* gene coding for proliferation marker Ki-67 (**Fig. S7D**), which is consistent with the result of a previous study [133]. Of note, cells of this cluster uniquely expressed mitotic cell cycle machinery-related genes including *Kif11*, *Hist1h3c*, *Kif4*, and *Hist1h2ab* [**Fig. S7A** (**C6**)] [134, 135]. Altogether, this information indicates that **Cluster 6** represent pre-cDCs.

Further analysis showed that **Cluster 4** (3.42%) cell subset in the UMAP coordinate preferentially expressed *Xcr1* [136], *Irf8* [137], *Wdfy4* [138]*, Batf3* [139], and *Cd24a* [**Fig. S7E** (1–5)], but not cDC2-specific genes *Mgl2* [140], *Sirpa*, and *Cx3cr1* [141, 142] [**Fig. S7E** (6–8)]. This indicates that the cell subset in **Cluster 4** represents cDC1s cells [138, 143]. However, **Cluster 1** (53.62%) subset cells preferentially expressed *Clec10a* (*CD301a*) and *Cd209a*, indicating that these cells belong to cDC2s [140, 144, 145] (**Fig. S7F**). Notably, Cluster 3 cells (10.06%) were distinguished by high expression of *Ccr7* [144], *Id2*, and *Fscn1* [146, 147] (**Fig. S7G**). These cells represent *MHC-II*^hi^ *CCR7*^+^ migratory DCs correlated with the local lymph node, the preauricular node [22, 144, 148].

The Monocle 2 algorithm was used to order these DC single cells and construct the whole differentiation trajectory with a tree-like structure to explore a dynamic process for possible cell differentiation among these four cDC subsets (**Fig. 16D–F**). *Mki67*^hi^ cells in **Cluster 6** might be the differentiation initiation point, as this subset of cells uniquely expressed *Mki67* at high levels (**Fig. S7D**) and other mitotic cell cycle machinery-related genes above described [**Fig. S7A**(**C6**)]; these molecules are essential for pre-cDC colonization in peripheral tissues [133, 149]. A global differential analysis comparing Branches 1 (States 2 and 3) and 2 (State 4) revealed four different kinds of kinetic gene expression modules with branch-dependent expression (**Fig. 16G**). Genes from four different modules were significantly enriched in different pathways, mainly including innate immunity, metabolism, growth, and repair (**Fig. 16H**). These data indicate that the DCs distributed in mouse ELG are heterogeneous and play different roles.

#### Mast cells

Mast cells (MCs) are another unique immune cell population in the lacrimal gland [150, 151]. Our dimensionality reduction analysis of cells with mast cell-specific characteristics clearly showed that the mast cell population in the ELG was separated into two clusters (**Fig. 17A–C)**. Further analysis revealed that **Cluster 1** cells (83.90%) uniquely expressed *Mrgprx2* (encoding Mas-related G protein-coupled receptor member X2, MRGPRX2) [152], *Mrgprb2*, *Mcpt4*, and *Ncam1* [153, 154] (**Fig. 17D**), indicative of embryonic-derived connective tissue MCs. [155, 156], whereas **Cluster 2** cells (16.10%) specifically expressed *Mcpt1* (encoding mast cell protease-1, Mcpt1) and *Mcpt2* (**Fig. 17E**), indicative of BM-derived mucosal MCs. In addition, both cell clusters co-expressed *Cpa3*, *Gata2*, *Ccl2, Cma1*, and *Kit* (**Fig. 17F**). Genes expressed from two MC clusters were mostly enriched in neuronal-, vascular activity-, and immune response-related pathways (**Fig. 17G, H**).

**Fig. 17.**
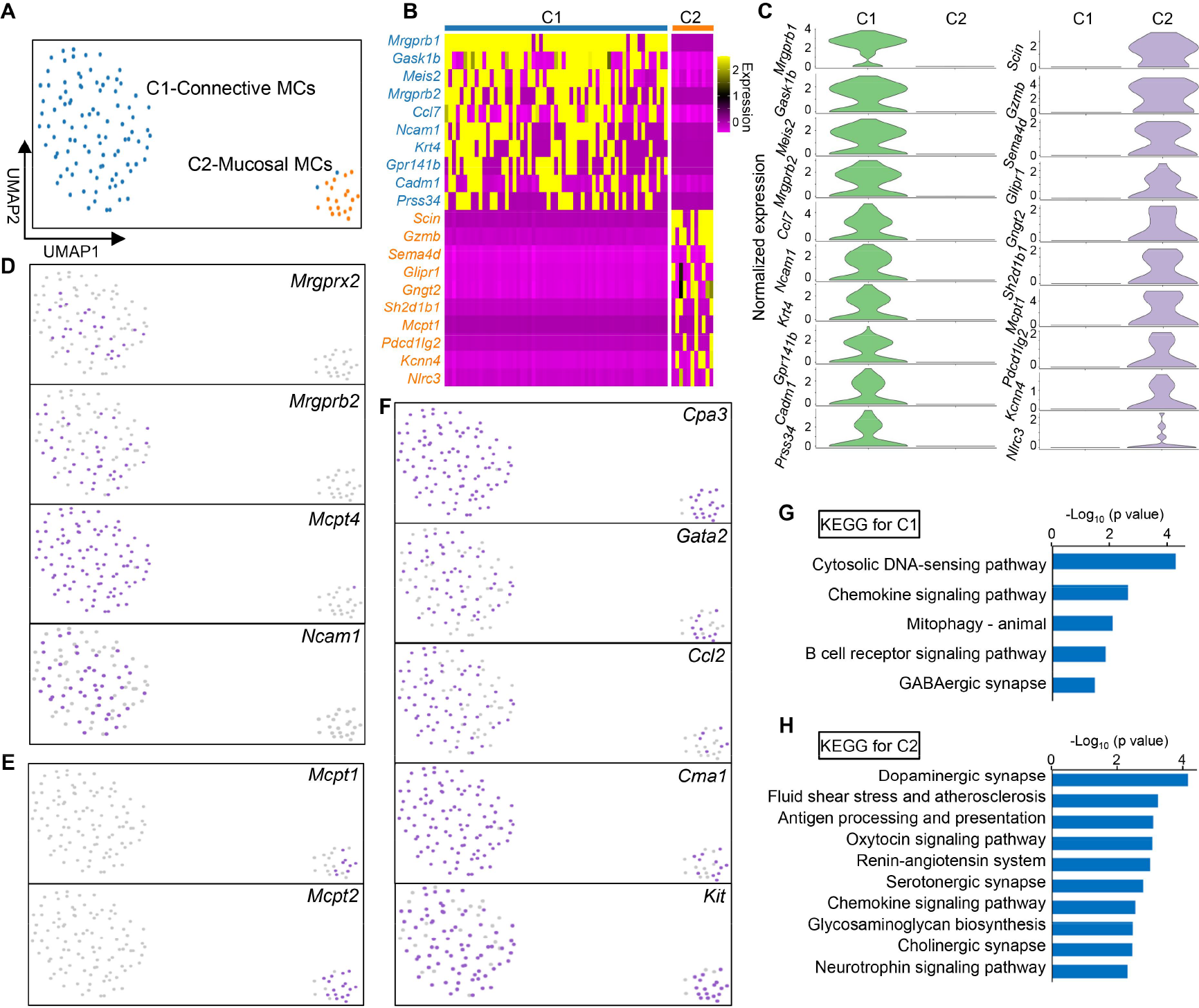
Subpopulation of mast cells in the murine ELG. **(A)** UMAP representation of mast cell clusters in the lacrimal gland. **(B,C)** Top 10 marker genes for each cluster of mast cells. Color scale: yellow, high expression; purple, low expression. **(D) Cluster 1** of mast cells specifically expresses *Mrgprx2*, *Mrgprb2*, *Mcpt4*, and *Ncam1*. **(E) Cluster 2** of mast cells specifically expresses *Mcpt1* and *Mcpt2*. **(F) Clusters 1** and **2** of mast cells express *Cpa3*, *Gata2*, *Ccl2*, *Cma1*, and *Kit*. **(G, H**) Functional KEGG analysis for Cluster 1 (G) and Cluster 2 (H) of two different mast cells.

### 3.9. T lymphocyte subsets in the ELG

Dimensional reduction analysis of all T lymphocyte populations (identified by *Cd3d*, *Cd3e*, and *Cd3g*) showed that four distinct clusters (**C1–C4**) are mainly present in resting lacrimal glands (**Fig. 18A, B**). The majority of cells in these clusters expressed effector-memory T cell-associated marker transcripts, including *Cd44*, *Itgb1, Igals1*, and *S100a*4 genes [157, 158], but not conventional naïve T cell marker genes such as S*ell* (encoding CD62L), *Ccr7*, *Dapl1,* and *Lrrn3* [159, 160] (**Fig. 18C**). These data indicate that the majority of T cell populations were effector-memory-like T cells in the resting murine ELG.

**Fig. 18.**
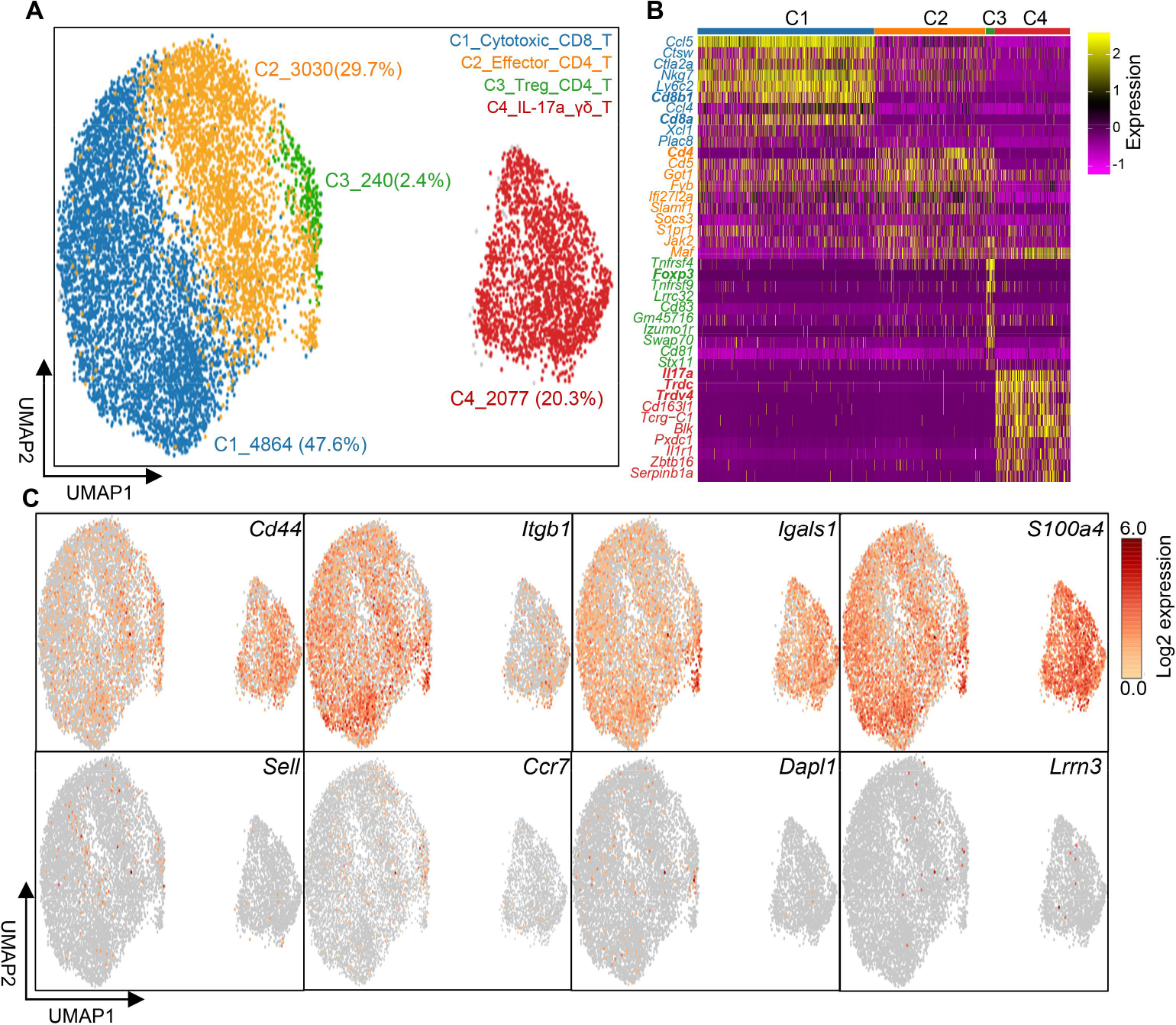
Subpopulation of T lymphocytes in the mouse ELG. **(A)** UMAP visualization of all T cell populations in the mouse ELG and the proportions of each subcluster within the total T cell population. **(B)** Top 10 gene-expression heatmap for each cluster of T cells. Color scale: yellow, high expression; purple, low expression. **(C)** Expressions associated with effector-memory and conventional naïve T cell marker genes under UMAP visualization. Color scale: dark brown, high expression; yellow, low expression.

#### CD4^+^ and CD8^+^ T lymphocytes in murine ELG

Further analysis demonstrated that the subset cells in **Cluster 1** were cytotoxic effector CD8^+^ T cells because most of these cells expressed characteristically cytotoxic effector transcripts closely linked with CD8 T cells, such as *Cd8a*, *Cd8b1*, *Nkg7* [161]*, Ctla2a* [162], *Gzmk*, *Prf1* (coding Perforin 1) [163], and *Eomes* [164] (**Fig. 19A, B**). Of note, this cell cluster highly expressed chemokines *Ccl5*, *Ccl4*, and *Xcl1*, which suggested a powerful effector cell differentiation (**Fig. 19B**) [165]. Finally, CD8^+^ T cells in the ELG stroma were verified by immunohistochemistry (**Fig. 19C**).

**Fig. 19.**
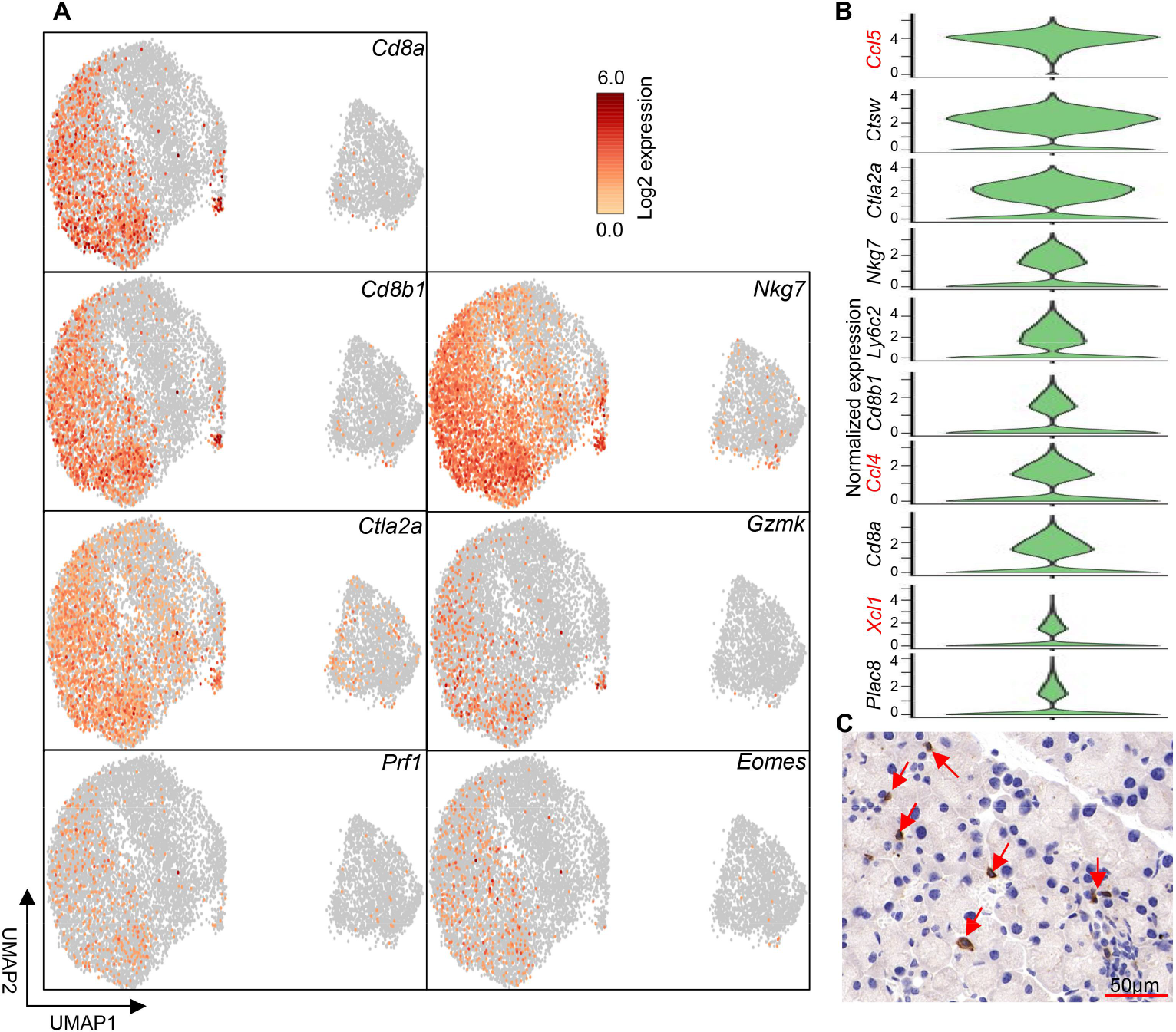
CD8+ cytotoxic T cell identity in the mouse ELG. **(A)** UMAP visualization and expression of classic genes of CD8^+^ T cells in the ELG. Color scale: dark brown, high expression; yellow, low expression. **(B)** Top 10 marker genes of CD8^+^ T cells in the ELG. **(C)** Histological immunofluorescence staining for the lacrimal gland showing CD8^+^ T cells using an anti-mouse CD8 antibody. Arrows point to positively stained cells (40× magnification).

Cells in **Cluster 2** were annotated as effector-memory CD4^+^ T cells since most of these cells express *CD4*^+^ and *Cd44*^+^, but do not express signature genes of naive T cells such as *Sell, Ccr7, Dapl1,* and *Lrrn3* (**Fig. 18C and Fig. 20A, B**). The cells in this cluster expressed the classical gene *Ifng* of T helper type 1 (Th1) cells, but did not express *Il4, Il13, Il9,* and *Il17a*, the signature genes of Th2-, Th9-, and Th17 cells, respectively (**Fig. 20A**). Therefore, these CD4^+^ T cells were classified as the Th1 subset, with the top ten marker genes shown in **Fig. 20B**. **Cluster 3** cells expressed classical marker genes of regulatory T cells (Tregs), including *Foxp3* (encoding transcription factor forkhead box P3) [166]*, Il10, Ikzf2* [167], *Ctla4* [168], *Gata3* [169], and TNF receptors (*Tnfrsf4, Tnfrsf9,* and *Tnfrsf18*) [170, 171] (**Fig. 20C**). **Fig. 20D and E** exhibited the top 10 marker genes and functional KEGG pathways of genes expressed by lacrimal niche-specific Tregs, respectively. Finally, the existence of CD4^+^ T cells was demonstrated by immunohistology (**Fig. 20F**).

**Fig. 20.**
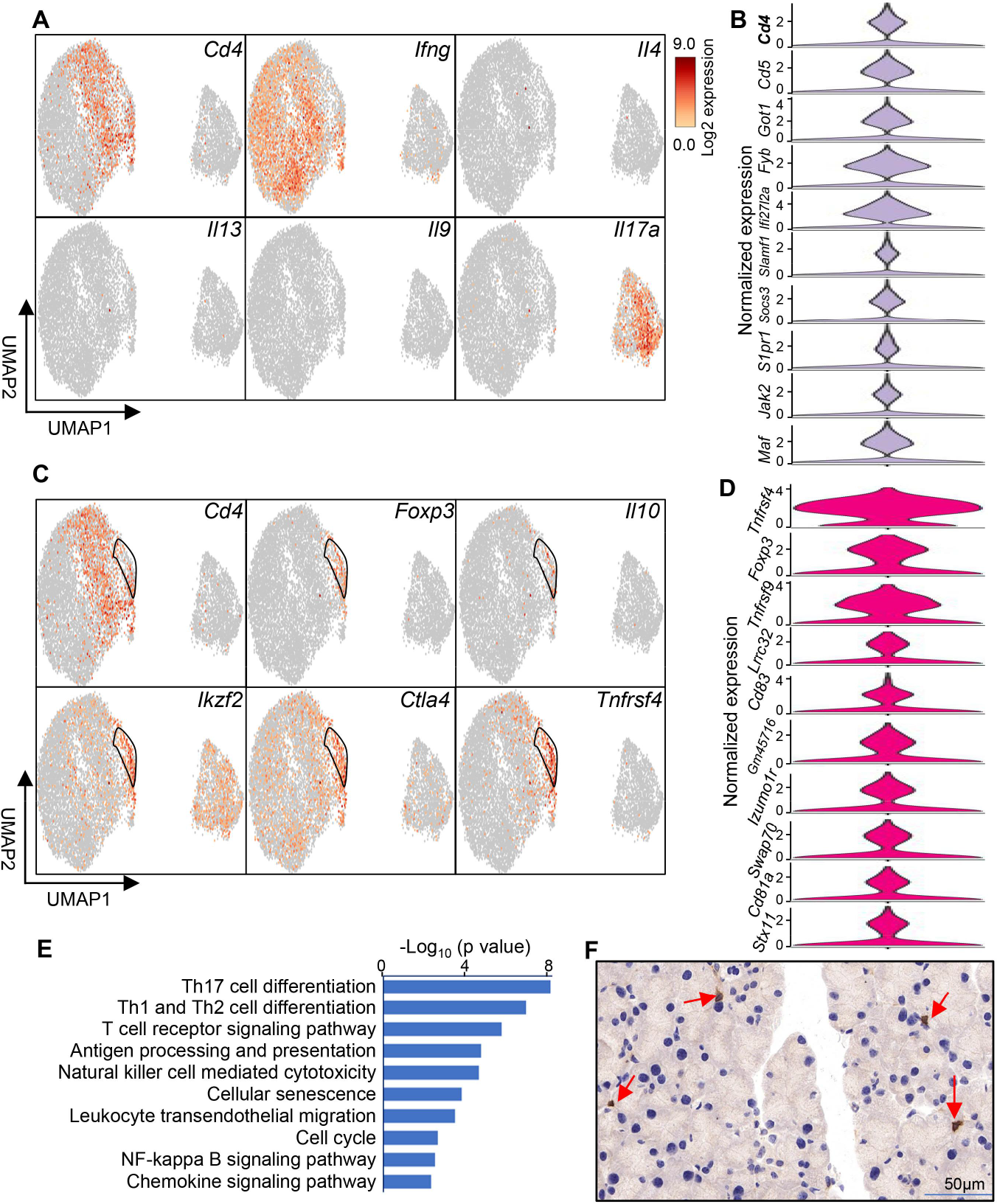
CD4^+^ T cell identity in the mouse ELG. **(A)** UMAP visualization and expression of classic genes of CD4^+^ T cells in the ELG. Color scale: dark brown, high expression; yellow, low expression. **(B)** Top 10 marker genes of CD4^+^ T cells in the ELG. **(C)** UMAP visualization and expression of classic genes of Foxp3^+^ Tregs in the ELG. **(D)** Top 10 marker genes of Foxp3^+^ Tregs in the ELG. **(E)** Functional KEGG analysis for DEGs in Foxp3^+^ Tregs. **(F)** Histological immunofluorescence staining for the lacrimal gland, showing CD4^+^ T cells using an anti-mouse CD4 antibody. Arrows point to positively stained cells (40× magnification).

#### γδ T-cells in murine ELG

γδ T-cells develop in the thymus and settle in peripheral tissues to provide physiological defense by rapidly releasing natural effector cytokines [172]. We further verified the presence of γδ T-cells in mouse ELG at the scRNA-seq level (**Fig. 18A, Cluster 4**; **Fig. 21A–E**) and using antigen-specific immunofluorescent staining (**Fig. 21F**), which is consistent with previous flow cytometry results [23]. Moreover, we confirmed that this subset of cells expressed *Trdc* (encoding T-cell receptor delta constant chain) and *Tcrg-C1* (encoding T-cell receptor gamma constant chain-1) (**Fig. 21A, E**). This subset of cells only expressed one of the variable region genes of T-cell receptor (TCR) gamma chain: *Trdv4*, and did not express *Trdv1*, *Trdv2-1*, *Trdv2-2*, *Trdv3*, or *Trdv5* (**Fig. 21A, E**). Importantly, this subset of cells highly expressed *Il17a* (**Fig. 21A, E**), weakly expressed *Ccr6* and *Cd27*, while *Il22* and *Ifng* mRNAs were barely detectable (**Fig. 21B, E**). These subset cells expressed four critical transcriptional factors for driving the differentiation of γδ T-cell progenitors toward mature γδ T-cells: *Sox13, Maf, Rorc,* and *Zbtb16* (zinc finger and BTB domain containing 16, or PLZF), which is consistent with findings of recent studies in mice [173, 174] (**Fig. 21C**). Furthermore, this subset of γδ T cells expressed B lymphoid kinase gene *Blk* (**Fig. 21D**) [175], which is critical for the development of IL-17- producing γδ T cells. We also readily detected *Cd163l1* (Scart1), which is important for conveying specific homing capabilities to γδT17 cells [176, 177], *Il1r1,* and *Serpinb1a,* which are required for homeostatic expansion of IL-17-producing γδ T cells (**Fig. 21D**) [178, 179]. Functional KEGG analysis suggests that the genes expressed by this γδ T-cell subset are mainly enriched in IL-17A-associated signaling pathways, Th cell differentiation, inflammatory response, and antigen presentation and processing pathways (**Fig. 21G**). Altogether, this information suggests that the resting mouse ELG predominantly harbors IL-17-producing Vγ4 γδ T cells.

**Fig. 21.**
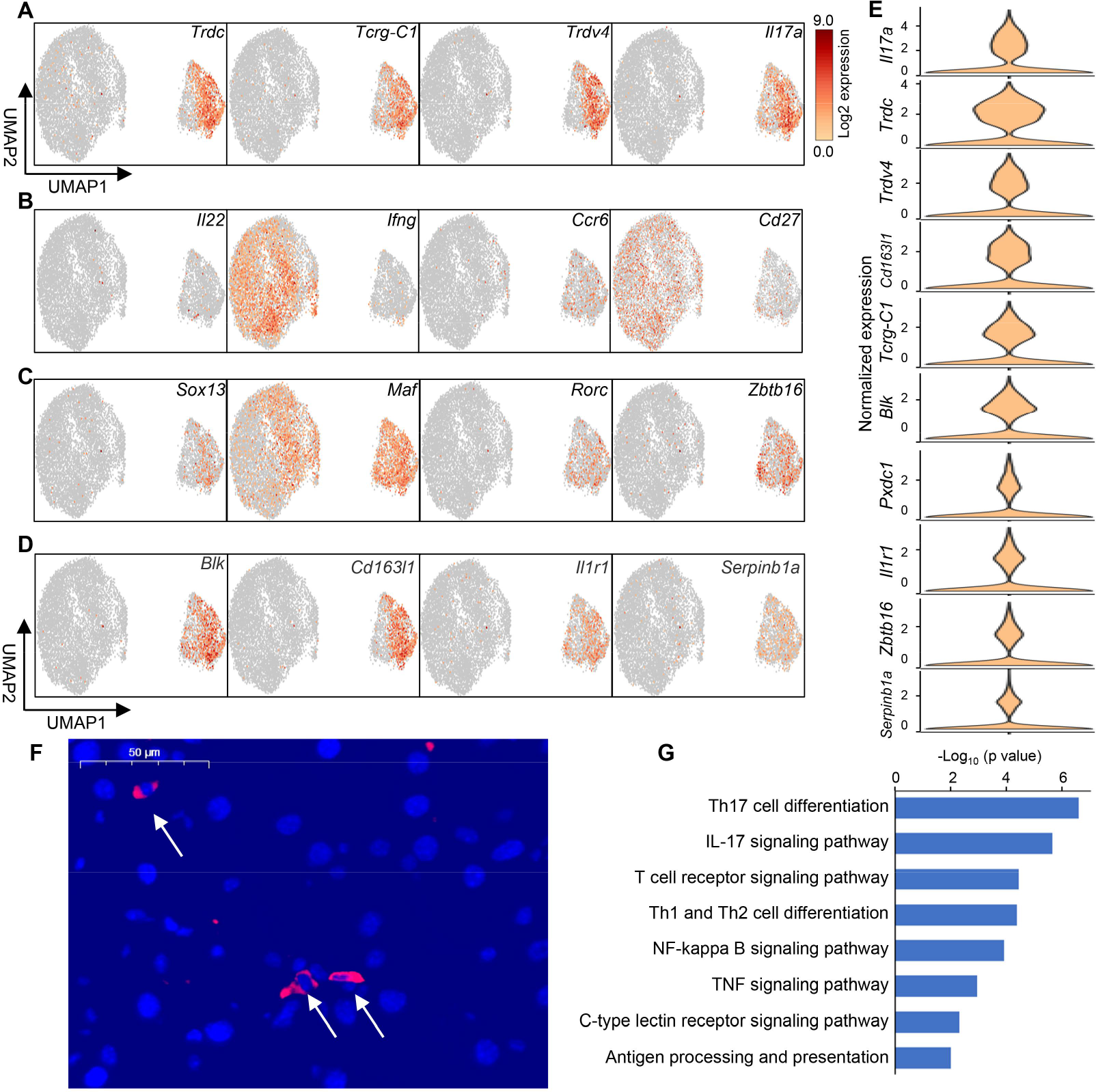
γδ T-cell subset Identity in the mouse ELG. **(A–D)** UMAP visualization of γδ T-cells in the mouse ELG. Color scale: dark brown, high expression; yellow, low expression. **(E)** Violin plot of the top 10 marker genes of γδ T-cells. **(F)** Histological immunofluorescence staining for the lacrimal gland, showing PE-conjugated anti-mouse TCRγ antibody (Gl3 clone, red) positive cells. Arrows point to positively stained cells. 40X magnification. **(G)** Functional KEGG analysis for lacrimal γδ T-cell-expressed DEGs.

### 3.10. B cells, plasma cells, and pDCs in murine ELG

B cells and plasma cells are another unique immune cell population in the lacrimal gland [22, 23, 180]. Dimensional reduction analysis showed that there are three cell populations with B cell-related characteristics in the ELG (**Fig. 22A, B**). Three B cell and B-like cell clusters co-expressed *Ly6a, Ly6d, Ly6e, Ptprc* (*B200*)*, Cd74, Tcf4, Klf2,* and *Ctss* [181–183] (**Fig. 22C**); *Ly6a, Ly6d, Ly6e,* and *Ptprc* (*B200*) are lymphoid-related genes.

**Fig. 22.**
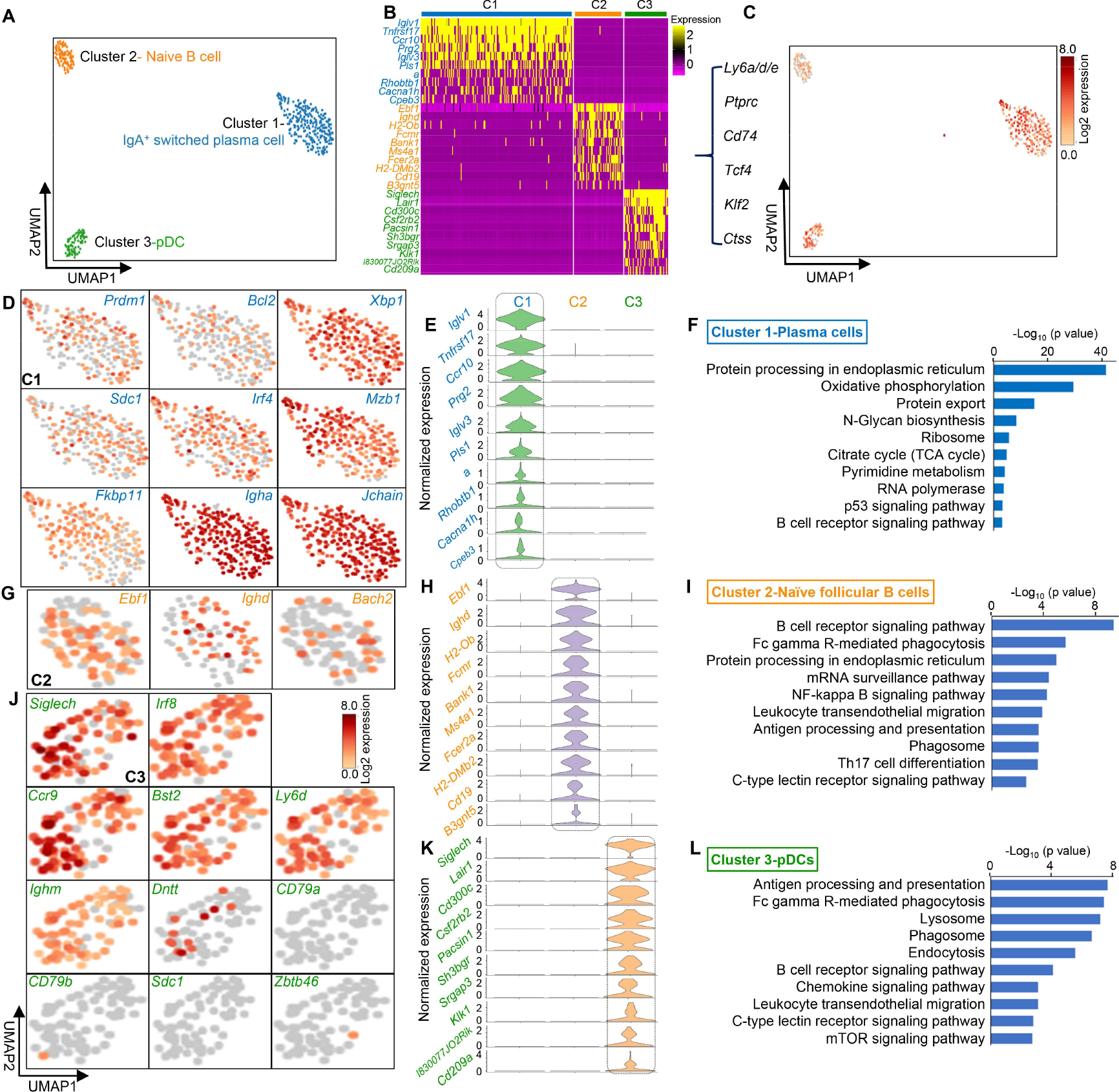
Identification-, cluster-specific marker genes-, and functional KEGG analysis of B cells, plasma cells, and pDCs in the murine ELG. **(A)** UMAP visualization of B cell-related clusters in the lacrimal gland. **(B)** Heatmap of the top 10 marker genes for each B cell-related cluster. Color scale: yellow, high expression; purple, low expression. **(C)** UMAP visualization of genes common to all three cell clusters. Color scale: dark brown, high expression; yellow, low expression. **(D)** Identification of **Cluster 1** of B cell-related cells using the classical plasma cell marker genes under UMAP visualization. **(E)** Violin plot of the top 10 cluster-specific marker genes of plasma cells (**Cluster 1**) in the ELG. **(F)** Functional KEGG analysis of plasma cells (**Cluster 1**) in the ELG. **(G) Cluster 2** of B cell-related cells identification using the classical naïve B cell-marker genes under UMAP visualization. **(H)** Violin plot of the top 10 cluster-specific marker genes of naïve B cells (**Cluster 2**) in the ELG. **(I)** Functional KEGG analysis of naïve B cells (**Cluster 2**) in the ELG. **(J) Cluster 3** of B cell-related cells identification using the classical pDC marker genes under UMAP visualization. **(K)** Violin plot of the top 10 cluster-specific marker genes of pDCs (**Cluster 3**) in the ELG. **(L)** Functional KEGG analysis of pDCs (**Cluster 3**) in the ELG. pDC, plasmacytoid dendritic cells.

**Cluster 1** expressed the genes of typical plasma cell specific protein-related genes, including *Prdm1* (PR domain containing 1), *Bcl2* (CD317) [184], *Xbp1* (X-box binding protein 1), *Iglv1* (immunoglobulin lambda variable 1), *Sdc1* (encoding CD138 or Syndecan 1) [185, 186], *Irf4, Mzb1, a*nd *Fkbp11* (**Fig. 22D**), and lacrimal niche-specific plasma cell marker genes (**Fig. 22E**). Additionally, it highly expressed *Igha* (encoding IgA) and *Jchain* (encoding joining (J) chain for covalent association with dimeric secretory IgA (SIgA) and polymeric IgM) (**Fig. 22D**). This indicated that murine ELG harbors IgA^+^- switched plasma cells similar to that of secondary lymphoid tissue [187, 188] and non-diseased meninges [189].

**Cluster 2** primarily expressed marker genes *Ebf1, Ighd* [190], *Bach2,* and other lacrimal niche-specific B cell marker genes (**Fig. 22G, H**), indicative of naïve follicular B cells [184]. These findings are consistent with the observation of IgA committed B-1 cells in murine ELG by flow cytometry [23] and plasma cells in human lacrimal gland stroma by immunohistochemistry [22].

Notably, **Cluster 3** cells highly expressed the plasmacytoid DC (pDC)-exclusive marker gene *Siglech* [191, 192], transcription factor *Irf8* for pDC development [182], *Ccr9*, a homing receptor of pDC to tissues such as the intestine [193, 194], *Bst2* [195, 196], *Ly6d* [181, 182], and lacrimal niche-specific pDC marker genes (*Lair1*, *Cd300c*, *Csf2rb2*, *Pacsin1*, *Sh3bgr*, *Srgap3*, and *Klk1*), but did not express B cell-specific and common marker *CD79a/b* and plasma cell-specific marker *Sdc1* (encoding CD138) (**Fig. 22J, K**). Thus, **Cluster 3** cell population represents pDCs. Furthermore, these pDCs also expressed *Ighm* and *TdT (Dntt)* (**Fig. 22J**). The *Ighm* gene encodes the C region of the μheavy chain to define the IgM isotype of naive B lymphocytes and *TdT (*encoding terminal deoxynucleotidyl transferase) inserts non-templated N-nucleotides into the junctions between gene segments in immunoglobulin V-region genes during their assembly [197]. Moreover, this pDC cluster did not express the recently identified DC specific-transcription factor *Zbtb46* (encoding zinc finger and BTB domain containing 46) [128] (**Fig. 22J**). This suggests that lacrimal pDCs are developmentally derived from the lymphoid progenitor lineage.

Enrichment analysis of the DEGs showed that the three different cells may perform different functions; plasma cells in **Cluster 1** were associated with B-cell signaling pathway activities (**Fig. 22F**), whereas naïve follicular B cells in **Cluster 2** and pDCs in **Cluster 3** were mostly associated with innate- and acquired immune responses (**Fig. 22I, L**), respectively.

### 3.11. Cell–cell communication network in murine ELG

We attempted to map ligand–receptor interactions using our scRNA-seq data to systematically assess the interaction between cell types in the lacrimal gland (**Fig. 23A**). The cell types were ranked according to the number of ligand–receptor interactions between cells as follows: macrophages, ILCs, pericytes, endothelial cells, monocyte/DCs, mast cells, T cells, fibroblasts, Schwann cells, epithelial cells, B cells, neutrophils, and plasma cells (**Fig. 23B**). In addition, the interactions between the various cell types in the lacrimal gland were predicted by ranking them according to the number of interactions between receptors and ligands. The number of interactions (receptor cells vs. ligand cells) was ranked by fibroblasts (receptor cells) vs. ILCs (ligand cells), followed by pericytes vs. ILCs, macrophages vs. macrophages, fibroblasts vs. pericytes, monocyte/DCs vs. macrophages, and fibroblasts vs. endothelial cells (**Fig. 23C**). Furthermore, a circle plot shows the communication between fibroblasts (receptor cells) and other types of cells based on ligand–receptor interactions (**Fig. 23D**). The dot plot in **Fig. 23E** shows the significant ligand–receptor pairs involved in the interactions between fibroblasts (receptor cells, red) and other cell types (ligand cells, blue). Fibroblasts expressed high levels of epidermal growth factor receptor (Egfr), while the corresponding ligands were widely expressed in other cell types, suggesting that these fibroblast interactions play key roles in maintaining the physiological function of the lacrimal gland (**Fig. 23E**). The communication between ILCs (ligand cells) and other cell types based on ligand–receptor interactions is shown in **Fig. 23F**. The interaction pairs between ILCs (ligand cells) and other lacrimal cells are illustrated based on receptor–ligand interactions and their expression levels (**Fig. 23G**). Notably, the high expression of Cd74/Ccr5 was detected in immune cells, including B cells, macrophages, monocyte/DCs, and plasma cells, and the corresponding ligands were also highly expressed in ILCs, indicating their important roles in maintaining the normal immune function of the lacrimal gland (**Fig. 23G**). Specifically, we evaluated the interactions of Icam1 and its ligands between the ILCs and other cell subsets (**Fig. 23G**). Overall, these results suggest that there is close communication between the various cells in the lacrimal gland, in which fibroblasts and ILCs play an important role.

**Fig. 23.**
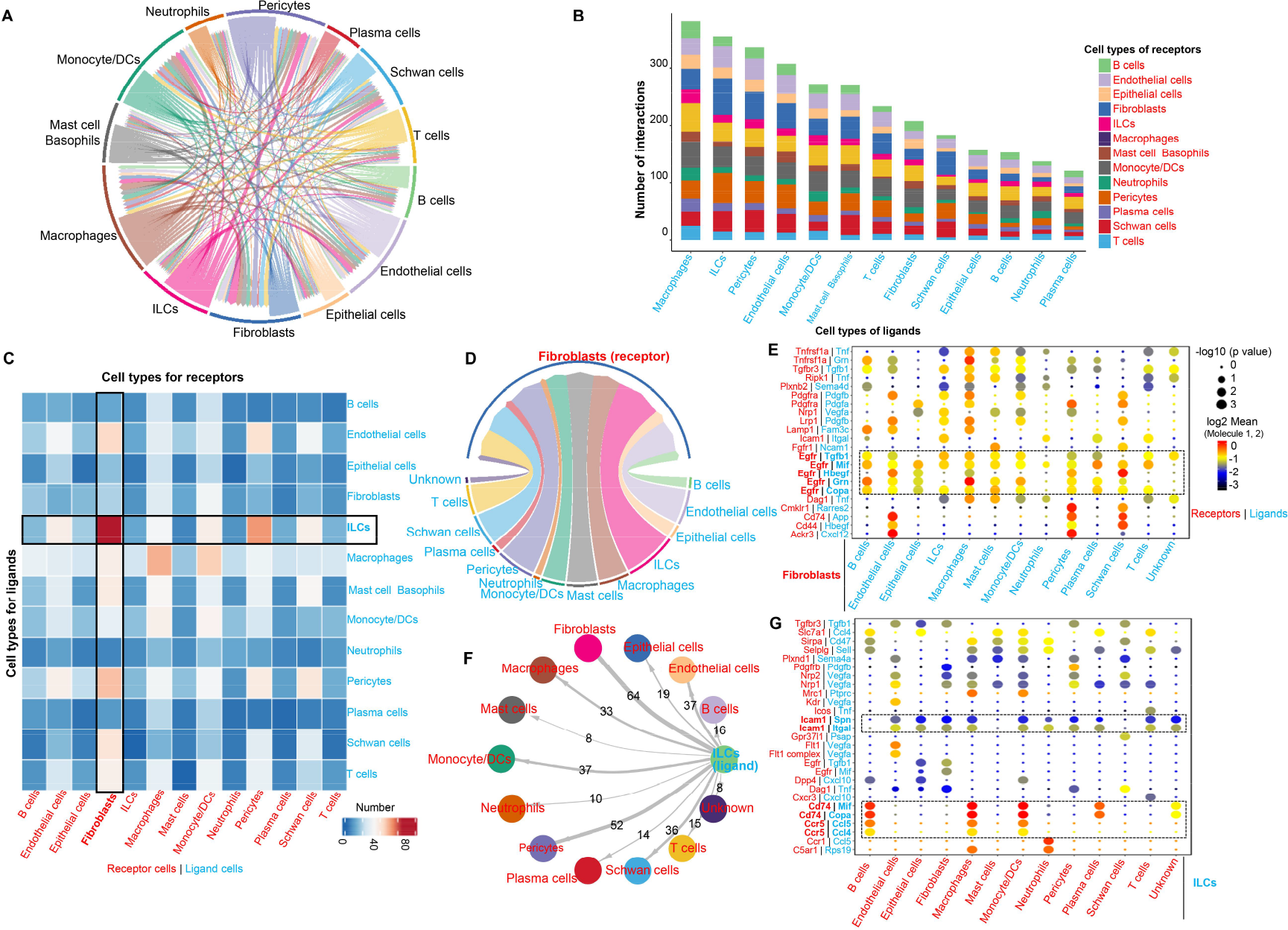
Interactions between various cell types in the murine ELG. **(A)** The number of intercellular ligand-receptor predictions for the indicated cell types based on scRNA-seq. Different cell populations were color-coded in the outermost circle and ligand-receptor pairs were linked with a solid line. The direction of the arrow indicates the interaction signal from the ligand cell to the receptor cell, and the thicker lines represent comparatively greater numbers of ligand-receptor interaction pairs. **(B)** Quantification of cell–cell communication frequencies in each cell type based on putative ligand–receptor interactions. **(C)** Heatmap showing the total number of interactions between cell types in the murine ELG. **(D)** Circle plot showing the communication of fibroblasts (receptor cells) with other cell types based on ligand-receptor interactions calculated by CellphoneDB. The direction of the arrow indicates the interaction signal from the ligand cell to the receptor cell, and the thicker lines represent comparatively greater numbers of ligand-receptor interaction pairs. **(E)** Dot plot showing the significant ligand-receptor pairs involved in the interaction between fibroblasts (receptor cells, red) and other cell types (ligand cells, blue). Receptor (fibroblasts) and ligand cell subsets are shown on the x-axis. Receptor (red) and ligand (blue) pairs are shown on the y-axis. **(F)** Circle plot showing the communication of ILCs (ligand cells) with other cell types based on ligand-receptor interactions calculated by CellphoneDB. The direction of the arrow indicates the interaction signal from the ligand cell to the receptor cell, and the thicker lines represent comparatively greater numbers of ligand-receptor interaction pairs. **(G)** Dot plot showing the significant ligand-receptor pairs involved in the interaction between ILCs (ligand cells, blue) and other cell types (receptor cells, red). Receptor and ligand cell (ILCs) subsets are shown on the x-axis. Receptor (red) and ligand (blue) pairs are shown on the y-axis.

## 4. Discussion

This work defines the transcriptomic landscape of adult mouse lacrimal gland cells and the heterogeneity of its various cells through unbiased scRNA-seq profiling of tens of thousands of murine ELG cells (**Fig. 24**). We identified classical and new cell types including ILC1s, ILC2s, “universal” stem-like fibroblast cell populations, and Schwann cells that were previously not of interest. Moreover, characteristics of *Cxcl17* and surface protein D-producing ductal cells were identified. In conclusion, these data provide a solid basis for understanding the detailed structure, function, and pathophysiology of lacrimal glands in the future.

**Fig. 24.**
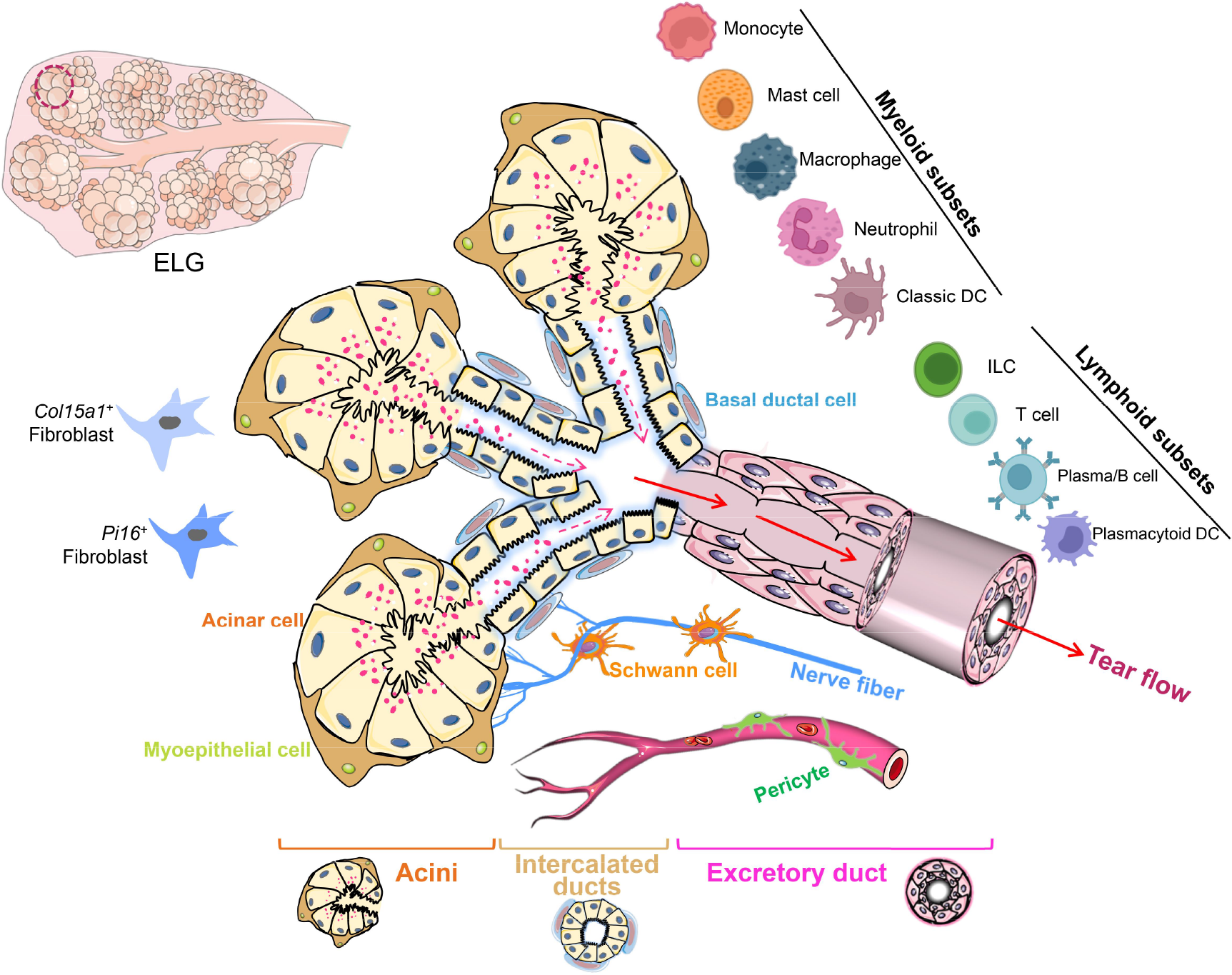
Schematic of murine ELG, the cellular neighborhood, and the identified cell types based on transcriptomics.

### 4.1. Glandular acinar epithelial cell population in murine ELG

Glandular acinar epithelial cells are main cells that secrete tears from the lacrimal gland [13, 14]. This study found that glandular acinar epithelial cells characteristically showed high expression of *Aqp5* and *Bhlha15* which correlated with recently studies of single cells in the murine and human lacrimal glands [11, 12]. In addition, our data confirmed and enriched other molecular signatures of murine lacrimal gland acinar cells at single-cell resolution [11, 12, 15]. Furthermore, we identified MECs surrounding the glandular epithelium which characteristically express marker genes associated with cell contraction such as *Acta2*, *Myl9*, *Mylk*, and *Myh11* in the lacrimal gland [19, 198, 199] and other exocrine glands [82, 83, 200, 201]. These cells also enable the synthesis of the basement membrane [19, 198, 199].

Additional exploration of the characteristics and functions of the following two cell types may be of particular and potentially important value: (1) Pheromone-related cell cluster: these cells preferentially express *Gm14743* and *Obp1b* subpopulations of cells.

*Gm14743* encodes a protein in the calycin superfamily and may be involved in binding lipophilic compounds to maintain the moist state of the ocular surface, while *Obp1b* encodes odorant-binding protein 1b which has unclarified pheromone activity by binding chemical odorants [202–204]. (2) Replicating cell cluster: these cells preferentially express mitotic cell cycle machinery-related genes and might be mostly associated with stem cells or stem cell-like features [205, 206]. However, these cell populations do not express the currently known marker genes of lacrimal stem cells by current scRNA-seq approaches [15, 207, 208]. Therefore, it may be important to further explore how this cell population maintains the integrity and homeostasis of the lacrimal glandular secretory epithelium.

Although there is some similarity in the top 10 expressed genes between the acinar cells in Cluster 1, 6, and 7, we considered there to be some differences between the three clusters for the following reasons: (1) The unbiased algorithm in the UMAP visualization map created a certain spatial distance between the three clusters of cells; (2) The top 10 marker genes of the three cell clusters do not overlap exactly; (3) KEGG and GO enrichment analysis of differentially expressed genes in Clusters 1, 6, and 7, respectively, showed that the potential functions of the three clusters of cells were somewhat different; and (4) The GSVA-BP and GSVA-KEGG results showed some differences in the GO-BP and the KEGG signaling pathways and its significance (FDR) enriched by the DEGs expressed among the three populations. In addition, another important reason for us to keep Clusters 1, 6, and 7 as separate clusters of acinar cells is related to several recent studies on the involvement of certain molecules secreted by human and rodent lacrimal glands in certain biological behaviors [202–204, 209]. We recognize that the secretory epithelium of the lacrimal gland has many unknown and intricate mechanisms that need to be explored. Therefore, we conservatively kept cell subsets separate rather than arbitrarily classifying them as the same class of cells.

However, the complexity of these mechanisms will require more research to validate in the future.

### 4.2. Glandular ductal cell population in murine ELG

The primary lacrimal fluid, secreted by acinar epithelial cells in the lacrimal gland, first enters the different segments of the ducts (intralobular, interlobular, intralobar, and interlober) [14]. The ductal cells of the lacrimal gland then form the final tear components by secretion or modification [16]. Detailed studies were performed regarding the lacrimal duct system histoarchitecture from humans, rabbits, rats, and mice [14, 18, 151, 210]; however, molecular characteristics at the single-cell level are unclear. In this study, the duct system cells of the murine lacrimal gland highly express ATPase at the single-cell level, which correlates with earlier studies [18, 80, 81]. There are three well-characterized cell populations at the single cell level based on differential gene expression: (**1) *Kit*^+^** intercalated ductal cells: this cell population generally connects the acini to the ductal system with enriched *Krt8* expression. In addition, this subset expressed stem/progenitor markers (*Kit* and *Cd14*), which might maintain ductal cell regeneration in the lacrimal gland [207] as reported in the salivary gland [82, 83, 200] and prostate regeneration [201].

Furthermore, this cell population highly expresses chemokine *Cxcl17* and two distinct anti-microbial molecule genes, *Sftpd* and *Ltf*. *Cxcl17* widely exists in mucosa (lung, epididymis, and uterus) and glands (thyroid, submaxillary, gland, oval, and prostate) [84, 211], which might function to recruit DCs and balance the immune stability of the ocular surface. **(2) *Sftpd*^+^** ductal cells: this population differs from *Kit*^+^ intercalated ductal cells and intermediately expresses *Sftpd* and *Ltf*, but not *Kit* and *Cd14*. This cell subset might represent larger secretory duct cells. Antimicrobial proteins SFTPD and LTF released from *Sftpd*^+^ and *Kit*^+^ ductal cell subsets might facilitate pathogen clearance on the ocular surface by inhibiting the growth of certain bacteria and regulating innate and adaptive immune-cell functions to provide a protective effect [212–215]. **(3) Basal ductal cells**: this cell population expresses typical basal ductal cell marker genes *Ly6a*, *Krt5*, *Krt14*, *Krt17*, and *Trp63*, but not MEC-specific genes including *Acta2*, *Myl9*, *Myh11*, and *Mylk*. This cell group generally surrounds the glandular duct, might undergo self-renewal, and produce luminal daughter cells by asymmetric division [11].

### 4.3. Fibroblasts in murine ELG

Fibroblasts are the basic structural cells within various organs, including the lacrimal gland [216, 217]. They secrete a matrix that provides a histological framework for vascular, neural, immune, and organ-specific structures within the organ [216, 218]. A series of recent studies showed that there is a large heterogeneous phenotype in different organs and tissues of mice and humans using scRNA-seq profiling [219–221]. These studies have shown that there is considerable heterogeneity in fibroblasts in phenotype, gene expression, and physiological function, which has expanded our prior knowledge. In particular, a recent study integrated single-cell transcriptome data to generate a map of fibroblasts in the steady state and the pathological states in 16 different tissues in mice and humans [88]. Two different groups of cells belong to “universal” fibroblasts in most organs or tissues. These are *Col15a1*+ parenchymal fibroblast cells and *Pi16*+ adventitial fibroblasts [88]. This study showed a baseline and specific fibroblast phenotype between species and pathology. Other fibroblast populations are mostly derived from the cellular differentiation of these two populations.

Similarly, we found at least 7 different fibroblast types in the lacrimal gland in the steady state based on scRNA-seq levels. Importantly, this study found two similar “universal” cells in the ELG which are highly expressed stemness-associated genes: *CD34* and *Ly6a* [90, 91]. In addition, the terminal differentiation process of these two stemness cell populations-, and the physiological processes and functions involving these cells was simulated in other cell populations by pseudotime analysis. However, further insight into the mechanisms and processes of differentiation between these cell populations and their physiological functions requires additional technical validation and in-depth analysis.

### 4.4. Vasculature-associated cell populations in murine ELG

The lacrimal gland is a broadly distributed exocrine organ within the vascular system [222]. Well-worked blood circulation is essential for oxygen- and nutrient delivery, immune-cell trafficking, and waste removal [223, 224]. Generally, blood vessels are composed of two cell types: endothelial cells (EC), arranged in the vascular lumen and mural cells surrounding and/or extending along the vessel walls. Mural cells include smooth muscle cells and pericytes. However, our understanding of lacrimal gland vasculature system is limited. This study defined and characterized the marker genes of the vascular-related cell types through transcriptomic profiling. This data highlights the pathophysiological mechanism of lacrimal gland vascular lesions under local and systemic diseases, such as angiopathy accompanied by diabetes [225]. However, this study did not capture lymphatic-associated cells.

The present study identified the presence of endothelial-to-mesenchymal transition (EndMT), a newly recognized type of cellular transdifferentiation, in adult mouse ELGs [226]. EndMT occurs not only during development but also persists after embryonic development, as endothelial cells expressing EndMT markers can be detected in the adult ovary and human valves [227]. Most of the current studies have focused on the association between the onset and development of cardiovascular disease [228]. In the field of ophthalmology, EndMT may also be involved in the development and progression of age-related macular degeneration [229]. However, further exploration is needed regarding the role of EndMT in lacrimal gland homeostasis and pathophysiology.

### 4.5. Lymphocyte and lymphoid subsets in murine ELG

This study confirmed that several different types of classical T lymphocytes exist in the mouse lacrimal gland, including effector CD4^+^ T cells, cytotoxic CD8^+^ T cells, Tregs, and γδ T-cells which is consistent with previous studies [23, 31, 230]. This study, for the first time, demonstrated the presence of type 1 and type 2 ILCs and NKs in the lacrimal gland. These immune cell subsets might be involved in the maintenance of lacrimal gland homeostasis through different immunological mechanisms.

**ILCs:** ILCs are a recently discovered group of diverse, plastic, and unique innate immune cells [99] exhibiting different phenotypes and functions depending on their localization [100, 231]. We found that at least 4 different ILC subsets reside in murine ELG. Although both NKs and ILC1s have cytotoxic activity [232], ILC1s might take an effect by producing Gzmc and NK cells by producing Gzma/Gzmb in the lacrimal gland.

Interestingly, NKs in the mouse lacrimal gland can also be further divided into NK1s and NK2s as in the spleen and peripheral blood of mice [103]. This is consistent with the higher cytotoxicity of human NK1s [233]. In addition, we found that ILC2s in the lacrimal glands specifically express epidermal growth factor receptor (EGFR) ligand *Areg* (encoding amphiregulin), which mediates epithelial repair following infection and injury [101, 111, 234]. Our further analysis of intercellular ligand and receptor interaction indicates the central role this group in maintaining normal physiology and structure.

However, the mechanisms of these ILC clusters in the pathophysiology of lacrimal gland disease require detailed studies.

**γδ T cells:**γδ T-cells are a major T cell subset of mucosal and peripheral tissues [235], including the ocular surface epithelium [236] and the lacrimal gland [23]. Recent research shows that γδ T-cells play an important role in immune surveillance by producing large amounts of cytokines to quickly trigger an inflammatory response, and play a key role in maintaining tissue integrity and repair after tissue injury by producing growth factors [235, 237–239]. Our single cell resolution found that γδ T-cells in the lacrimal gland are Vγ4 γδ T-cells, which seem to be similar to the mammary gland [240]. Vγ4 γδ T-cells develop in the late fetal thymus from embryonic day 17 until birth and throughout adulthood through TCR arrangement [173, 241–244]. Currently, γδ T-cells are divided into two main subsets [245]: IL-17-producing γδ T-cells associating with IL-17-mediated neutrophil recruitment during the early immune response [246–248] or IFN-γ-producing γδ T-cells linked to beneficial roles in cancer [249] and viral, parasitic and intracellular bacterial infections [250]. This study found that IL-17-producing γδ T-cells are the dominant subpopulation in the lacrimal gland. Collectively, this information provides a basic framework for further exploration of γδ T-cells in the physiological function of the lacrimal gland.

**Classic T lymphocytes**: This study identified two main types of CD4^+^ T cells in the lacrimal gland: Th1s and Tregs. Th1s may promote cell-mediated immune responses by secreting IFN-γ, IL-2, and TNF-α/β to defend against intracellular viral- and bacterial pathogens [251]. Meanwhile, Tregs play a key role in maintaining self-tolerance by actively downregulating immune activation [170, 171, 252, 253]. Current knowledge indicates that there are two subsets of Tregs: thymus derived Tregs (tTregs) and peripheral induced Treg cells (iTregs) [254–256]. tTregs are directly differentiated from the thymus, and their differentiation depends on the transcription factor Helios (*ikzf2*) [253, 257], while iTregs are derived from peripheral CD4^+^Foxp3^−^T cells, which are differentiated into T cells expressing *Foxp3*^+^ and acquire immunosuppressive functions in peripheral tissues [255, 258]. Tregs exist in the lacrimal gland and play an important inhibitory regulatory role to prevent autoimmune diseases [230, 259, 260]; however, their exact characteristics are unclear. This study found that Tregs in adult lacrimal glands highly express Helios (*ikzf2*), suggesting that Tregs in the lacrimal gland are mainly developed from the thymus under a steady state. However, further studies are required to determine the underlying mechanism of Treg evolution under pathological conditions, especially involving lacrimal gland inflammation accompanied by autoimmune diseases and aging [259, 260].

**B cells/plasma cells:** The majority of the body’s entire pool of activated B cells is located near the mucosae and the exocrine glands [261]. Human- and murine lacrimal glands harbor some B cells and plasma cells [22, 23]. Plasma cells might mainly produce IgA antibody, which is secreted by gland epithelial transmembrane transportation into the duct system and then into tear films [262]. IgA prevents bacterial and viral adhesion, and inactivates bacterial toxins, thus providing a barrier against potentially dangerous pathogens on the ocular surface [263]. This study found that plasma cells in the lacrimal gland predominantly produce IgA. Meanwhile, single-cell resolution of the murine lacrimal gland revealed that it contains two different kinds of B cell subsets: short-lived and long-lived plasma cells and naive follicular B cells, which correlated with previous findings [264]. The gene markers and transcriptomic profiling in this study might provide more insights to further understand B cells, the possibly associated lymphoid structure, and physiological function.

### 4.6. Myeloid cells in murine ELG

**Macrophages:** Macrophages are dispersed in all tissues of the body [265], including the lacrimal gland [14]. Their main functions include immune surveillance, phagocytosis of dead cell debris, and organ-specific tropic functions [266]. It is currently considered that macrophages residing in tissues are derived from two sources: (1) Yolk sac and fetal monocyte precursor-derived macrophages, with their maintenance occurring by self-renewal [267–269]. (2) Persistent differentiation of monocytes from bone marrow [270].

Their cell phenotype and transcriptomic profiling show tissue-specific behavior driven by unique local niches which evolve during development and aging [121, 271]. All these factors contribute to the heterogeneity and plasticity of resident macrophages in various organizations [272–274]. This study found that there are 2 subsets of macrophage populations in the lacrimal gland: *Itgax*^hi^ and *Folr2*^hi^. Notably, the *Itgax*^hi^ subset characteristically expressed high levels of matrix metalloproteinase (MMP) related to tissue remodeling and proinflammatory molecules. This subset of cells might be related to the triggering of lacrimal gland inflammatory responses and tissue remodeling after injury or other harmful events, similar to some macrophage subsets in the skin [275].

However, the *Folr2*^hi^ cell population (expressing *Timd4* and/or *Lyve1* and/or *Folr2*) highly expresses C-type lectin receptor CD209. CD209 recognizes and binds viral-, bacterial-, and fungal high-mannose type N-glycans with high affinity, thereby activating their phagocytosis [276]. Furthermore, this subset of macrophages highly expresses *Lyve-1*. Most of these cells are distributed around blood vessels to maintain tissue structure and hydration by binding the extracellular matrix (ECM) [277]. Depletion of *Lyve1*^+^ macrophages exacerbates experimental fibrosis [277], partly due to the production of collagen cleaving proteases [278]. Therefore, it is speculated that this subset of macrophages may be related to maintaining the physiological and structural integrity of the lacrimal gland. However, both *Itgax*^hi^ and *Folr2*^hi^cluster macrophages co-expressed transcription factors *Mafb* and *Atf3,* and some macrophage-associated transcripts such as *Cd14, Cd86, Ms4a7, Fcgr3,* and *Csf1r*.

**Mon/DCs:** Monocytes develop in the bone marrow and differentiate into two major populations present in the peripheral blood, classical monocytes and patrolling monocytes [279, 280]. In mice, classical monocytes express high levels of *Ly6C* and constitutively trafficked into peripheral tissues and differentiate into macrophages or DCs

[281]. However, when infection occurs in peripheral tissues these classical monocytes can rapidly mobilize to the site of injury and exert inflammatory or anti-inflammatory effects depending on the local microenvironment and the stage of injury [282, 283].

Patrolling monocytes are a smaller group of monocytes and roll along the vascular endothelium to examine endothelial injury, but they do not differentiate into tissue macrophages [284]. In mice, patrolling monocytes express *Ly6C* at low levels. The present study found that both types of cells are also present in the lacrimal gland. This suggests that two distinct subpopulations of monocytes could seed the resting lacrimal gland and might contribute to the maintenance of its homeostasis and respond to inflammatory stimuli from microbial infections and diverse injuries.

DCs are professional antigen presenting cells mainly distributed in barrier- and lymphoid tissues [285–287]. Currently, DCs are divided into three subgroups: conventional DC type 1 (cDC1), conventional DC type 2 (cDC2), and pDCs [288, 289]. Early studies found a small amount of DCs in the lacrimal gland [132, 290]. Recently, Ortiz and associate observed cell kinetics of cDC migration in the ELG of dry eye mouse models expressing CD11c enhanced yellow fluorescent protein using intravital multiphoton microscopy [132]. However, our data revealed that DCs in the lacrimal gland showed significant heterogeneity at single-cell resolution, with at least four subpopulations: cDC1s, cDC2s, *CCR7*^+^ migratory DCs, and pDCs. cDC2s were more abundant and heterogeneous in the ELG which is consistent with that observed in other locations [291]. The main function of the cDC2 subset is to efficiently capture exogenous antigens and induce CD4^+^ T cell responses [292], whereas cDC1s exclusively present antigens to naïve CD8^+^ T cells through a process called cross-presentation [143]. *CCR7*^+^ DCs migrate to locally draining lymph nodes (the preauricular node), then present antigens to Th cells and trigger an immune response [144, 293, 294]. Therefore, migratory *CCR7*^+^ DCs in the mouse ELG might function in a similar manner.

pDCs are a distinct lineage dedicated to the production of high amounts of type 1 interferons in response to viral infections [295–297]. However, our data revealed a small cell subset of pDCs localized in the B cell-associated cluster of the lacrimal glands after dimensionality reduction. Moreover, these pDCs co-express lymphoid-related genes with two other distinct B cells. This supports a recent hypothesis that pDCs have parallel development and differentiation from lymphoid precursors in bone marrow with B cells, not cDCs from myeloid-lineage progenitors [181, 182].

Conventional dendritic cell developmental pathways have been studied in most mouse and human tissues [144, 298]. However, cDC development in lacrimal glands remains poorly understood. Our observation suggests that cDCs in the lacrimal glands might differentiate from a specific cell cycle machinery gene cluster. This hypothesis is consistent with recent multicolor fate mapping performed on the developmental process of cDCs in peripheral tissues showing that cDC1s and cDC2s are derived from circulating precursors of cDCs (pre-cDCs) originating from bone marrow [133]. At a steady state, pre-cDCs entering the lacrimal gland have a residual proliferative capacity and differentiate into clones of sister cDCs, either cDC1s or cDC2s.

**Mast cells:** This study confirms the existence of mast cells in murine ELG at a scRNA-seq resolution which correlates with earlier studies [150, 151]. However, we found that both T cell-dependent mucosal- and connective tissue-types co-exist in the lacrimal gland. Mast cells are currently considered to be multifunctional innate immune cells [299], which trigger IgE-mediated allergic reactions, sense pathogens as innate immune cells, and regulate gland secretion by increasing blood flow into adrenal glands via histamine and serotonin production [300, 301]. Moreover, mast cells can release mast cell-specific tryptases, chymases, and growth factors involved in lacrimal gland growth and development by influencing the differentiation of epithelial cells and fibroblasts as seen in other glands [302, 303]. Finally, local mast cells might be linked to neurogenic inflammation in the lacrimal gland by producing MRGPRX2 as seen in other tissues [304, 305]. However, distribution of the two types of mast cells in the lacrimal glands and their corresponding physiological functions are worth further exploration.

### 4.7. Limitations of the study

**First**, the obtained dataset and cell clusters from mice cannot represent the total composition found in human lacrimal glands [14]. Therefore, caution must be used in interpreting human results. **Second**, this study only provided the cell composition and transcriptomic profile of male ELG. Humans and rodents have sex-dimorphism in cellular composition and gene expression of organs and tissues [306–308]. **Third**, only some cell phenotypes were verified by immunohistology due to the cost and availability of antibodies during the COVID-19 pandemic. **Fourth**, although this study found Schwann cells in lacrimal glands, neural cell populations from all samples were not captured. This limitation also exists in the scRNA-seq study of other glandular tissues such as the salivary gland [82]. Similarly, we did not detect the presence of lymphatic endothelial cells. Single nuclei isolation technology might solve this limitation [309]. **Fifth**, the lineage, evolution, and specific functions of some cell types, including glandular epithelial cells, fibroblasts, macrophages, and dendritic cells requires verification using fate mapping, proteomics, and genetic manipulation technology. **Finally**, further detailed localization of many cell types in lacrimal glands requires validation using spatial scRNA-seq approaches, especially some specific subclusters along the lacrimal duct tree and vessel tree. **Collectively**, database and other information from this study should be carefully interpreted and further validated.

## 5. Conclusions

Our findings reveal the diversity of lacrimal gland cell types and their transcriptomic profile. More importantly, this study provides a dataset as a resource, which deepens our understanding of the development-, composition-, and function of lacrimal glands and the occurrence of lacrimal gland diseases. However, more research is required to validate transcriptomic expression specificity and consistency, and the corresponding physiological function in different kinds of cells in the lacrimal gland.

## Statement of author contributions

Z.L. designed the study and wrote the manuscript. S.Z., X.J., and S.H. collected and prepared the samples. S.Z., Z.L., and X.J. performed scRNA-seq and bioinformatics analysis with help from S.H., J.L., H.S., D.Q., D.L., and X.P., Y.W. made some images and data analysis.

## Acknowledgements

This work was supported by the National Natural Science Foundation of China (82171014 and 81770962 to ZL, and 82101089 to SH), the Basic Science Project of Henan Eye Institute/Henan Eye Hospital (21JCZD001 to ZL, 21JCQN004 to SZ, and 20JCQN003 to SH), the Natural Science Foundation of Henan Province (232300421317 to SZ), the Doctoral Research and Development Foundation of Henan Provincial People’s Hospital (ZC20200229 to SZ, and ZC20190146 to SH), and the Henan Provincial Medical Science and Technology Research Joint Co-construction Project (LHGJ20210079 to SZ, and LHGJ20200064 to SH), the Key R&D and Promotion Special Program of Henan Province (212102311011 and 222102310120 to SH). The sequencing was conducted by OE Biotech Co., Ltd. (Shanghai, China).

## Declaration of competing interest

None of the authors have conflicts of interest to declare.

## Appendix A. Supplementary data

Supplementary data including **Fig. S1-7** and **Table. S1-8** to this article can be found online at https://XXX

## Supplementary Figure Legends

**Table. 1.** Antibodies used in this study.

| Cells | Antibody | Cat. |
| --- | --- | --- |
| MECs and SMCs | anti-alpha smooth muscle actin antibody | Abcam (USA, Cat #: ab5694) |
| B lymphocyte | CD79a monoclonal antibody | Thermo Fisher Scientific (USA, Cat #: MA5-16344) |
| CD4<sup>+</sup> T cells | anti-CD4 antibody | Servicebio (China, Cat #: GB13064-2) |
| CD8<sup>+</sup> T cells | anti-CD8 antibody | Servicebio (China, Cat #: GB13429) |
| γδ T-cells | PE anti-mouse γδ T-cell receptor | BD Biosciences (USA, clone GL3, Cat #: 553178 ) |
| Macrophages | FITC anti-mouse CD64 (FcγRI) antibody | BioLegend (USA, Cat #: 139316) |

Fig. S1. Violin plots of nGene, nUMI and percent.mito for each sample before (A–C) and after (D–F) quality control.

Fig. S2. UMAP visualization (A) and proportions (B) of all clusters for adult ELG cells from six donors by sample.

Fig. S3. Top 10 GSVA-BP and GSVA-KEGG in the glandular epithelial cells of the ELG.

**(A)** Top 10 GSVA-BP in each lacrimal glandular epithelial cell cluster.

**(B)** Top 10 GSVA-KEGG in each lacrimal glandular epithelial cell cluster.

BP, biological process; ELG, extraorbital lacimar gland; GSVA, gene set variation analysis; KEGG, Kyoto Encyclopedia of Genes and Genomes.

Fig. S4. Top 10 marker genes and annotations of Clusters 1, 6, and 7 in the

**glandular acinar cells of the ELG.**

**(A)** Top 10 marker genes of **Clusters 1, 6,** and **7** of lacrimal glandular epithelial cells.

**(B, C)** Top functional enrichment of KEGG (B) and GO (C) for genes expressed in

**Clusters 1, 6,** and **7** of lacrimal glandular epithelial cells.

GO, Gene ontology.

Fig. S5. Top 10 marker genes, BP, and KEGG of each cluster for seven clusters of fibroblasts in the mouse ELG.

**(A)** Top 10 marker genes of each cluster for seven clusters of fibroblasts in the mouse ELG.

**(B)** Top 5 BP of seven fibroblast clusters in the mouse ELG from the gene set variation analysis (GSVA).

**(C)** Top 5 functional KEGGs of seven fibroblast clusters of in the mouse ELG from the GSVA.

Fig. S6. Characterization of macrophages in the murine ELG.

**(A)** UMAP visualization of genes by the two macrophage clusters. Color scale: dark brown, high expression; yellow, low expression.

**(B)** The two macrophage clusters do not express classical dendritic cell chemokine receptor *CCR7* and transcription factor *Zbtb46*.

**(C)** Uniquely expressed genes in **Cluster 1** *Itgax*^hi^ macrophages.

**(D)** *Timd4*, *Lyve1*, and *Folr2* are uniquely expressed genes **in Cluster 2** macrophages.

**(E)** Uniquely expressed genes in **Cluster 2** *Folr2*^hi^ macrophages.

**(F)** Functional KEGG analysis for two different macrophage subsets, respectively.

Fig. S7. Characterization of Mon/DCs in the mouse ELG.

**(A)** Top 10 marker genes expressed by the six clusters of Mon/DC populations in the mouse ELG.

**(B)** Genes with relatively high co-expression in six Mon/DCs population clusters in the mouse ELG. Color scale: dark brown, high expression; yellow, low expression.

**(C)** Identification of monocyte signatures in the mouse ELG.

**(D)** *Mki67*^+^ committed precursor cDCs (pre-cDCs) in the mouse ELG.

**(E)** Identification of cDC1 (Cluster 4) signatures in the mouse ELG.

**(F)** Identification of cDC2 (Cluster1) signatures in the mouse ELG.

**(G)** *CCR7*^+^ migratory DC subset cells (Cluster3) in the mouse ELG.

**Supplementary Tables**

Table. S1. **All marker genes for each cluster of the six integrated samples (generated by the FindAllMarkers function in Seurat)**

**Table. S2. All marker genes for each cluster of each sample (generated by the FindAllMarkers function in Seurat)**

Table. S3. **Number and frequency (n [%]) of cells in each cluster for each sample Table. S4. The differentially expressed marker genes in Fig. 1E for each cell type of the six integrated samples.**

Table. S5. Top 10 marker genes for each cluster of **Fig. 1C**. Table. S6. Top 10 marker genes for each cluster of **Fig. 2B**. Table. S7. Top 10 marker genes for each cluster of **Fig. 7B**. Table. S8. Top 10 marker genes for each cluster of **Fig. 16B**.

## Notes

### Competing Interest Statement

The authors have declared no competing interest.

