## Supplementary material for "Analysis of the heterogeneity and complexity of murine extraorbital lacrimal gland via single-cell RNA sequencing": Table. S3

**Table. S3. Number and frequency [n(%)] of cells in each cluster for each sample.**

| **Cluster** | **Sample_1** | **Sample_2** | **Sample_3** | **Sample_4** | **Sample_5** | **Sample_6** | **All sample** |
| --- | --- | --- | --- | --- | --- | --- | --- |
| 1 | 2501(29.59) | 1404(21.54) | 1192(13.22) | 801(11.11) | 1259(14.26) | 1033(12.82) | 8190(17.04) |
| 2 | 571(6.76) | 2485(38.13) | 628(6.97) | 1407(19.51) | 2365(26.80) | 44(0.55) | 7500(15.60) |
| 3 | 283(3.35) | 94(1.44) | 137(1.52) | 1584(21.97) | 148(1.68) | 2474(30.70) | 4720(9.82) |
| 4 | 1257(14.87) | 531(8.15) | 1134(12.58) | 421(5.84) | 555(6.29) | 490(6.08) | 4388(9.13) |
| 5 | 1281(15.16) | 450(6.91) | 673(7.47) | 433(6.01) | 420(4.76) | 630(7.82) | 3887(8.08) |
| 6 | 626(7.41) | 494(7.58) | 1373(15.23) | 366(5.08) | 145(1.64) | 360(4.47) | 3364(7.00) |
| 7 | 344(4.07) | 177(2.72) | 641(7.11) | 502(6.96) | 351(3.98) | 391(4.85) | 2406(5.00) |
| 8 | 571(6.75) | 333(5.11) | 391(4.34) | 152(2.11) | 221(2.50) | 392(4.86) | 2060(4.28) |
| 9 | 75(0.89) | 20(0.31) | 212(2.35) | 263(3.65) | 393(4.45) | 1041(12.92) | 2004(4.17) |
| 10 | 31(0.37) | 197(3.02) | 346(3.84) | 66(0.92) | 1231(13.95) | 48(0.60) | 1919(3.99) |
| 11 | 73(0.86) | 30(0.46) | 708(7.85) | 239(3.31) | 547(6.20) | 45(0.56) | 1642(3.42) |
| 12 | 38(0.45) | 21(0.32) | 322(3.57) | 58(0.80) | 669(7.58) | 23(0.29) | 1131(2.35) |
| 13 | 3(0.04) | 14(0.21) | 16(0.18) | 351(4.87) | 147(1.67) | 531(6.58) | 1062(2.21) |
| 14 | 159(1.88) | 26(0.40) | 616(6.83) | 9(0.12) | 17(0.19) | 0(0.00) | 827(1.72) |
| 15 | 382(4.52) | 41(0.63) | 146(1.62) | 68(0.94) | 54(0.61) | 5(0.06) | 696(1.45) |
| 16 | 84(0.99) | 17(0.26) | 33(0.37) | 212(2.94) | 25(0.28) | 290(3.60) | 661(1.37) |
| 17 | 100(1.18) | 109(1.68) | 61(0.68) | 85(1.18) | 123(1.38) | 129(1.60) | 607(1.26) |
| 18 | 21(0.25) | 42(0.64) | 348(3.85) | 18(0.25) | 34(0.39) | 3(0.04) | 466(0.97) |
| 19 | 30(0.35) | 12(0.18) | 23(0.26) | 116(1.61) | 43(0.49) | 42(0.52) | 266(0.55) |
| 20 | 6(0.07) | 9(0.14) | 9(0.10) | 41(0.57) | 45(0.51) | 53(0.66) | 163(0.34) |
| 21 | 16(0.19) | 11(0.17) | 5(0.06) | 18(0.25) | 34(0.39) | 34(0.42) | 118(0.25) |
| All | 8452(100) | 6517(100) | 9014(100) | 7210(100) | 8826(100) | 8058(100) | 48077(100) |
