## Supplementary figures and images for "Analysis of the heterogeneity and complexity of murine extraorbital lacrimal gland via single-cell RNA sequencing"

### Fig. S1

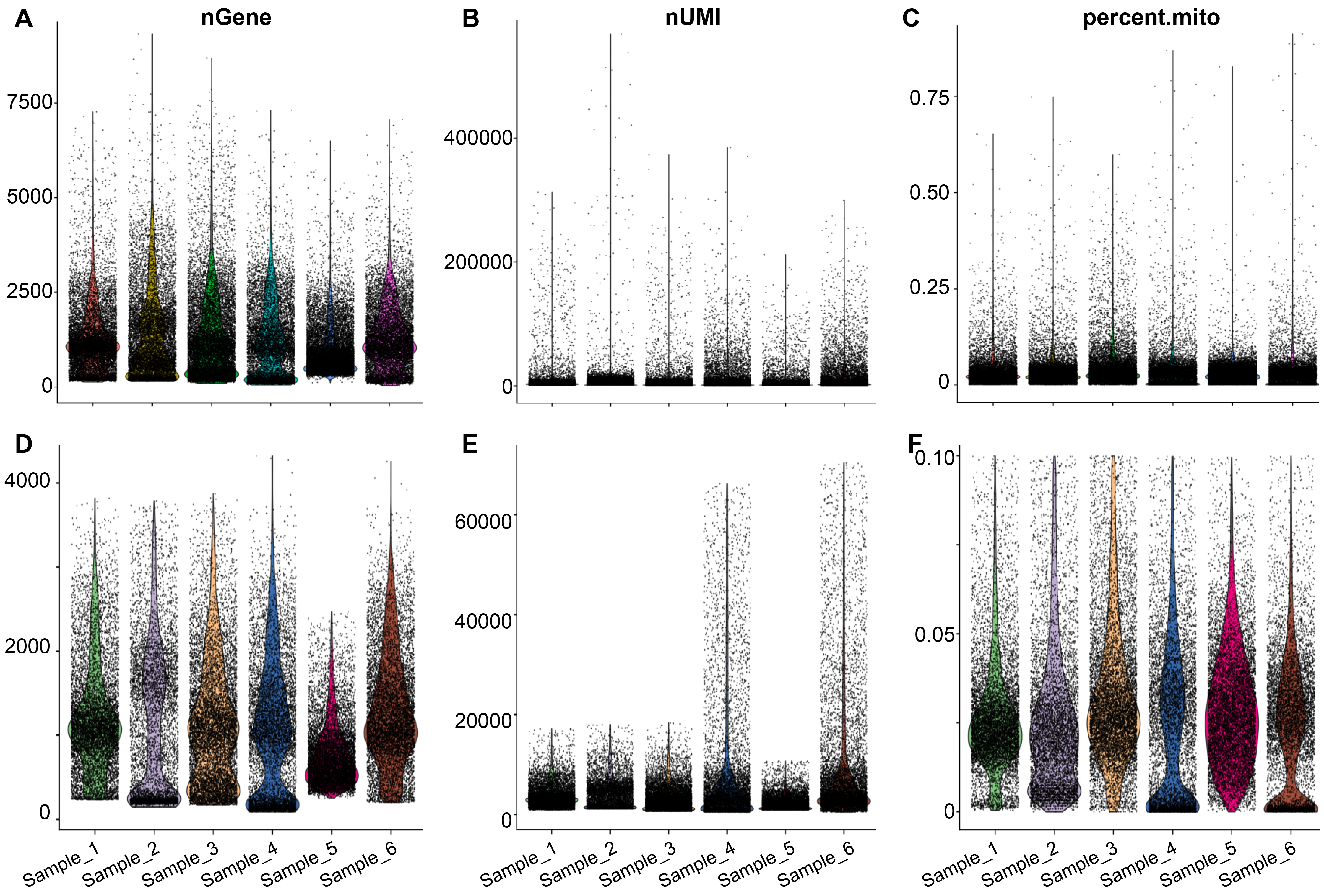

### Fig. S2

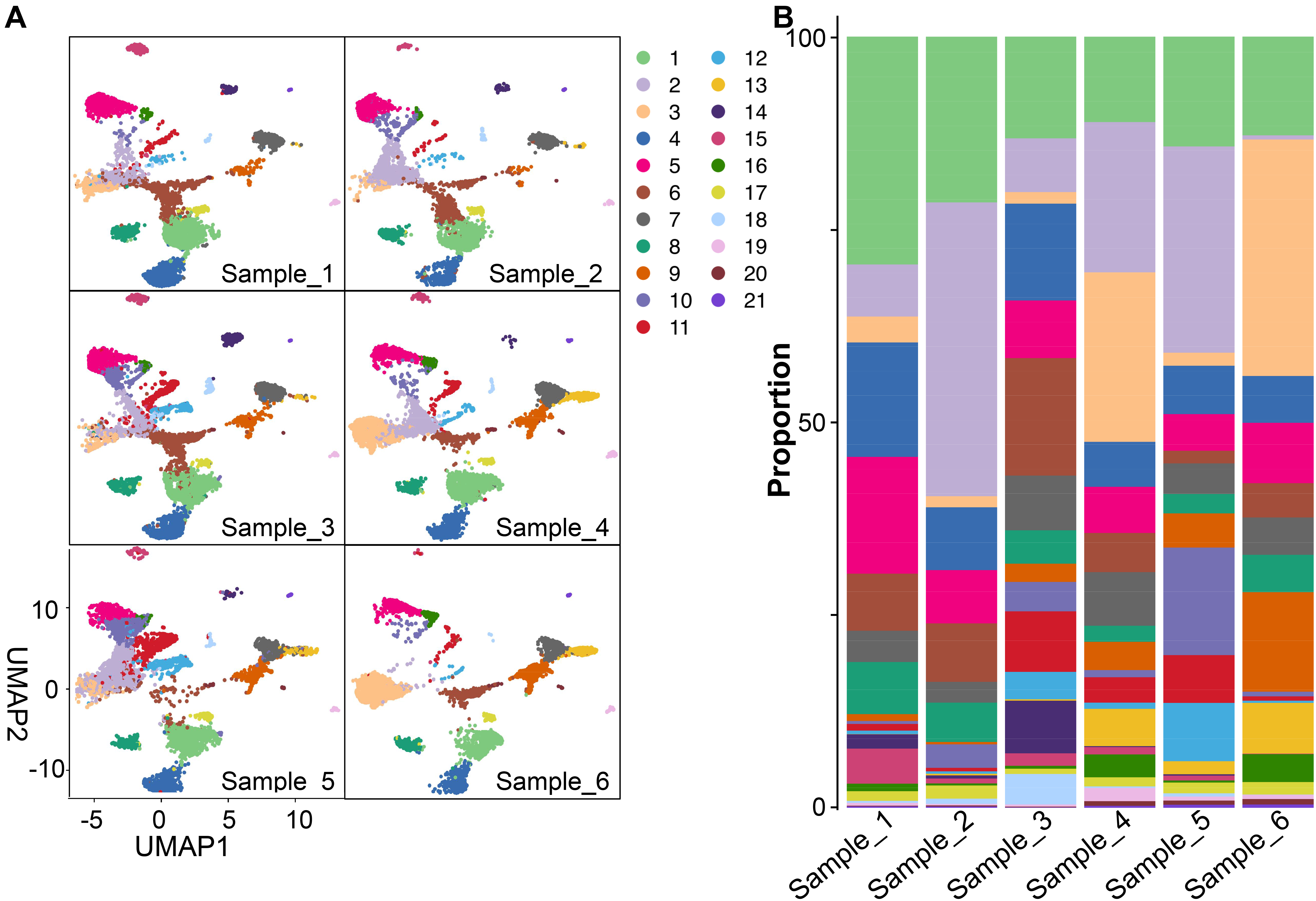

### Fig. S3

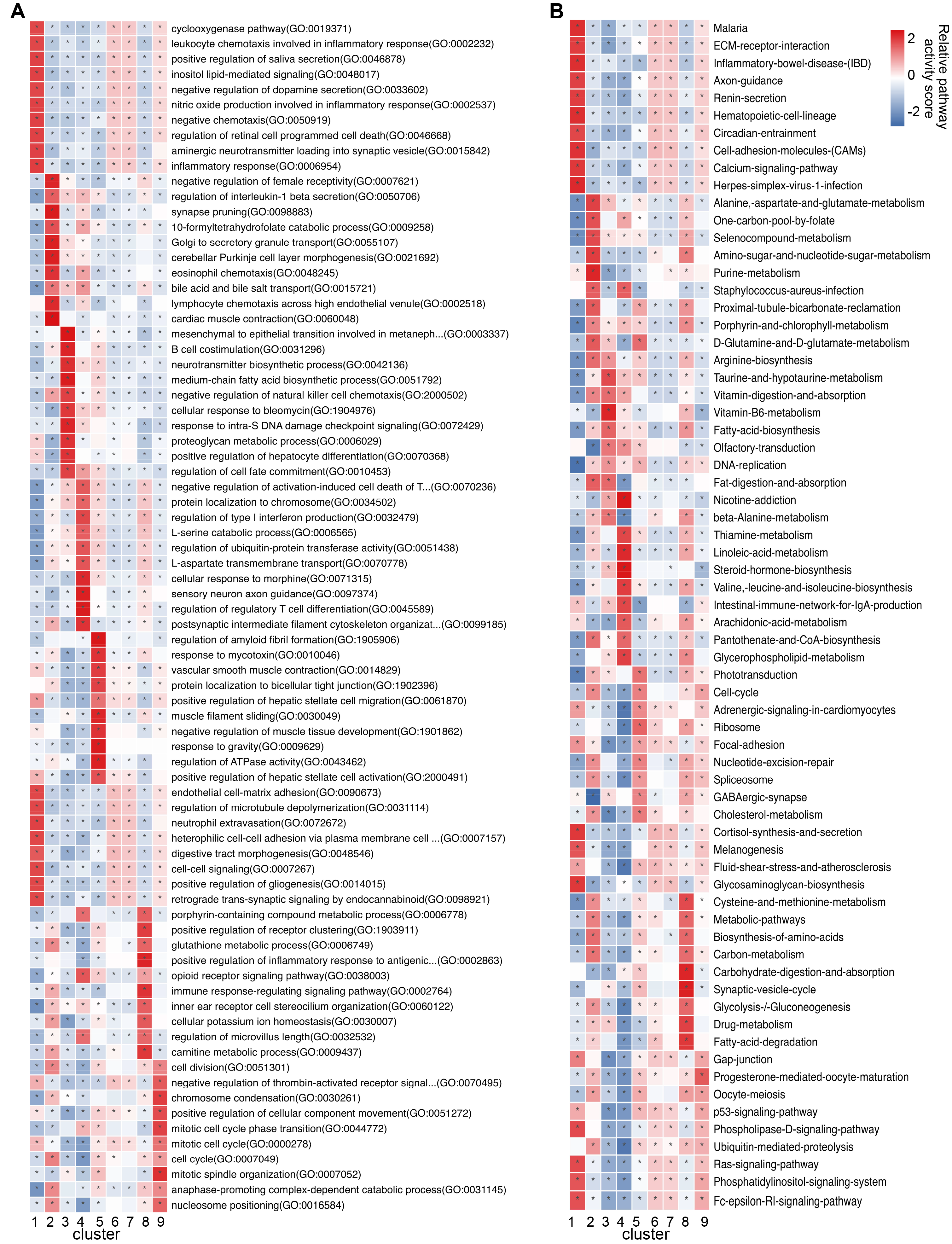

### Fig. S4

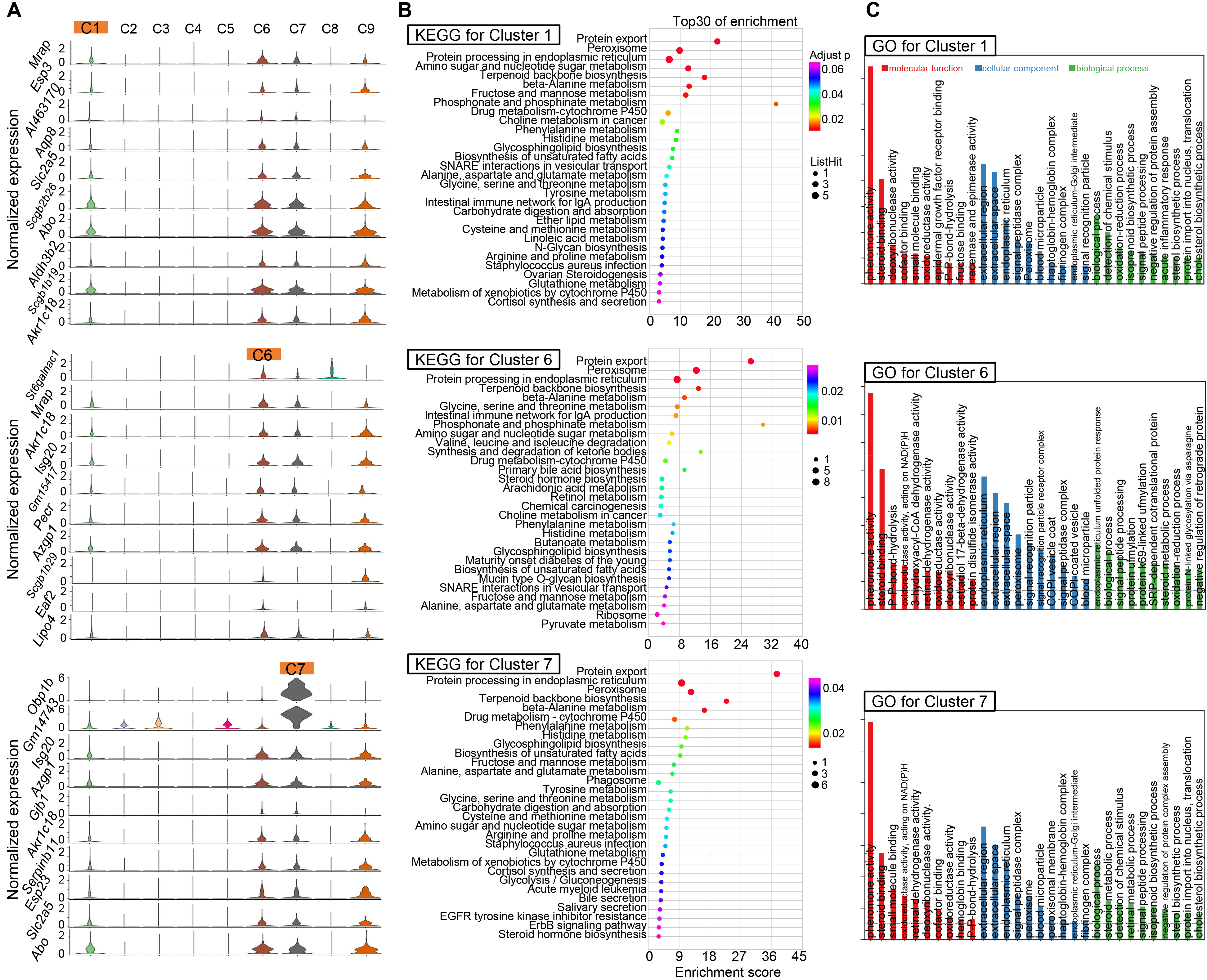

### Fig. S5

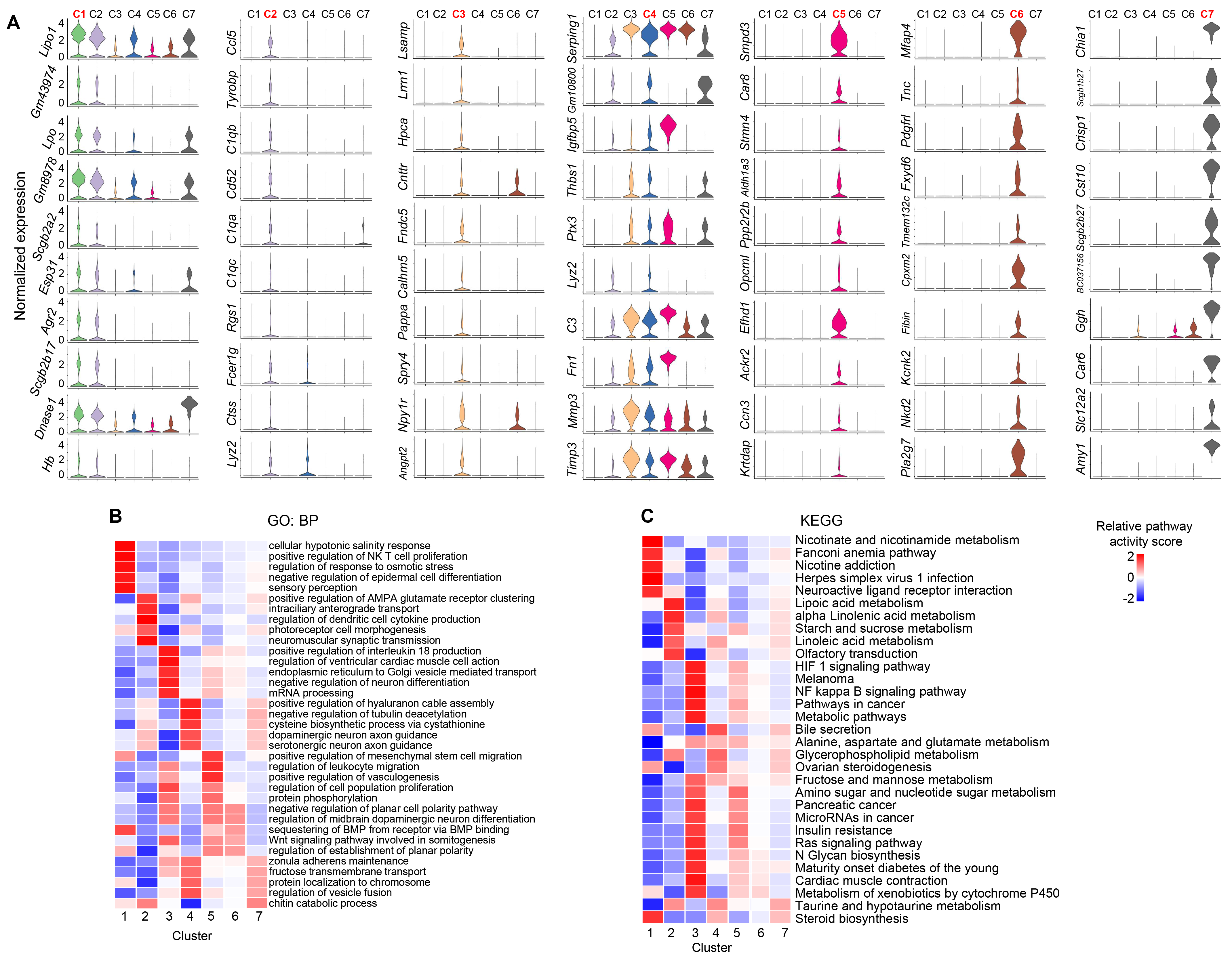

### Fig. S6

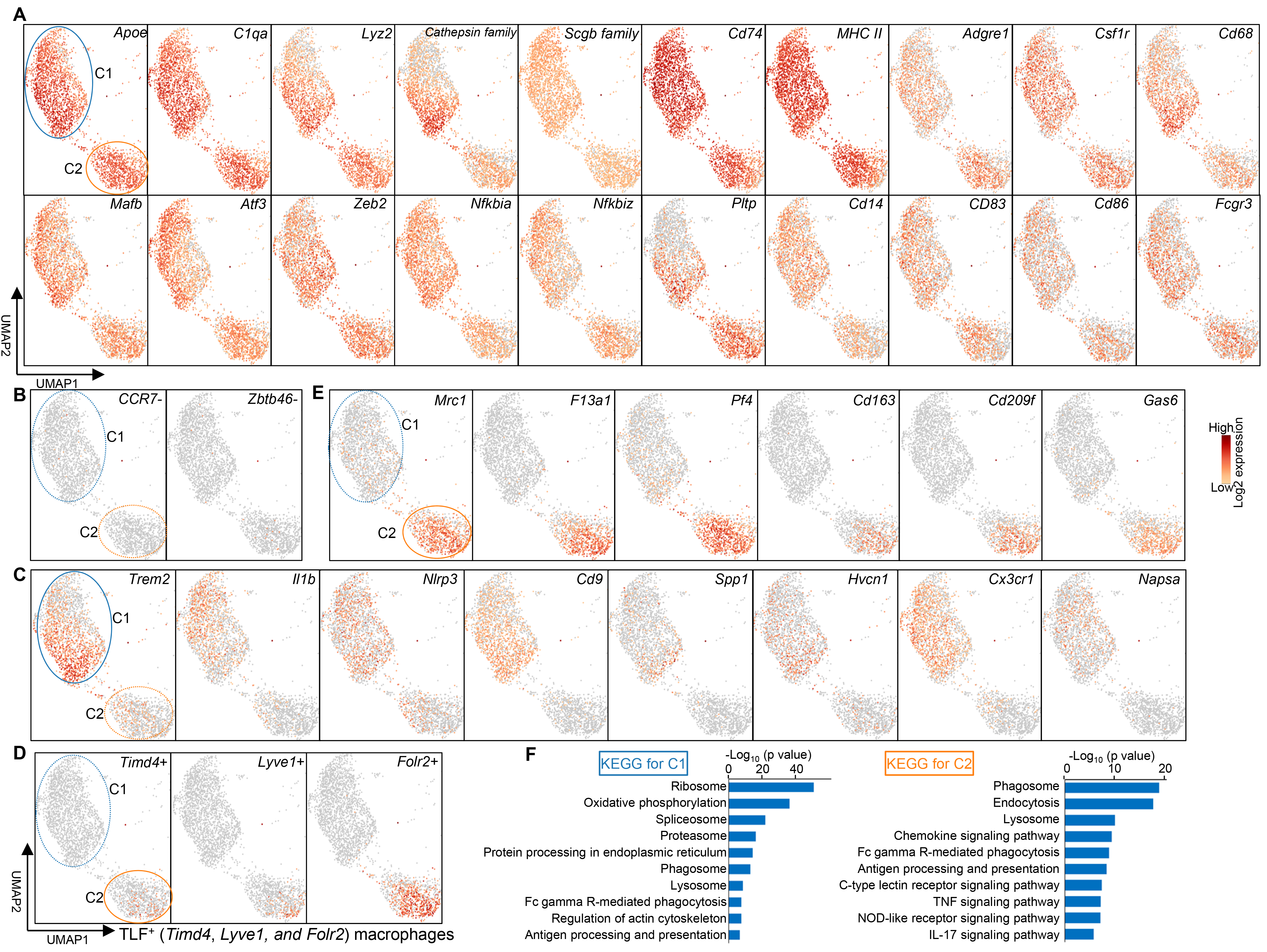

### Fig. S7

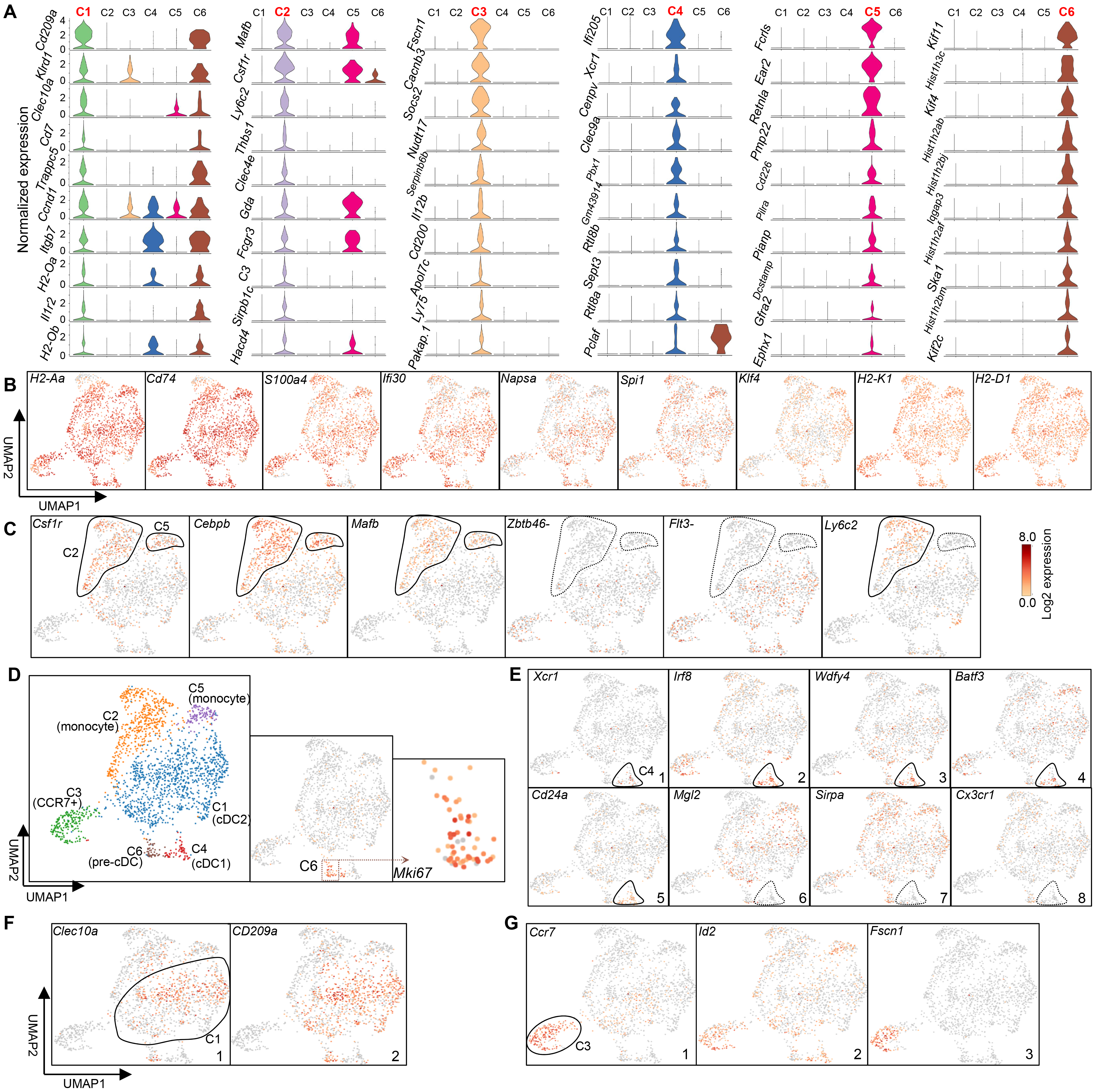
